# RAMPs regulate signalling bias and internalisation of the GIPR

**DOI:** 10.1101/2021.04.08.436756

**Authors:** Matthew Harris, Duncan I. Mackie, John B. Pawlak, Sabrina Carvalho, Tin T. Truong, Dewi Safitri, Ho Yan Yeung, Sarah Routledge, Matthew T. Harper, Bashaier Al-Zaid, Mark Soave, Suleiman Al-Sabah, Asuka Inoue, David R. Poyner, Stephen J. Hill, Stephen J. Briddon, Patrick M. Sexton, Denise Wootten, Peishen Zhao, Kathleen M. Caron, Graham Ladds

**Author notes:** To whom correspondence should be addressed: Dr. Graham Ladds, Department of Pharmacology, University of Cambridge, Tennis Court Rd., Cambridge, CB2 1PD, United Kingdom. Prof. Kathleen Caron, Department of Cell Biology and Physiology, University of North Carolina, Chapel Hill, North Carolina, USA., Dr. Peishen Zhao, Drug Discovery Biology, Monash Institute of Pharmaceutical Sciences, Monash University, Parkville 3052, Victoria, Australia.

## Abstract

Gastric inhibitory polypeptide (GIP) receptor is a class B1 GPCR, that responds to GIP and physiologically potentiates glucose-stimulated insulin secretion. Like most class B1 GPCRs, GIPR has been shown to interact with RAMPs, yet the effects of RAMPs on its signalling and trafficking remain poorly understood. We demonstrate that RAMPs modulate G protein activation and GIPR internalisation profiles. RAMP3 reduced GIPR G_s_ activation and cAMP production but retained GIPR at the cell surface, and this was associated with prolonged ERK1/2 phosphorylation and β-arrestin association. By contrast, RAMP1/2 reduced G_q/11/15_ activation of the GIPR. Through knockout mice studies, we show that RAMP1 is important to the normal physiological functioning of GIPR to regulate blood glucose levels. Thus, RAMPs act on G protein/β-arrestin complexes, having both acute and chronic effects on GIPR function, while this study also raises the possibility of a more general role of RAMP3 to enhance GPCR plasma membrane localisation.

## Introduction

Gastric inhibitory polypeptide (GIP) is a 42 amino acid peptide secreted postprandially from K-enteroendocrine cells^1^. After its release, it is rapidly broken down by dipeptidylpeptidase IV (DPPIV) to GIP (3-42), a weak partial agonist. The substitution of L-Ala for D-Ala to produce GIP (D-Ala2) reduces this breakdown, whilst substitution of the third amino acid, Glu, for Pro in GIP results in the partial agonist (GIP (Pro3))^2^. Together, GIP and glucagon-like peptide 1 (GLP-1) function to potentiate glucose stimulated insulin secretion from pancreatic β-cells^3^, a process severely impaired in type 2 diabetes mellitus (T2DM)^4^. Away from the pancreas, GIP has effects on adipocytes, osteoblasts and neurons, and is thus a potential therapeutic target for diseases such as, T2DM, osteoporosis, Parkinson’s disease and Alzheimer’s disease^5–8^.

GIP (1-42) acts via the GIP receptor (GIPR), a class B1 G protein-coupled receptor (GPCR). Like most class B1 GPCRs, GIPR classically activates Gα_s_ leading to accumulation of cAMP. The notion of pleiotropic signalling - the ability of a receptor to stabilise multiple active conformations to couple to numerous G protein effectors^9^ - has since been demonstrated for most class B1 GPCRs^10^, including the closely related glucagon-like peptide-1 receptor (GLP-1R) and glucagon receptor (GCGR)^11,12^. It has been reported that GIPR activation results in ERK1/2 phosphorylation^13^ and is likely to mobilise intracellular calcium, but additional studies on pleiotropic signalling of the GIPR are lacking.

Further diversity in GPCR signalling can be bought about by interaction with receptor activity-modifying proteins (RAMPs). The three RAMPs (RAMP1, RAMP2 and RAMP3) are single transmembrane spanning proteins initially discovered as molecular chaperones for the calcitonin receptor-like receptor (CLR)^14^. RAMPs and CLR associate in the endoplasmic reticulum and are trafficked to the plasma membrane (PM) since neither can efficiently migrate to the cell surface alone. Beyond their roles as molecular chaperones, RAMPs can modulate ligand binding, G protein coupling, downstream effector recruitment and receptor internalisation and recycling of other class B1 GPCRs including; calcitonin receptor (CTR); parathyroid hormone 1 receptor (PTH1R); parathyroid hormone 2 receptor (PTH2R); secretin receptor (SCTR); GCGR; corticotrophin releasing factor receptor 1 (CRFR1) and; vasoactive intestinal polypeptide receptor 1 (VPAC1R)^15–19^. A recent study demonstrated that the majority of class B1 GPCRs are capable of interacting with RAMPs^20^. However, previous studies demonstrating that the GLP-1R has little, if any, effect on cell surface expression of RAMPs^21,22^, indicate that not all identified interactions are productive for cell surface expressed complexes.

In this study, we report that GIPR indeed signals pleiotropically, activating a wide range of G protein subtypes, promoting cAMP production, mobilising intracellular calcium and ERK1/2 phosphorylation. By utilising flow cytometry and BRET methods for screening GPCR-RAMP interactions, we demonstrate that GIPR interacts with all three RAMPs, and identify multiple other interacting GPCRs that also alter RAMP trafficking. The interaction of RAMPs with GIPR modulates signalling bias of the receptor and shows a clear separation of effects when comparing GIPR complexes with RAMP1 or 2, and RAMP3. This modulation is dependent on a complex interplay between effects on G protein activation and differences in receptor localisation. We also demonstrate that RAMPs play an important role in modulating GIPR activity *in vivo*. Importantly, this study highlights the influence of RAMPs on GIPR pharmacology and demonstrates the need to consider RAMPs when investigating GIPR signalling, as well as other RAMP-interacting GPCRs.

## Results

### BRET and flow cytometry identify GIPR as a novel RAMP interacting GPCR

RAMPs interact with many class B1 GPCRs, although different approaches to measure interaction have yielded differences in their outcomes and are not without controversy^15,20^. We therefore preformed a systematic screen of each RAMP-class B1 GPCR combination using a BRET-based screening assay to identify receptor:RAMP interactions^23^ (Figure 1A-B and Table S1) and a flow cytometry-based assay to verify if these interactions translated to effects on cell surface expression^24^ (Figure 1C-E). In the BRET screen, cells were cotransfected with a constant amount of GPCR-Rluc (GPCR with a C-terminal fusion to Rluc) with increasing amounts of each RAMP-YFP. Based on the screening thresholds established in Mackie et al.^23^ our BRET screen shows that the majority of class B1 GPCRs form either good or strong interactions with all three RAMPs, and corresponds well with a recent suspension bead array-based screen of class B1 GPCR:RAMP interactions^20^. CRFR2 formed only a poor interaction with RAMP1, whilst CRFR1, PAC1R and PTH1R with RAMP1, were the only GPCR: RAMP pairs where an interaction was deemed negative.

**Figure 1.**
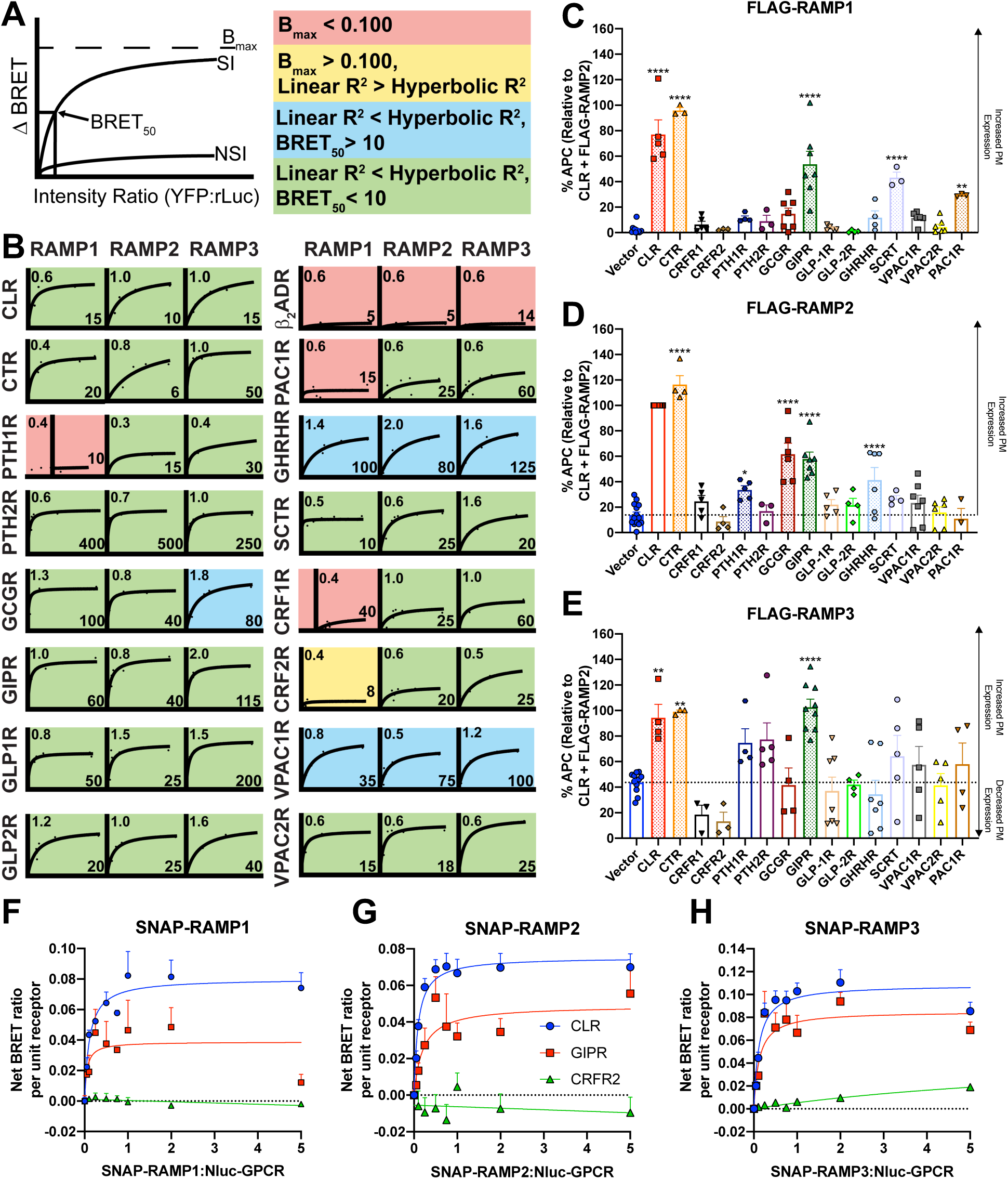
GIPR interacts with all three RAMPs at the plasma membrane. *A-B*. BRET screen for formation of heterodimers between class B1 GPCRs and RAMPs. HEK-293T cells were cotransfected with a constant concentration of GPCR C-terminally tagged with the Rluc-donor moiety and increasing amounts of YFP-acceptor labelled RAMPs. ΔBRET was determined for each receptor:RAMP pair and plotted as a function of the total fluorescence/total luminescence ratio. Data were fitted using one site-binding (hyperbola) and representative saturation isotherms are displayed for each receptor as described in (Mackie et al., 2019)^23^. *A*. A systematic, multi-component approach was used to score the interactions. Firstly, all interactions that failed to reach B_max_ > 0.1 were deemed negative (salmon). Secondly, a comparison of fits between hyperbolic and linear models was used where Linear R^2^ > Hyperbolic R^2^ was deemed a poor interaction (yellow). Finally, the remaining interactions were deemed good (blue) or strong (green) based on the BRET_50_ values: BRET_50_ > 10 (good) or BRET_50_ < 10 (strong). *B*. Representative curves of 3 individual data sets for each RAMP-receptor interaction, with quantitative data reported in Table S1. *C-E*. Flow cytometry analysis of class B1 GPCR-dependent plasma membrane (PM) expression of RAMPs. HEK-293S cells were cotransfected with GPCR and FLAG-RAMP at a 1:1 ratio. PM expression of FLAG-RAMPs was determined by flow cytometry using an APC-conjugated anti-FLAG monoclonal antibody. Surface expression was normalised to FLAG-RAMP2 when cotransfected with CLR as 100%. Endogenous surface expression of FLAG-RAMPs was determined by cotransfection with pcDNA3.1. All values are the mean ± SEM of at least 3 individual data sets. Data were assessed for statistical differences, at *p*<0.05, in cell surface FLAG-RAMP expression compared to expression in the absence of receptor using a one-way ANOVA with Dunnett’s post-hoc test, (*, *p* < 0.05; **, *p* < 0.01; ****, *p* < 0.0001). *F-H*. Cell surface BRET between class B1 GPCRs and RAMPs. HEK-293S cells were cotransfected with a constant concentration of GPCR N-terminally tagged with the Nluc-donor moiety and increasing amounts of SNAP-RAMP. ΔBRET was determined for each receptor:RAMP pair at each ratio. A comparison of linear and hyperbolic fits was performed with the fit with the highest R^2^ value shown (hyperbolic for CLR and GIPR and linear for CRFR2). Data are the mean + SEM of 3-11 individual data sets.

RAMP-interacting GPCRs typically promote plasma membrane localisation of RAMPs^15^. Therefore, we next performed flow cytometry using an APC-conjugated anti-FLAG monoclonal antibody to detect levels of FLAG-RAMP surface expression upon cotransfection with each class B1 receptor to verify if potential protein-protein interactions translated to effects on RAMP surface expression. Little to no cell surface expression of FLAG-RAMP1 or FLAG-RAMP2 was observed when cotransfected with vector control (pcDNA3.1) in HEK-293S cells, although FLAG-RAMP3 partially localised to the plasma membrane in the absence of receptor (Figure 1C-E). This effect has also previously been observed in HEK-293 and Cos-7 cells^23,25^.

Not all RAMP-GPCR interactions identified in the BRET screen translated to effects on RAMP plasma membrane expression. The majority of receptor-RAMP combinations that increased surface expression of FLAG-RAMPs correlated well with previous functional studies^16,21,22,24–27^. We also identify novel RAMP-GPCR interacting partners that promote plasma membrane localisation of RAMPs including the SCTR with RAMP1, GHRHR with RAMP2 and the GIPR with all three RAMPs (Figure 1C-E). Also, of note, whilst there was no BRET interaction between PAC1R and RAMP1 there was a significant elevation in FLAG-RAMP1 surface expression with PAC1R (Figure 1C). Overall, these data exemplify the variety of cellular interactions between RAMPs and GPCRs, demonstrating a necessity for orthogonal approaches to screening.

Coexpression of GIPR with each FLAG-RAMP significantly promoted RAMP surface expression (Figure 1C-E), with the effect on RAMP3 comparable to that of CLR (the archetypal RAMP-interacting receptor). This novel interaction was also verified using ELISA (Figure S1A). Whilst there was no reciprocal effect of RAMP1 or RAMP2 coexpression on the plasma membrane expression of GIPR, RAMP3 co-expression resulted in a small (∼20%), but significant increase in GIPR cell surface expression compared to GIPR alone (Figure S1B).

To interrogate the difference observed between the BRET and flow cytometry screens, we generated SNAP-RAMP and Nluc-CLR/GIPR/CRFR2 fusion constructs and utilised a cell impermeable SNAP reagent (SNAP-Surface® Alexa® Fluor 488) to measure BRET between GPCR and RAMP only at the cell surface. All of the newly generated fusion constructs were shown to be functional (Figure S2). While the rank order of potency for CLR agonists in the presence of SNAP-RAMP3 was identical to FLAG- or HA- tagged RAMPs, there were some discrepancies in the potencies of the non-cognate ligands with the CLR-SNAP-RAMP1/2 complexes. However, SNAP-RAMP constructs were used solely for the purpose of a BRET assay to assess direct receptor-RAMP interactions at the cell surface, to support data achieved using the ELISA and flow cytometry methods reporting receptor and RAMP expression. As predicted, CLR and GIPR exhibited saturable increases in ΔBRET with increasing concentration of all three SNAP-RAMPs, indicating direct interactions (Figure 1F-H), whilst there was only a very small, linear increase in ΔBRET between CRFR2 and RAMP3, and no effect with RAMP1 or RAMP2. Cell surface localised BRET between RAMPs and CLR/GIPR was confirmed through cell surface BRET imaging (Figure S1C-D). The two BRET assays, together with the lower cell surface expression of FLAG-RAMP3 in the presence of CRFR2, indicate that CRFR2 may interact with RAMP2, but that this complex does not traffic to the cell surface, an effect previously observed with the atypical chemokine receptor 3 (ACKR3)^23^. Thus, many of the potential GPCR:RAMP complexes identified in the initial BRET screen may only occur intracellularly. Overall, these data provide evidence for an interaction between GIPR and all three RAMPs and that these complexes are present at the cell surface.

### Signalling pleiotropy of the GIPR

GIPR is traditionally considered to be a Gα_s_-coupled GPCR whereby activation leads to production of cAMP^1^. However, many class B1 GPCRs, including the closely related glucagon and GLP-1 receptors are known to signal pleiotropically^10^. GCGR and GLP-1R stimulate release of calcium from intracellular stores ([Ca^2+^]_i_), and ERK1/2 phosphorylation; events that are reported to be downstream of G_s_, G_q_, G_i_ and/or β-arrestins^11,12,28,29^. Therefore, we firstly sought to characterise the ability of the GIPR to pleiotropically couple to different transducers, namely distinct G protein subtypes and β-arrestins. We utilised a NanoBiT system to evaluate agonist-dependent dissociation of each individual Gα-LgBiT from Gβγ_2_-SmBiT^29,30,31^, along with BRET to assess recruitment of β-arrestin1/2-YFP to myc-GIPR-Rluc (functionally validated in Figure S2B) as measures of G protein activation and β-arrestin1/2 recruitment at the GIPR (Figure 2A, Figure S3). The GIPR coupled to multiple distinct G protein subfamilies and recruited β-arrestins when stimulated with its cognate ligand GIP (1-42) in HEK-293 cells transiently expressing the GIPR. A rank order of potency was established for activation of each Gα in combination with the optimal Gβγ_2_ complex (that which resulted in the largest range and most robust response^32^, Figure S4), and, recruitment of β-arrestin1/2: G_s_ ≈ G_12_ > G_i2_ > G_q_ ≈ G_13_ > G_z_ > β-Arr2 > G_i3_ ≈ β-Arr1 ≈ G_11_> G_15_ > G_14_ (Figure 2A, Figure S3). These experiments demonstrate that GIPR couples to more than one G protein, albeit with significantly lower potency relative to G_s_ for all effectors except G_12_, G_i2_ and G_q_ (Figure 2A). Whilst most G proteins fit to a monophasic curve, both G_z_ and G_12_ displayed two components to the response: a high potency first component (pEC_50_ of 8.77±0.55 and 11.00±0.22, respectively) and a lower potency second component (pEC_50_ of 6.18±0.61 and 8.18±0.24, respectively). To further assess signalling, the ability of GIP (1-42) to activate three intracellular signalling pathways (cAMP accumulation, [Ca^2+^]_i_ mobilisation and phosphorylation of ERK1/2) was assessed (Figure 2B-D, Table 1). In all cases, GIP (1-42) was able to stimulate concentration-dependent responses. Both cAMP and [Ca^2+^]_i_ mobilisation were monophasic, whilst ERK1/2 phosphorylation displayed a pronounced biphasic response. Unsurprisingly, the cAMP response was the most potent, with lower potency observed for [Ca^2+^]_i_ mobilisation and ERK1/2 phosphorylation (cAMP: pEC_50_ of 9.83± 0.08; [Ca^2+^]_i_: pEC_50_ of 8.70±0.16; pERK1/2: three-parameter fit pEC_50_ of 7.93±0.2; pEC_50_1_ of 9.13±0.61, pEC_50_2_ of 6.22±0.50 as determined using the biphasic model).

**Table 1:** Potency (pEC_50_) and E_max_values for cAMP production, intracellular calcium mobilisation and ERK1/2 phosphorylation at the GIPR, stimulated with various GIP-based and glucagon family ligands measured in HEK 293S cells.

|  |  | Relative to system E<sub>max</sub><sup>a</sup> | Relative to GIP (1-42) E<sub>max</sub><sup>c</sup> |  |  |  |  |  |
| --- | --- | --- | --- | --- | --- | --- | --- | --- |
|  |  | GIP (1-42) | GIP (D-Ala2) | GIP (Pro3) | GIP (3-42) | Glucagon | GLP-1 (7-36)NH<sub>2</sub> | Oxyntomodulin |
| cAMP | pEC<sub>50</sub><sup>b</sup> | 9.83±0.08 | 10.06±0.10 | 7.64±0.11 | 8.07±0.48 | 6.97±0.27 | 6.68±0.33 | 6.77±0.88 |
|  | E<sub>max</sub> | 70.6±2.2 | 106.6±4.0 | 68.0±2.5 | 23.0±3.2 | 26.8±2.9 | 37.2±4.1 | 14.4±3.80 |
|  | n | 7 | 7 | 5 | 10 | 8 | 8 | 7 |
| [Ca<sup>2+</sup>]<sub>i</sub> | pEC<sub>50</sub><sup>b</sup> | 8.70±0.16 | 8.93±0.28 | 7.84±0.23 | ND | ND | ND | 6.81±0.62 |
|  | E<sub>max</sub> | 23.1±1.0 | 91.6±7.6 | 23.9±2.8 | ND | ND | ND | 16.3±5.9 |
|  | n | 4 | 3 | 3 | 3 | 3 | 3 | 3 |
| pERK1/2 | pEC<sub>50</sub><sup>b</sup> | 7.93±0.26 | 7.81±0.20 | 7.16±0.27 | 6.61±0.90 | 5.92±0.34 | ND | 5.32±0.59 |
|  | E<sub>max</sub> | 31.1±2.5 | 91.8±6.1 | 79.8±9.5 | 31.3±13.2 | 62.9±17.2 | ND | 90.5±46.2 |
|  | pEC<sub>50_1</sub><sup>d</sup> | 9.13±0.61 | 8.72±0.60 | 7.54±0.49 | ND | ND | ND | ND |
|  | pEC<sub>50_2</sub><sup>e</sup> | 6.22±0.50 | 6.88±0.46 | 5.21±0.46 | ND | ND | ND | ND |
|  | Frac<sup>f</sup> | 40.6±25 | 45.3±22 | 45.6±14 | ND | ND | ND | ND |
|  | n | 10 | 5 | 4 | 3 | 3 | 3 | 3 |

Data are the mean ± SEM of *n* individual data sets.
<sup>a</sup> The maximal response to the ligand expressed as a percentage of the maximal response of the system for each pathway; 100 μM forskolin for cAMP, 10 μM ionomycin for [Ca<sup>2+</sup>]<sub>i</sub> and 100 μM PMA for pERK1/2.
<sup>b</sup> The maximal response to the ligand expressed as a percentage of the GIP (1-42) response for each pathway
<sup>c</sup> The negative logarithm of the agonist concentration required to produce a half-maximal response, determined by fitting to the three-parameter Log [Agonist] vs response model for each individual experiment.
<sup>d</sup> The negative logarithm of the agonist concentration required to produce a 50% of the first component of the response, determined by fitting to the biphasic Log [Agonist] vs response model
<sup>e</sup> The negative logarithm of the agonist concentration required to produce a 50% of the second component of the response, determined by fitting to the biphasic Log [Agonist] vs response model
<sup>f</sup> The fraction of the concentration-response curve derived from the first, high potency component
ND = Not defined as data could not be fitted using the appropriate model

**Figure 2.**
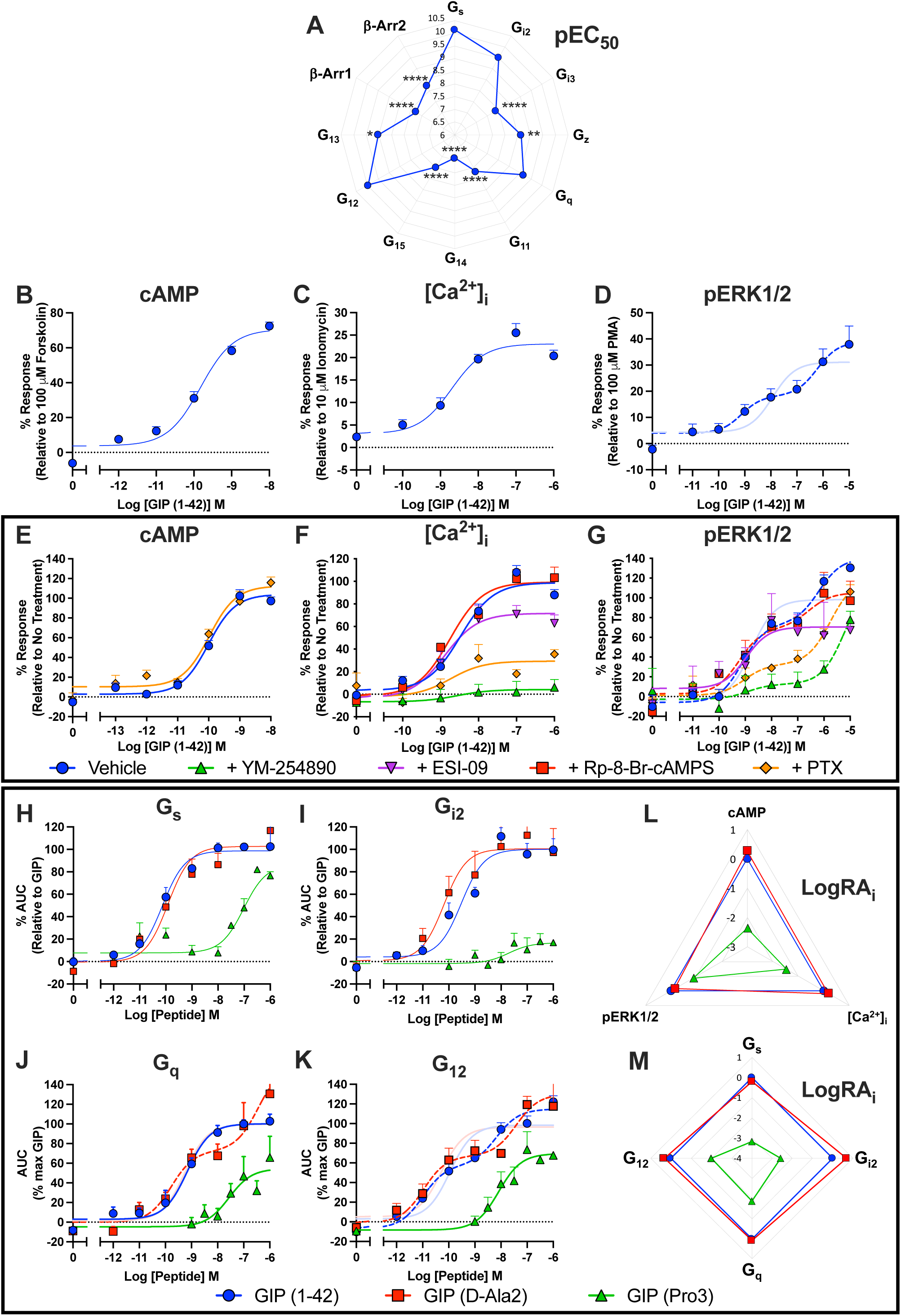
GIPR activation results in promiscuous G protein activation, recruitment of β-arrestins and stimulation of cAMP accumulation, intracellular calcium mobilisation ([Ca^2+^]_i_) and ERK1/2 phosphorylation. *A*. Average pEC_50_ values, obtained from three-parameter logistic fits, for activation of each G protein and recruitment of β-arrestin measured by loss of luminescence upon Gα-LgBiT and Gβγ_2_-SmBiT dissociation and BRET between GIPR-Rluc and β-arrestin1/2-YFP, respectively. Data were assessed for statistical differences, at *p*<0.05, in pEC_50_ compared to activation of G_s_ using a one-way ANOVA with Dunnett’s post-hoc test (*, *p*<0.05; **, *p*<0.01; ****, *p*<0.0001). *B*-*D*. HEK-293S cells were transiently transfected with untagged GIPR. cAMP accumulation was measured following 8 min stimulation with GIP (1-42), and data are expressed relative to 100 μM forskolin (*B*). For [Ca^2+^]_i_ mobilisation (*C*), cells were stimulated for 2 min with GIP (1-42) with the intensity at the time of peak response used to construct the concentration-response curves. Data are expressed relative to 10 μM ionomycin. Phosphorylation of ERK1/2 was determined after 5 min stimulation with GIP (1-42) and data are expressed relative to 100 μM PMA (*D*). The dashed line represents the biphasic fit, with the equivalent three-parameter fit faded with quantitative data displayed in (Table 1). *E-G*. To determine G protein and signalling pathway contribution to cAMP accumulation (*E*), [Ca^2+^]_i_ mobilisation (*F*) and ERK1/2 phosphorylation (*G*), cells were pretreated with or without pertussis toxin (PTx, to inhibit Gα_i/o_), YM-254890 (to inhibit Gα_q/11/14_), Rp-8-Br-cAMPS (to inhibit PKA) or ESI-09 (to inhibit EPAC1/2) and cAMP accumulation, [Ca^2+^]_i_ mobilisation and ERK1/2 phosphorylation measured as described in Figure 2. Dashed lines represent biphasic fits, with the equivalent three-parameter fit for vehicle faded. Data are normalised to the maximal response in the absence of treatment, determined by fitting to the three-parameter logistic model and are mean + SEM of 3-7 individual experiments. All data for *B-G* are expressed as mean + SEM. *H-K*. HEK-293A cells were cotransfected with GIPR, one of Gα_s_-LgBiT (*H*), Gα_i2_-LgBiT (*I*), Gα_q_-LgBiT (*J*) or Gα_12_-LgBiT (*K*), Gβ_1_, Gγ_2_-SmBiT, and pcDNA3.1 at a 2:1:3:3:2 ratio. RIC8A was also included for G_q_. G protein activation was measured by the agonist-induced change in relative luminescence units (RLU). Data points were corrected to baseline and vehicle and the AUC used to produce concentration-response curves for each G protein, expressed relative to the maximal response to GIP (1-42), determined by three-parameter logistic fits. Data are expressed as mean + SEM of 4-7 individual experiments with quantitative data displayed in (Table S2). Biphasic fits for GIP (1-42) (G_12_) and GIP (D-Ala2) (G_q_ and G_12_) are displayed as dashed lines, with the equivalent three parameter fits faded. *L-M*. Radial plots demonstrating the intrinsic relative activity (LogRA_i_) on a linear scale for GIP (D-Ala2) and GIP (Pro3)-mediated cAMP accumulation, [Ca^2+^]_i_ mobilisation and ERK1/2 phosphorylation (*L*) or activation of G_s_, G_q_, G_i2_ and G_12_ (*M*), relative to GIP (1-42).

To determine the contribution of individual G protein subfamilies and downstream effectors to modulation of these three GIPR-mediated signalling pathways, we stimulated the GIPR, transiently expressed in HEK-293S cells, after pre-treatment with a number of pharmacological inhibitors (Figure 2E-G). In this system, cAMP accumulation was independent of Gα_i/o_ coupling as overnight pre-treatment with 200 ng/ml pertussis toxin (PTx) did not alter cAMP responses (Figure 2E). Given the biphasic nature of the pERK1/2 response, it was not surprising that multiple inhibitors modulated the GIP (1-42) response (Figure 2G). 30-minute pre-treatment with 100 nM YM-254890, to selectively inhibit G_q/11/14_ activation, or pre-treatment with PTx significantly attenuated the high potency first component of ERK1/2 phosphorylation (*p*=0.0027 and *p*=0.0465, respectively). The lower potency, second pERK1/2 component appeared to be EPAC1/2-dependent as pre-treatment with non-selective EPAC1/2 inhibitor, ESI-09, removed any low affinity response. Despite being monophasic, the [Ca^2+^]_i_ response was also affected by these same effectors (Figure 2F). Pre-treatment with YM-254890, or PTx significantly reduced [Ca^2+^]_i_ mobilisation, by approximately 90% (*p*<0.0001) and 70% (*p*<0.0001), respectively. There was also a small EPAC1/2-dependent component to [Ca^2+^]_i_ mobilisation (*p*=0.0021). This indicates that GIPR stimulates [Ca^2+^]_i_ mobilisation and promotes phosphorylation of ERK1/2 via at least Gα_q/11/14_, Gα_i/o_ and EPAC1/2-dependent mechanisms.

GCGR and GLP-1R peptide agonists exhibit cross reactivity with related receptors and have potential for differences in biased agonism^22,33,34^. Consequently, we investigated a series of GIP peptide analogues and GCGR and GLP-1R agonists for interaction with GIPR and potential biased agonism in second messenger assays (Figure S5, Table 1). In all cases, GIP (D-Ala2) had almost identical potency and E_max_ to GIP (1-42), GIP (Pro3) was a partial agonist with lower potency and, GIP (3-42) had a substantially lower maximum response with similar potency to GIP (Pro3). For ERK1/2 phosphorylation, GIP (D-Ala2) and GIP (Pro3) responses were biphasic (pEC_50_1_ of 8.72±0.60 and 7.49±0.12 and pEC_50_2_ of 6.88±0.46 and 4.50±0.41, respectively). Furthermore, all GCGR and GLP-1R family ligands were monophasic and very weak partial agonists with potency values in the high nM to low μM range. Overall, these findings illustrate that GIPR signals pleiotropically, has very little cross-reactivity with glucagon or GLP-1 family ligands in common second messenger assays and only the three most potent agonist for ERK1/2 phosphorylation were able to generate a biphasic response. Assessment of G protein activation for the two most potent GIP-peptide analogues after GIP (1-42); GIP (D-Ala2) and GIP (Pro3) was also performed using representatives for each G protein subtype (Figure 2H-K, Figure S6, Table S2), revealing a similar rank order of potency to that observed for the intracellular signalling pathways. At the G protein level, within the concentration ranges assessed, there was no evidence of biphasic responses for GIP (Pro3) at any of the four G proteins, whilst, similar to GIP (1-42), GIP (D-Ala2) was biphasic for G_12_ (pEC_50_1_ of 10.97±0.30 and pEC_50_2_ of 7.42±0.46). Interestingly, GIP (D-Ala2) also displayed two phases of response at G_q_ (pEC_50_1_ of 9.77±0.43 and pEC_50_2_ of 6.45±0.40), unlike GIP (1-42). Calculation of the Log relative intrinsic activity (LogRA_i_, a measure of the E_max_ and EC_50_ ratio for test and reference) for GIP (D-Ala2), relative to GIP (1-42), at each signalling pathway and G protein tested revealed no substantial biased agonism (Figure 2L and M). However, GIP (Pro3) exhibited biased agonism, relative to GIP (1-42), with significant bias towards ERK phosphorylation (*p*=0.005) relative to cAMP, with G_q_ also trending towards bias relative to G_s_ (*p*=0.09).

### RAMPs differentially modulate GIPR signalling

After establishing that GIPR interacts with, and promotes, RAMP surface expression, we set out to determine whether RAMPs modulate GIPR signalling. For this, GIPR was co-expressed with each FLAG-RAMP and the ability of GIP (1-42), GIP (D-Ala2) and GIP (Pro3) to stimulate cAMP accumulation, [Ca^2+^]_i_ mobilisation and ERK1/2 phosphorylation was assayed under the same experimental conditions as Figure 2 (Figure 3A-C Table 2). Despite co-expression of RAMP3 resulting in increased cell surface expression of GIPR, RAMP3 led to a small, but significant, reduction in the potency and E_max_ for cAMP accumulation for all three agonists (Figure 3A). Co-expression of GIPR with RAMP1 or RAMP2 did not alter GIP-mediated cAMP accumulation. In contrast, co-expression with RAMP1 or RAMP2 significantly reduce the E_max_ for [Ca^2+^]_i_ mobilisation for all 3 peptides (Figure 3B, Table 2). A similar trend was observed for ERK1/2 phosphorylation, with RAMP1 or RAMP2 co-expression reducing E_max_ for all three agonists when a three-parameter curve fit was applied to the data (Figure 3C, Table 2). Analysis using the biphasic model, which better describes the data, revealed that the attenuated pERK1/2 response upon coexpression with RAMP1 and RAMP2 was a result of significant attenuation to the fraction of the response mediated by the high potency component for GIP (1-42) (*p*=0.012 and *p*=0.0038). Similarly, RAMP2 significantly reduced the fraction of the response mediated by the high potency component for GIP (D-Ala2) (*p*=0.028), whilst the effect with RAMP1 trended towards significance (*p*=0.11). RAMP3 had no effect on [Ca^2+^]_i_ mobilisation or ERK1/2 phosphorylation.

**Table 2:** Potency (pEC_50_) and E_max_ values for cAMP production, intracellular calcium mobilisation ((Ca^2+^)_i_) and ERK1/2 phosphorylation in HEK 293S cells expressing the GIPR and either pcDNA3.1, FLAG-RAMP1, FLAG-RAMP2 or FLAG-RAMP3 in response to GIP (1-42), GIP (D-Ala2) or GIP (Pro3).

|  |  | GIPR |  |  | + RAMP1 |  |  |
| --- | --- | --- | --- | --- | --- | --- | --- |
|  |  | GIP (1-42) | GIP (D-Ala2) | GIP (Pro3) | GIP (1-42) | GIP (D-Ala2) | GIP (Pro3) |
| cAMP | pEC<sub>50</sub><sup>a</sup> | 10.24±0.10 | 10.27±0.07 | 7.69±0.07 | 10.12±0.12 | 10.13±0.09 | 7.58±0.15 |
|  | E<sub>max</sub><sup>b</sup> | 99.9±3.5 | 100.2±2.7 | 100.1±2.6 | 105.8±5.3 | 103.1±3.6 | 91.3±5.1 |
|  | n | 4 | 4 | 4 | 4 | 4 | 4 |
| [Ca<sup>2+</sup>]<sub>i</sub> | pEC<sub>50</sub><sup>a</sup> | 8.81±0.13 | 9.01±0.09 | 7.90±0.12 | 8.69±0.14 | 8.82±0.20 | 7.80±0.38 |
|  | E<sub>max</sub><sup>b</sup> | 99.6±4.4 | 99.8±3.0 | 98.4±5.0 | 68.6±3.4*** | 46.5±3.5**** | 19.6±3.8**** |
|  | n | 3 | 3 | 3 | 3 | 3 | 3 |
| pERK1/2 | pEC<sub>50</sub><sup>a</sup> | 8.47±0.26 | 8.09±0.35 | 5.77±0.41 | 8.07±0.37 | 7.93±0.43 | 6.59±0.80 |
|  | E<sub>max</sub><sup>b</sup> | 100.1±7.7 | 105.3±12.3 | 121.9±34.9 | 56.4±7.9** | 62.6±9.7* | 65.9±26.7 |
|  | pEC<sub>50_1</sub><sup>c</sup> | 8.59±0.27 | 8.51±0.57 | ND | 8.15±0.47 | 8.36±0.68 | ND |
|  | pEC<sub>50_2</sub><sup>d</sup> | 4.70±0.63 | 5.38±0.63 | ND | 4.59±0.33 | 4.93±0.29 | ND |
|  | Frac<sup>e</sup> | 66.8±7 | 54.9±15 | ND | 34.0±7* | 31.2±9 | ND |
|  | n | 3 | 3 | 3 | 3 | 3 | 3 |
|  |  | GIPR |  |  | + RAMP2 |  |  |
|  |  | GIP (1-42) | GIP (D-Ala2) | GIP (Pro3) | GIP (1-42) | GIP (D-Ala2) | GIP (Pro3) |
| cAMP | pEC<sub>50</sub><sup>a</sup> | 10.27±0.08 | 10.27±0.06 | 7.70±0.06 | 10.09±0.09 | 10.16±0.07 | 7.45±0.11 |
|  | E<sub>max</sub><sup>b</sup> | 99.9±2.9 | 100.1±2.3 | 100.0±2.2 | 106.7±4.0 | 101.2±3.0 | 103.3±4.3 |
|  | n | 5 | 5 | 5 | 5 | 5 | 5 |
| [Ca<sup>2+</sup>]<sub>i</sub> | pEC<sub>50</sub><sup>a</sup> | 8.78±0.08 | 8.50±0.095 | 7.85±0.14 | 8.61±0.14 | 8.60±0.14 | 7.65±0.34 |
|  | E<sub>max</sub><sup>b</sup> | 100.2±2.7 | 99.9±3.7 | 99.3±6.2 | 68.4±3.1**** | 65.2±3.3**** | 50.3±6.8*** |
|  | n | 3 | 3 | 3 | 3 | 3 | 3 |
| pERK1/2 | pEC<sub>50</sub><sup>a</sup> | 8.14±0.19 | 8.06±0.24 | 5.89±0.28 | 8.94±0.70 | 7.42±0.47 | 6.05±0.49 |
|  | E<sub>max</sub><sup>b</sup> | 98.3±5.9 | 101.4±8.0 | 122.3±21.9 | 39.5±8.6*** | 41.7±9.5*** | 66.4±19.9 |
|  | pEC<sub>50_1</sub><sup>c</sup> | 8.38±0.21 | 8.36±0.34 | ND | 9.09±0.83 | 7.87±0.71 | ND |
|  | pEC<sub>50_2</sub><sup>d</sup> | 4.98±0.33 | 5.24±0.43 | ND | 4.42±0.47 | 4.58±0.32 | ND |
|  | Frac<sup>e</sup> | 61.0±6 | 58.0±9 | ND | 23.4±7** | 25.4±9* | ND |
|  | n | 3 | 3 | 3 | 3 | 3 | 3 |
|  |  | GIPR |  |  | + RAMP3 |  |  |
|  |  | GIP (1-42) | GIP (D-Ala2) | GIP (Pro3) | GIP (1-42) | GIP (D-Ala2) | GIP (Pro3) |
| cAMP | pEC<sub>50</sub><sup>a</sup> | 10.00±0.06 | 9.95±0.05 | 7.59±0.04 | 9.58±0.07*** | 9.57±0.07*** | 7.08±0.09**** |
|  | E<sub>max</sub><sup>b</sup> | 100.1±2.4 | 98.3±2.2 | 99.2±1.5 | 91.8±2.7* | 87.5±2.9** | 75.7±2.9**** |
|  | n | 7 | 8 | 14 | 8 | 8 | 13 |
| [Ca<sup>2+</sup>]<sub>i</sub> | pEC<sub>50</sub><sup>a</sup> | 8.60±0.09 | 8.88±0.088 | 8.07±0.17 | 8.59±0.12 | 8.92±0.087 | 7.70±0.19 |
|  | E<sub>max</sub><sup>b</sup> | 98.2±2.9 | 99.6±3.0 | 98.9±6.6 | 93.3±3.7 | 92.7±2.6 | 89.3±6.3 |

|  | n | 6 | 3 | 3 | 6 | 3 | 3 |
| --- | --- | --- | --- | --- | --- | --- | --- |
| pERK1/2 | pEC<sub>50</sub><sup>a</sup> | 8.18±0.14 | 7.77±0.18 | 6.87±0.19 | 8.03±0.17 | 7.89±0.19 | 6.41±0.29 |
|  | E<sub>max</sub><sup>b</sup> | 98.6±4.6 | 92.1±5.8 | 85.8±8.0 | 106.1±6.0 | 96.5±6.0 | 104.0±14.3 |
|  | pEC<sub>50_1</sub><sup>c</sup> | 8.54±0.17 | 8.40±0.28 | 7.70±0.44 | 8.94±0.24 | 8.96±0.26 | 8.51±0.89 |
|  | pEC<sub>50_2</sub><sup>d</sup> | 5.45±0.17 | 5.43±0.23 | 5.52±0.32 | 5.98±0.21 | 5.90±0.18 | 5.91±0.27 |
|  | Frac<sup>e</sup> | 56.1±5 | 45.8±6 | 37.6±12 | 47.0±6 | 41.8±6 | 24.2±12 |
|  | n | 6 | 7 | 8 | 8 | 7 | 8 |

Data are the mean ± SEM of *n* individual data sets. Values were obtained by fitting to the three-parameter logistic model.
<sup>a</sup> The negative logarithm of the agonist concentration required to produce a half-maximal response.
<sup>b</sup> The maximal response to the ligand normalized to the maximal response for GIPR expressed alone, determined by fitting to the three-parameter Log [Agonist] vs response model for each individual experiment.
<sup>c</sup> The negative logarithm of the agonist concentration required to produce a 50% of the first component of the response, determined by fitting to the biphasic Log [Agonist] vs response model
<sup>d</sup> The negative logarithm of the agonist concentration required to produce a 50% of the second component of the response, determined by fitting to the biphasic Log [Agonist] vs response model
<sup>e</sup> The fraction of the concentration-response curve derived from the first, high potency component
Data were assessed for statistical differences, at $p < 0.05$ , in the change in pEC<sub>50</sub> compared to GIPR expressed with pcDNA3.1 using Student's *t* test (\*, $p < 0.05$ ; \*\*, $p < 0.01$ ; \*\*\*, $p < 0.0001$ ).

**Figure 3.**
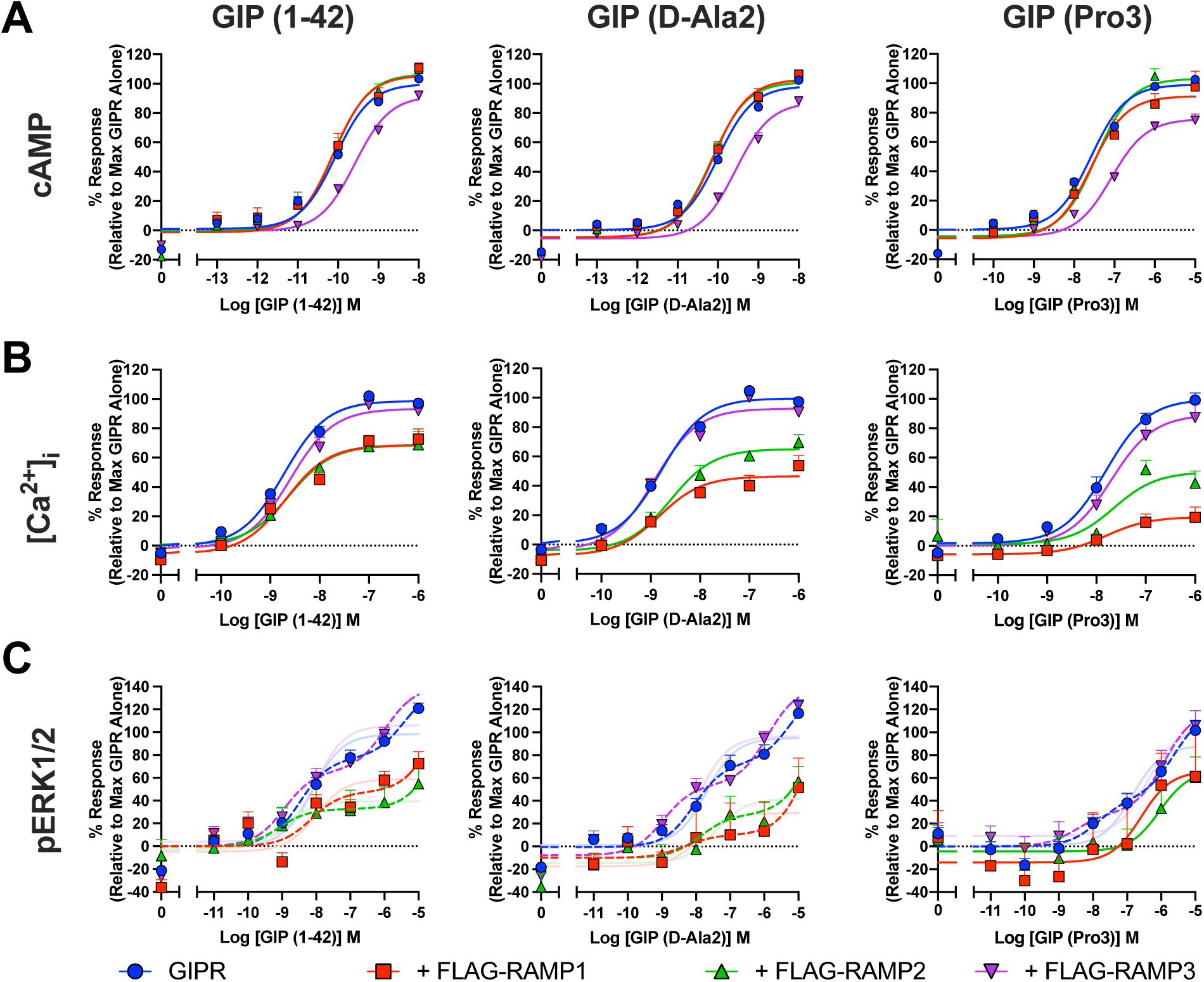
RAMPs differentially modulate GIPR signaling. *A*. cAMP accumulation was determined in HEK-293S cells transfected with GIPR and one of FLAG-RAMP1, FLAG-RAMP2 or FLAG-RAMP3 following 8 min stimulation with GIP (1-42), GIP (D-Ala2) or GIP (Pro3). *B*, [Ca^2+^]_i_ mobilisation was measured in cells transfected with GIPR and one of FLAG-RAMP1, FLAG-RAMP2 or FLAG-RAMP3. Cells were stimulated for 2 min with GIP (1-42), GIP (D-Ala2) or GIP (Pro3) with the intensity at the time of peak response used to construct the concentration-response curves. *C*, ERK1/2 phosphorylation following 5 min stimulation with GIP (1-42), GIP (D-Ala2) or GIP (Pro3) was determined in cells transfected with GIPR and one of FLAG-RAMP1, FLAG-RAMP2 or FLAG-RAMP3. In all cases data are expressed relative to the maximal response, determined by three-parameter logistic fits, in the absence of FLAG-RAMP. Biphasic fits for pERK1/2 are displayed as dashed lines, with the equivalent three parameter fits faded. All data are expressed as mean + SEM of at least 3 individual data sets with quantitative data displayed in (Table 2).

Overall, there is a clear separation of the effects of RAMP3 (to reduce cAMP signalling) and RAMP1 and RAMP2 (to reduce [Ca^2+^]_i_ mobilisation and ERK1/2 phosphorylation) relative to GIPR expressed alone indicating that RAMPs modulate GIPR function.

### RAMPs differentially modulate GIPR signalling by altering G protein activation

As RAMPs can modulate G protein activation of various class B1 GPCRs^16,21,22,35^, we hypothesised that the observed effects on GIPR signalling were a result of changes to the profile of GIPR-mediated G protein activation. Therefore, we investigated the effect of RAMPs on ligand-mediated GIPR G protein activation and β-arrestin recruitment in the presence of each FLAG-RAMP (Figure 4A-L, Figure S7, Table S3, Table 3). Coexpression of RAMP3 with GIPR led to a significant reduction in the potency of G_s_ activation for GIP (1-42) (*p*=0.0003, Figure 4A, Table S3). RAMP3 coexpression had no significant effect on activation of any other G protein or β-arrestin recruitment (Figure 4B-L, Figure S7, Table S3). In stark contrast, coexpression with either RAMP1 or RAMP2 significantly reduced the potency for activation of G_q_, G_11_ and G_15_ (Figure 4E, F and H, Figure S7, Table S3), whilst having no effect on the other G proteins assayed, or β-arrestin1/2 recruitment (Figure 4A-D, G, I-L, Figure S7, Table S3). The changes in potency for GIP (1-42) induced by coexpression with each RAMP relative to GIPR alone are displayed in Figure 4M. Similar effects of RAMPs on GIPR mediated activation of members from each G protein subtype (G_s_, G_q_, G_i2_ and G_12_) were observed for GIP (D-Ala2) and GIP (Pro3) (Figures S8-11. Table S4-5). RAMP3 reduced the potency of activation of G_s_, whilst RAMP1/2 reduced the potency of activation of G_q_. Extending the analysis of the GIP (D-Ala2) G_q_ response to the biphasic model revealed that RAMP1/2 significantly attenuated the fraction of the response mediated by the high potency first component (*p*=0.0066 and *p*=0.0046, respectively). Together these data show that RAMP3 significantly shifts G protein activation away from G_s_, while RAMP1 and RAMP2 significantly shift G protein activation away from G_q_, G_11_ and G_15_.

Combined with the inhibitor data for attenuation of specific signalling pathways shown in Figure 2E-G, these data indicate that the RAMP3-dependent effect on cAMP accumulation may be associated with a reduced ability to activate G_s_, and the RAMP1 and RAMP2-dependent effects on [Ca^2+^]_i_ mobilisation and ERK1/2 phosphorylation may be linked to reduced activation of G_q_, G_11_ and G_15_, but not G_i/o_.

**Table 3:** Potency (pEC_50_) and E_max_ values for β-arrestin-1 and β-arrestin-2 recruitment, following stimulation with GIP (1-42), in HEK 293T cells expressing GIPR and either pcDNA3.1, FLAG-RAMP1, FLAG-RAMP2 or FLAG-RAMP3.

|  |  | 6 Min |  |  |  | 60 Min |  |  |  |
| --- | --- | --- | --- | --- | --- | --- | --- | --- | --- |
|  |  | GIPR | + RAMP1 | + RAMP2 | + RAMP3 | GIPR | + RAMP1 | + RAMP2 | + RAMP3 |
| β-arrestin-1 | pEC<sub>50</sub><sup>a</sup> | 7.80±0.18 | 8.19±0.29 | 7.96±0.36 | 8.08±0.26 | 8.35±0.40 | 8.45±0.36 | 8.08±0.24 | 9.60±0.19* |
|  | E<sub>max</sub><sup>b</sup> | 2.4±0.5 | 4.1±0.6 | 4.1±0.9 | 4.0±0.5 | 4.8±0.7 | 2.7±1.1 | 3.4±1.1 | 4.3±1.0 |
|  | n | 6 | 8 | 7 | 8 | 6 | 8 | 8 | 6 |
| β-arrestin-2 | pEC<sub>50</sub><sup>a</sup> | 8.19±0.12 | 8.35±0.13 | 8.19±0.12 | 8.13±0.13 | 8.77±0.11 | 9.14±0.67 | 8.57±0.37 | 9.04±0.17 |
|  | E<sub>max</sub><sup>b</sup> | 10.7±1.9 | 10.0±1.1 | 11.1±1.8 | 13.2±0.7 | 5.9±0.9 | 6.6±1.3 | 7.5±0.4 | 11.5±1.7* |
|  | n | 4 | 3 | 3 | 4 | 4 | 3 | 3 | 4 |

Data are the mean ± SEM of *n* individual data sets.
<sup>a</sup> The negative logarithm of the agonist concentration required to produce a half-maximal response.
<sup>b</sup> The maximal response to GIP (1-42) minus the baseline in mBRET units.
Data were assessed for statistical differences, at *p*<0.05, in the change in pEC<sub>50</sub> compared to GIPR expressed with pcDNA3.1 using a one-way ANOVA with Dunnett's post-hoc test (\*\*, *p* < 0.01).

**Figure 4.**
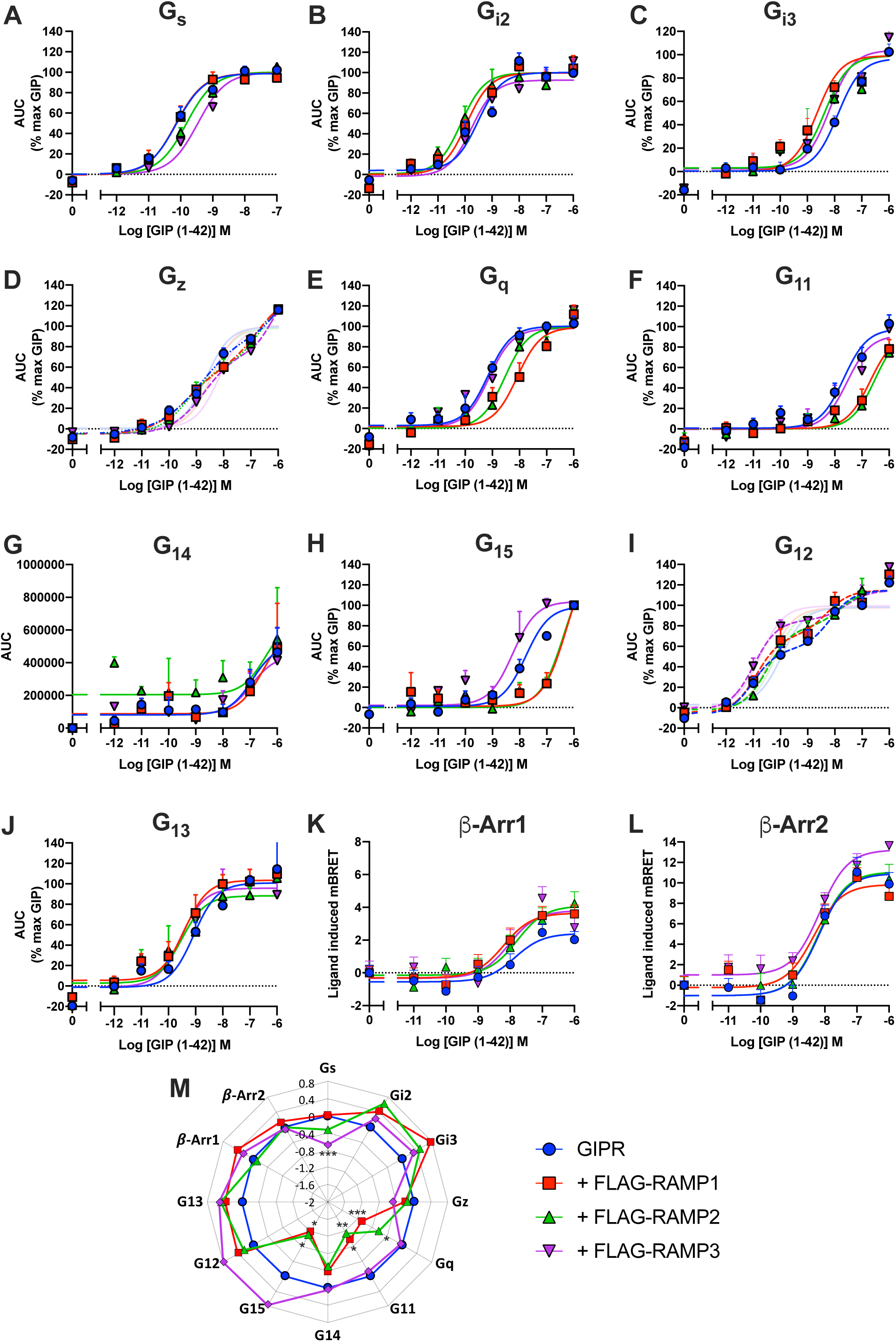
RAMPs differentially alter potency of activation of individual G proteins by GIPR. *A-J*. HEK-293A cells were transfected with GIPR, appropriate Gα-LgBiT and Gβ subunits, Gγ_2_-SmBiT and each FLAG-RAMP/pcDNA3.1 at a 2:1:3:3:2 ratio. RIC8A was also included for G_q_, G_11_, G_14_ and G_15_. G protein activation was measured by the GIP (1-42)-induced change in RLU. Data points were corrected to baseline and vehicle and the AUC used to produce concentration-response curves for each G protein. Data were normalised to the maximum response to GIP (1-42), determined by three-parameter logistic fits, for each condition and are expressed as mean + SEM of 3-7 individual experiments with quantitative data displayed in (Table S3). Biphasic fits for G_z_ and G_12_ are displayed as dashed lines, with the equivalent three parameter fits faded. *K-L*. Peak β-arrestin-1/2 recruitment was measured in HEK-293T cells transiently expressing GIPR-Rluc, GRK5, each FLAG-RAMP/pcDNA3.1 and β-arrestin-1/2-YFP after 6 min stimulation with GIP (1-42). Data are expressed as ligand-induced delta milli BRET (mBRET) and are the mean + SEM of 3-8 individual experiments with quantitative data displayed in (Table 3). *M*. Radial plot showing the change in log potency, obtained from three-parameter logistic fits, of G protein activation and β-arrestin recruitment induced by coexpression with each FLAG-RAMP relative to GIPR + pcDNA3.1. Data were assessed for statistical differences, at *p*<0.05, in pEC_50_ compared to GIPR expressed with pcDNA3.1 using a one-way ANOVA with Dunnett’s post-hoc test (*, *p*<0.05; **, *p*<0.01; ***, *p*<0.001).

### RAMPs control internalisation of GIPR

Beyond effects on G protein activation and ligand specificity, RAMPs are known to influence receptor internalisation and recycling ^17,18,23,36^. We therefore set out to determine whether RAMPs had any effect on GIPR internalisation or recycling using flow cytometry and confocal microscopy (Figure 5).

**Figure 5.**
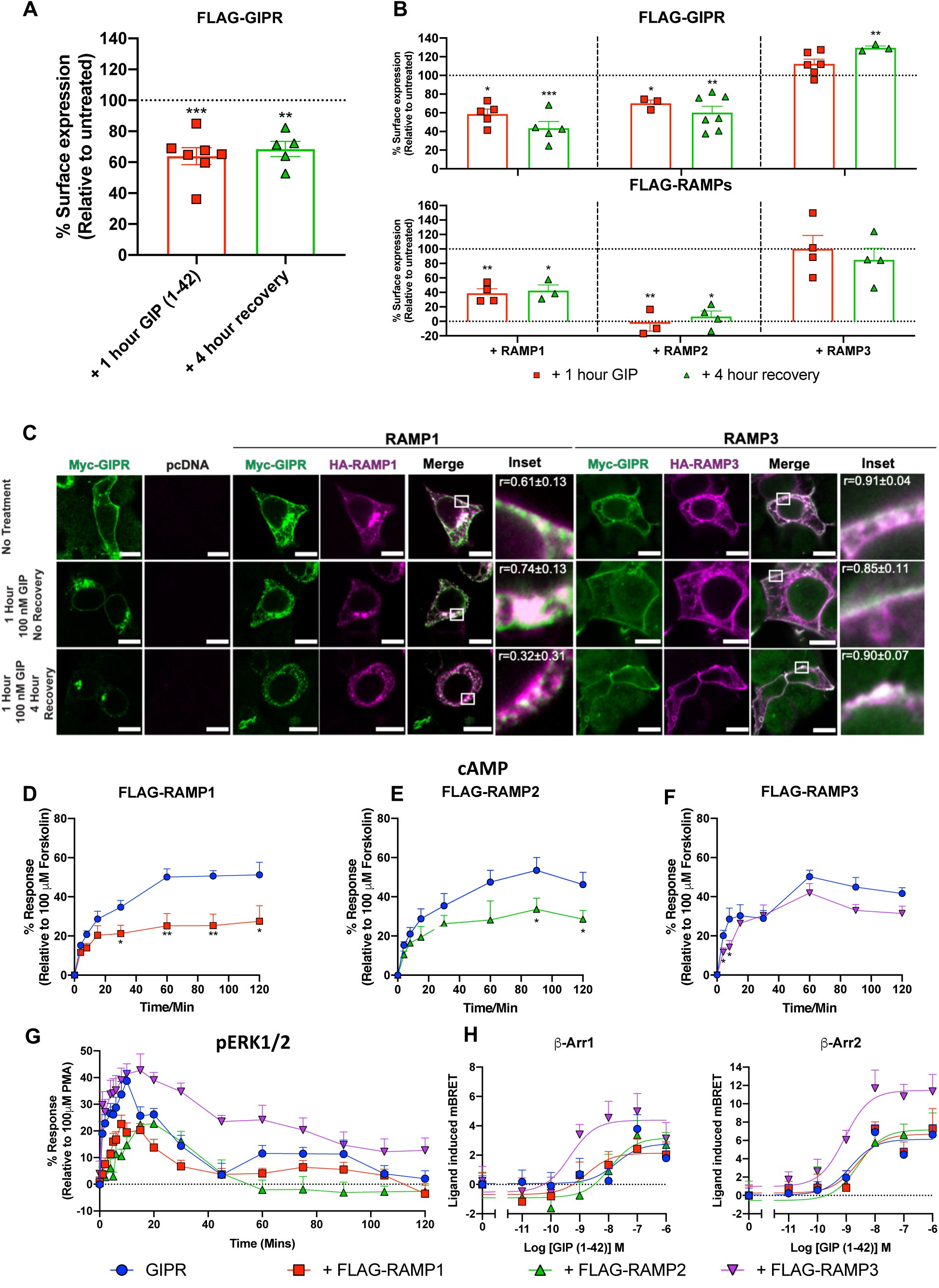
RAMP3 maintains GIPR at the PM. *A-B*. Flow cytometry analysis of GIPR and RAMP PM expression after treatment with GIP (1-42) with or without recovery from agonist stimulation. Cells cotransfected with FLAG-GIPR and pcDNA3.1 (*A*) or GIPR and FLAG-RAMPs (for RAMP surface expression), or FLAG-GIPR and HA-RAMPs (for GIPR surface expression) (*B*) were either treated with 100 nM GIP (1-42) for 1 hour, or treated, washed and allowed to recover for 4 hours, in the presence of cycloheximide. PM expression of FLAG-RAMPs or FLAG-GIPR was determined by flow cytometry as described in Figure 1. Surface expression was normalised to the level observed in the absence of treatment for each GIPR:RAMP complex (as 100%) and pcDNA3.1 (for FLAG-GIPR) or pcDNA3.1 + FLAG-RAMP (as 0%). Data are the mean + S.E.M and were assessed for statistical differences, at *p*<0.05, in surface expression compared to surface expression in the absence of GIP (1-42) treatment using a Kruskal-Wallis test (*, *p*<0.05; **, *p*<0.01; ***, *p<0*.*001*). *C*. HEK-293T cells transfected with GIPR (green) and RAMP1 or RAMP3 (purple) were untreated, treated as in *A*. Scale bar 10 μM. Pearson’s correlation coefficients (r) were determined using the Image J colocalization threshold plugin. Values are mean ± SD. *D-F*. cAMP production was measured following stimulation with 0.1 nM GIP (1-42) in the absence of PDE inhibition in cells transfected with GIPR and FLAG-RAMP/pcDNA3.1. Data were assessed for statistical differences, at *p*<0.05, compared to GIPR expressed with pcDNA3.1 using Student’s t-test (*, *p*<0.05; **, *p*<0.01). *G*. Temporal ERK1/2 phosphorylation following stimulation with 100 nM GIP (1-42) was determined in HEK-293S cells transfected with GIPR and one of FLAG-RAMP1, FLAG-RAMP2 or FLAG-RAMP3 with quantitative data displayed in (Table S7). *H*. β-arrestin-1/2 recruitment was measured in cells expressing GIPR-Rluc, GRK5, FLAG-RAMP/pcDNA3.1 and β-arrestin-1/2-YFP after 60 min stimulation with increasing concentrations of GIP (1-42). Data are expressed as ligand-induced delta mBRET and are the mean + SEM of 3-6 independent experiments with quantitative data displayed in (Table 3).

Using flow cytometry, we assessed GIPR-RAMP internalisation and recycling by measuring cell surface expression of FLAG-GIPR in the presence or absence of HA-RAMPs (addition of the FLAG-tag to GIPR did not influence signalling, whilst CLR signalling was comparable between HA-RAMPs and FLAG-RAMPs - Figure S2) and FLAG-RAMP (with untagged GIPR), in the absence of GIP (1-42), after 1 hour treatment with 100 nM GIP (1-42), and after 4 hours recovery from agonist stimulation in the presence of cycloheximide, to prevent *de novo* protein synthesis (Figure 5A-B). After 1 hour treatment with GIP (1-42), there was a significant reduction in cell surface expression of FLAG-GIPR when expressed alone (Figure 5A), or coexpressed with RAMP1 or RAMP2 (Figure 5B), indicative of homologous desensitisation and internalisation. There was no reduction in cell surface expression of FLAG-GIPR when expressed in HEK-293 cells lacking β-arrestin (Figure S12), suggesting that agonist-stimulated internalisation of GIPR is β-arrestin-dependent. Interestingly, there was no significant change in FLAG-GIPR surface expression when co-expressed with RAMP3 (Figure 5B). A similar pattern was observed for RAMP surface expression, indicating that GIPR and RAMP1 or RAMP2 internalise as a complex, whereas RAMP3 retains the GIPR in a complex at the cell surface. Following 4 hours recovery from agonist stimulation, there was no significant recycling of either GIPR or RAMP in GIPR:pcDNA3.1 (Figure 5A), GIPR:RAMP1 or GIPR:RAMP2 expressing cells (Figure 5B), while surface expression remained at approximately untreated levels in GIPR:RAMP3 expressing cells. This implies that GIPR alone, or in complex with RAMPs, does not recycle to the plasma membrane after ligand-induced internalisation, at least when saturating concentrations of agonist are used.

To confirm these observations, we used confocal microscopy to track the cellular localisation of myc-GIPR-Rluc in the presence and absence of RAMP1 or RAMP3 (Figure 5B). In the absence of RAMP, GIPR was clearly localised to the plasma membrane before treatment, internalised to intracellular vesicles following 1 hour treatment with GIP (1-42) and did not appear to recycle to the plasma membrane after ligand wash-out and 4 hours recovery (Figure 5B). When RAMP1 was coexpressed, colocalisation was observed between GIPR and RAMP1 at the plasma membrane before treatment (r = 0.61±0.13). After treatment with GIP (1-42), both GIPR and RAMP1 were no longer localised to the plasma membrane but remained colocalised in intracellular vesicles (r = 0.74±0.13). Interestingly, after 4 hours recovery, whilst both GIPR and RAMP1 were still localised intracellularly, although reduced colocalisation was observed (r = 0.32±0.31), suggesting that GIPR and RAMP1 may be sorted to separate intracellular trafficking pathways. In the presence of RAMP3, GIPR and RAMP3 were colocalised at the plasma membrane before treatment, after treatment and following 4-hour recovery (r = 0.91±0.04, 0.85±0.11 and 0.90±0.07, respectively), supporting the conclusion that RAMP3 prevents internalisation of the GIPR.

### RAMPs regulate temporal dynamics of GIPR signalling

The differences in cellular localisation of GIPR in the presence of RAMPs, raised the possibility that there may be differences in the temporal profile of signalling of the GIPR when co-expressed with RAMPs. To investigate this, we assayed the cAMP level in the absence of PDE inhibitor up to 2 hours post-stimulation with an ∼EC_50_ concentration of GIP (1-42) (0.1 nM) in the presence and absence of FLAG-RAMPs (Figure 5C-E). Consistent with earlier experiments performed with 8 minutes of stimulation, after 4 or 8 minutes of agonist stimulation, the GIPR in the presence of RAMP3 displayed lower levels of cAMP (Figure 5E), whilst in the presence of RAMP1 and RAMP2 had no significant effect relative to GIPR expressed alone (Figure 5C-D). As duration of agonist stimulation increased, there was no further attenuation of GIPR-mediated cAMP accumulation in the presence of RAMP3, although there were small significant reductions after both 90 and 120 minutes stimulation (Figure 5E). GIPR co-expression with RAMP1 or RAMP2, on the other hand, resulted in progressively greater reductions of cAMP with significance reached for RAMP1 after 30 minutes stimulation and RAMP2 after 90 minutes stimulation (Figure 5C-D). Given the previous observations that RAMP1/2 had no effect on G_s_ activation or GIPR internalisation, these data again suggest that GIPR and RAMP1/2 may be sorted to different intracellular trafficking pathways to GIPR alone.

Long term ERK1/2 phosphorylation was also assayed with or without FLAG-RAMP1, FLAG-RAMP2 or FLAG-RAMP3 in response to 100 nM GIP (1-42) (Figure 5F). In the absence of RAMP, GIPR-stimulated ERK1/2 phosphorylation reached a maximum between 4-10 minutes, decayed to basal levels after 40 minutes and displayed a small second phase after around 45 minutes stimulation. This second phase of ERK1/2 phosphorylation was determined to be β-arrestin-dependent as it was absent in HEK-293AΔβ-arrestin cells expressing the GIPR (Figure S13, Table S6). Coexpression with RAMP1 or RAMP2 resulted in a reduction in amplitude of the first phase relative to GIPR expressed alone (Figure 5F, Table S7). Additionally, the second phase was attenuated in the presence of RAMP1 and absent with RAMP2. RAMP3 coexpression, on the other hand, significantly prolonged ERK1/2 phosphorylation, with levels of activation declining much more gradually, with a sustained response even after 120 minutes. Therefore, as well as modulating G protein activation, RAMPs can modulate GIPR signalling in response to chronic agonist stimulation.

### RAMP3 enhances β-arrestin recruitment to GIPR

The fact that the second phase of ERK1/2 phosphorylation was β-arrestin-dependent, and RAMPs appeared to modulate long term ERK1/2 signalling raised the possibility that RAMPs may also alter the duration of β-arrestin recruitment to the GIPR. β-arrestin1/2 recruitment was, therefore, measured in the presence and absence of each FLAG-RAMP after 60-minute stimulation with increasing concentrations of GIP (1-42) (Figure 5G, Table 3). Coexpression of RAMP3 with GIPR increased the potency of β-arrestin1 recruitment and increased the E_max_ of β-arrestin-2 recruitment after 60 minutes stimulation, relative to GIPR expressed alone. This demonstrates RAMP3-specific effects on sustained β-arrestin recruitment, but also demonstrates that RAMP3 does not prevent internalisation by blocking β-arrestin recruitment, even though β-arrestin is required for internalisation.

Overall, these data indicate that RAMPs alter the temporal profile of intracellular signalling, as well as influencing the cellular localisation of the GIPR. Thus, it is plausible that the effects of RAMPs on the long-term signalling of the GIPR are, at least in part, due to the RAMP-induced changes to cellular localisation of GIPR.

### RAMP3 effects on GIPR internalisation are dependent on its PDZ-motif

The C-terminal PDZ-motif in RAMP3 is required for promoting recycling of CLR and ACKR3 (via interaction with NSF)^17,23^ and for reducing agonist-stimulated internalisation of CLR (via interaction with NHERF1)^37^. We therefore hypothesised that the observed effects of RAMP3 on GIPR internalisation were dependent on the PDZ-motif of RAMP3, which is not present in RAMP1 or RAMP2.

To investigate the role that the PDZ-motif of RAMP3 plays in controlling GIPR membrane localisation, we deleted the last 4 amino acids from RAMP3 to generate SNAP-, HA- and FLAG-RAMP3ΔPDZ fusion constructs. These constructs were functional, as coexpression with CLR resulted in similar acute cAMP responses to that of CLR:RAMP3 (Figure S2). Firstly, GIPR promoted cell surface expression of RAMP3ΔPDZ to similar levels as wild type (WT) RAMP3, as assessed using flow cytometry, and a direct, plasma membrane localised, interaction was also observed using cell surface BRET, (Figure S14A-B). Similar effects on FLAG-GIPR plasma membrane expression, compared to WT RAMP3, were observed upon coexpression with RAMP3ΔPDZ (Figure S14C).

We next explored GIPR: RAMP3ΔPDZ internalisation (Figure 6A-B). In contrast to WT RAMP3, after 1 hour treatment with GIP (1-42), there was a significant reduction in plasma membrane expression of the GIPR:RAMP3ΔPDZ complex, which was clearly visualised intracellularly. (Figure 6A-B). This indicates that, similar to CLR, the PDZ-motif of RAMP3 may be responsible for preventing internalisation of the GIPR. Interestingly, following ligand washout and 4 hours recovery, the GIPR:RAMP3ΔPDZ complex displayed plasma membrane localisation to a similar extent to untreated cells, suggesting receptor recycling back to the plasma membrane. This recycling demonstrates that removal of the PDZ-motif produces a GIPR:RAMP complex with a distinct phenotype to that of GIPR alone or in combination with any WT RAMP.

**Figure 6.**
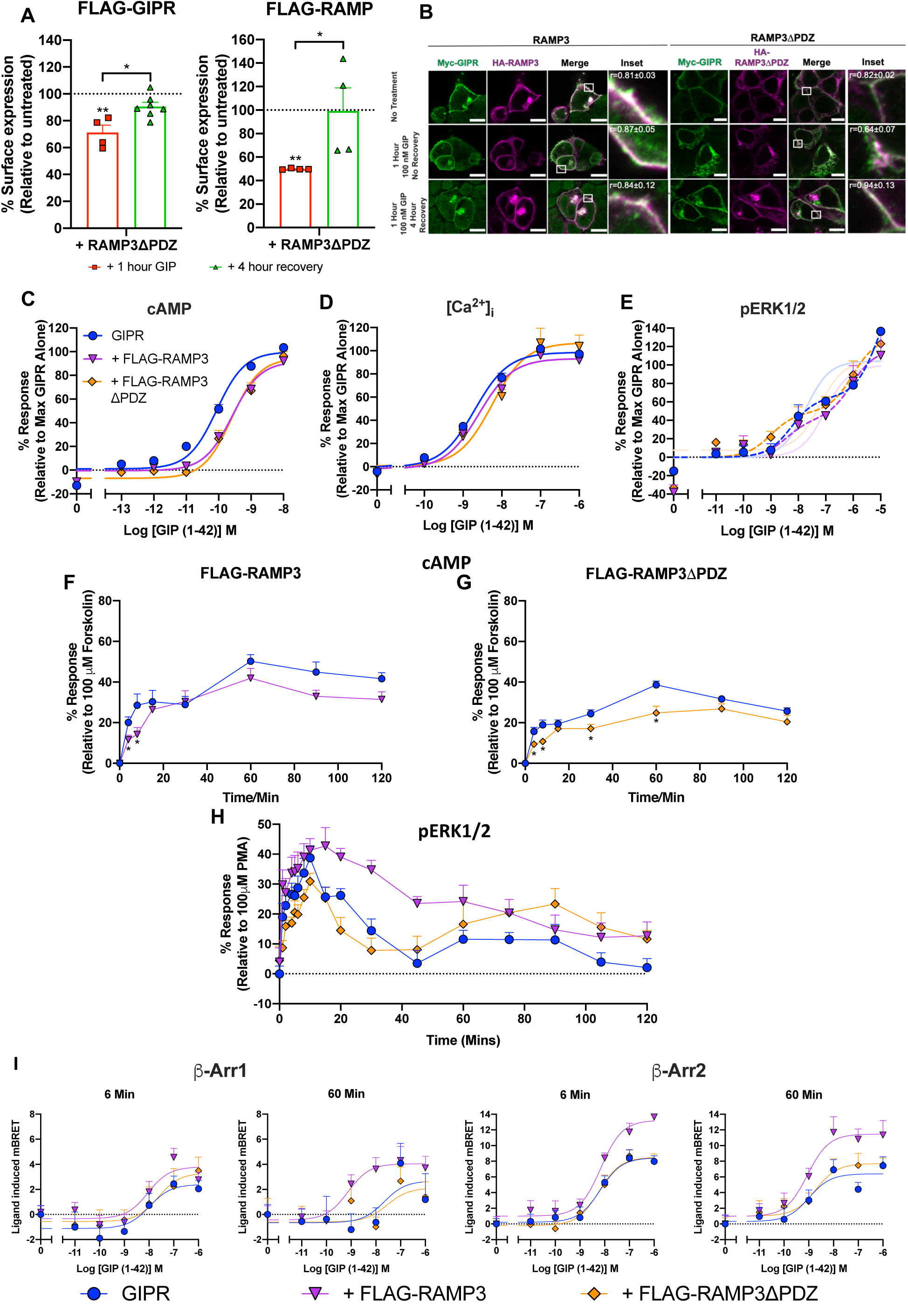
RAMP3 maintenance of GIPR at the plasma membrane is dependent on it PDZ-recognition sequence. *A*. HEK-293S cells cotransfected with GIPR and FLAG-RAMP3ΔPDZ (for RAMP surface expression), or FLAG-GIPR and HA-RAMP3ΔPDZ (for GIPR surface expression) were either treated with 100 nM GIP (1-42) or vehicle for 1 hour or treated with 100 nM GIP (1-42) for 1 hour, washed and allowed to recover for 4 hours, in the presence of cycloheximide. PM expression of FLAG-RAMP3ΔPDZ or FLAG-GIPR was determined by flow cytometry as described in Figure 1. Surface expression was normalised to the level observed in the absence of treatment (as 100%) and pcDNA3.1 (for FLAG-GIPR) or pcDNA3.1 + FLAG-RAMP3ΔPDZ (as 0%). Data are the mean + S.E.M and were assessed for statistical differences, at *p*<0.05, in the change in pEC_50_ compared to surface expression compared to surface expression in the absence of GIP (1-42) treatment using a Kruskal-Wallis test (*, *p*<0.05; **, *p*<0.01). *B*. HEK-293T cells transfected with GIPR and RAMP3ΔPDZ were treated as in *A*. Scale bar 10 μM. Pearson’s correlation coefficients (r) were determined using the Image J colocalization threshold plugin. Values are mean ± SD. *C-E*. cAMP accumulation (*C*), [Ca^2+^]_i_ mobilisation (*D*) and ERK1/2 phosphorylation (*D*) were determined in HEK-293S cells expressing GIPR and either FLAG-RAMP3 or FLAG-RAMP3ΔPDZ as previously described in Figure 2. Data are expressed relative to the maximal response, determined by three-parameter logistic fits, in the absence of FLAG-RAMP. Biphasic fits for pERK1/2 are displayed as dashed lines, with the equivalent three parameter fits faded. *F-G*. cAMP production was measured following stimulation with 100 nM GIP (1-42) in the absence of PDE inhibition in HEK-293S cells transfected with GIPR and one of FLAG-RAMP3, FLAG-RAMP3ΔPDZ or pcDNA3.1. Data were assessed for statistical differences, at *p*<0.05, compared to GIPR expressed with pcDNA3.1 using Student’s t-test (*, *p*<0.05). *H*. Temporal ERK1/2 phosphorylation following stimulation with 100 nM GIP (1-42) was determined in cells transfected with GIPR and FLAG-RAMP3ΔPDZ with quantitative data displayed in (Table S8) *I*. β-arrestin-1/2 recruitment was measured in HEK-293T cells expressing GIPR-Rluc, GRK5, FLAG-RAMP3ΔPDZ/pcDNA3.1 and β-arrestin-1/2-YFP after 6 min and 60 min stimulation with 100 nM GIP (1-42). Data are expressed as ligand-induced delta mBRET and are the mean + SEM of 4-6 independent data sets with quantitative data displayed in (Table S9).

cAMP accumulation, [Ca^2+^]_i_ mobilisation and ERK1/2 phosphorylation were measured in response to GIP (1-42), in HEK-293S cells expressing GIPR and either WT RAMP3 or RAMP3ΔPDZ to assess the effect of the PDZ-motif on intracellular signalling (Figure 6C-E). The absence of the PDZ-motif had no significant effect on the initial phase of intracellular signalling, thus indicating that receptor complexes with RAMP3ΔPDZ have the same intrinsic ability to activate second messengers as WT RAMP3. When determining temporal cAMP levels, RAMP3ΔPDZ coexpression resulted in the expected reduction at 4 and 8 minutes, similar to WT RAMP3 (Figure 6F-G). Similar to WT RAMP3, there was no significant effect on cAMP levels after 15, 90, or 120 minutes stimulation, although, in contrast to WT RAMP3, there was a significant reduction after 30 and 60 minutes stimulation. When assaying long term ERK1/2 phosphorylation, GIPR:RAMP3ΔPDZ signalling more closely matched that of GIPR alone, with a rapid decline in initial ERK1/2 signalling and an even more pronounced second phase (Figure 6H and Table S8). Furthermore, the enhancement of β-arrestin recruitment observed with RAMP3 was abolished by removal of the PDZ-motif (Figure 6I and Table S9). This provides further evidence consistent with internalisation of the GIPR:RAMP3ΔPDZ complex, and the prolongation of ERK1/2 phosphorylation by RAMP3 that is a result of sustained β-arrestin recruitment from maintained cell surface expression. Overall, these data provide evidence that the PDZ-motif at the C-terminal tail of RAMP3 is necessary for preventing internalisation of the GIPR and that this modulates long-term cAMP and ERK1/2signalling of the GIPR.

### RAMPs are coexpressed with GIPR in pancreatic islets and RAMP1^-/-^ alters regulation of blood glucose levels

Having established that RAMPs significantly alter signalling and cellular fate of GIPR, it was important to establish whether this interaction was physiologically relevant. GIP, as an incretin hormone, plays a crucial role in the maintenance of blood glucose levels, by promoting insulin and glucagon secretion from pancreatic β- and α-cells, respectively (Figure 7A). As this is the most well characterised function of GIPR, we have focused on pancreatic islets and insulin secretion. We firstly determined RNA expression levels of RAMP1 and RAMP3 in mouse pancreatic islets, using RNAscope to detect RAMP1 and RAMP3 transcripts (RAMP2 was not included as the effects of RAMP2 knockout could not be determined in subsequent mouse experiments^38^) in cells positive for glucagon (α-cells) or insulin (β-cells) (Figure 7B). The average signal intensity per cell, indicates that both RAMP1 and RAMP3 are expressed in α- and β-cells, with significantly greater expression at the cellular level in α-cells (Figure 7C).

**Figure 7.**
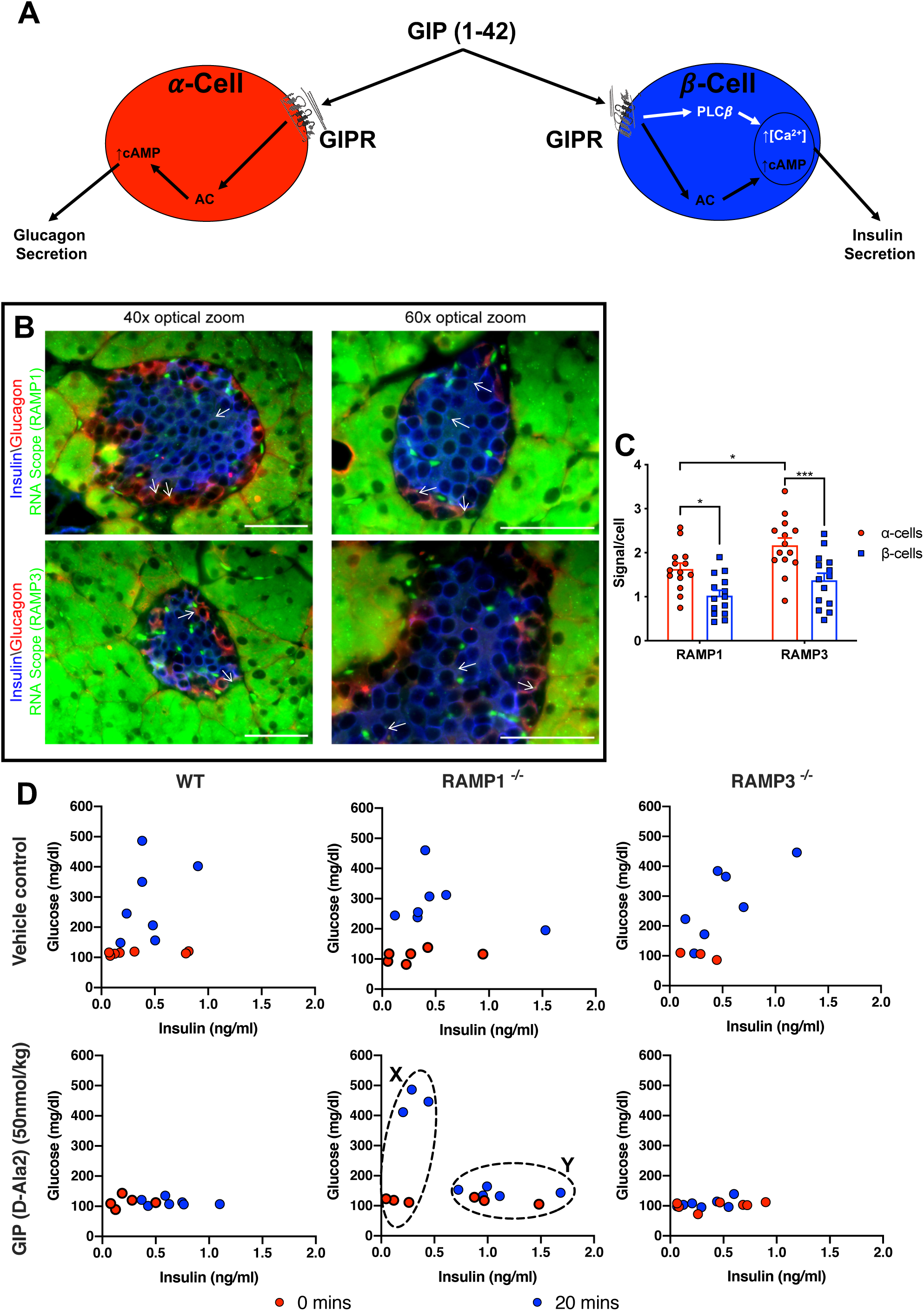
RAMPs are coexpressed with GIPR in pancreatic α- and β-cells and regulate the normal physiological function of GIPR. *A*. Schematic diagram representing the physiological effects of GIP (1-42) on pancreatic α- and β-cells. *B*. Pancreatic tissue from six wildtype 129/S6-SvEv-TC1 mice was fixed and stained for glucagon (red) and insulin (blue) to differentiate between pancreatic α- and β-cells. RNAscope *in situ* hybridisation was performed on fixed pancreatic tissue to detect RNA transcripts for RAMP1 and RAMP3 at the cellular level (green, indicated by white arrows). Scale bar 50 μm. *C*. Average signal intensity per cell for RAMP1 or RAMP3 RNA transcripts in pancreatic α- and β-cells. Data are the mean + SEM of 14 individual cells. Data were assessed for statistical differences, at *p*<0.05, in RAMP expression using a one-way ANOVA with Dunnett’s post-hoc test (*, *p*<0.05; ***, *p*<0.001). *D*. Blood glucose and insulin levels were measured in WT, RAMP1^-/-^ and RAMP3^-/-^ 129/S6-SvEv-TC1 mice immediately before (red circles), and 20 min after (blue circles) intraperitoneal injection with 1g/kg glucose with or without 50 nmol/kg GIP (D-Ala2). Data are expressed as the insulin vs glucose concentration for each individual mouse. Dashed circled areas represent two distinct populations within the RAMP1^-/-^ mice treated with GIP (D-Ala2).

Having determined that GIPR and RAMPs are coexpressed, at the mRNA level, in mouse pancreatic islets we next explored whether RAMPs play any role in the normal insulinotropic action of GIP. For this purpose, we exposed WT, RAMP1^-/-^ and RAMP3^-/-^ mice^38,39^ to intraperitoneal glucose challenge in the presence or absence of metabolically stabilised GIP (D-Ala2) (to prolong the circulating half-life of GIP by preventing breakdown by DPPIV) (Figure 7D). Immediately before injection, resting blood glucose levels were all approximately 100 mg/dL. As expected, in the absence of GIP, blood glucose levels were elevated 20 minutes after injection in WT, RAMP1^-/-^ and RAMP3^-/-^ mice. For WT and RAMP3^-/-^ mice glucose levels after 20 minutes were similar to resting levels, in the presence of GIP (D-Ala2), indicative of GIP potentiation of glucose-stimulated insulin secretion. However, there were two distinct populations of RAMP1^-/-^ mice after 20 minutes treatment in the presence of GIP (D-Ala2). One population (Figure 7D, marked x) displayed elevated blood glucose levels, suggesting an insensitivity to GIP. The other population (Figure 7D, marked y) exhibited resting glucose levels but elevated insulin levels. Therefore, it appears that RAMP1 is required for the normal functioning of GIPR in pancreatic islets and thus suggests interactions between GIPR and RAMPs are potentially important for aspects of GIPR physiology.

## Discussion

The pharmacology of the GIPR has, to date, been relatively poorly characterised. In this study, we have shown that GIPR pleiotropically activates multiple different G proteins and β-arrestins to stimulate cAMP accumulation, release of intracellular calcium and phosphorylation of ERK1/2. In addition, activation of the GIPR by GIP peptides promotes receptor internalisation. Moreover, we have demonstrated novel interactions between GIPR and RAMPs. These interactions modulate the selectivity profile of GIPR for G protein activation to influence the initial phase of intracellular signalling events, alter the cellular localisation of GIPR, control its long-term activity in terms of signalling and, are important for the normal physiological functioning of the GIPR.

### Widespread interactions of RAMPs with Family B GPCRs

RAMPs heterodimerise with select class B1 GPCRs to modulate their pharmacology to effectively create “new” receptors with distinct characteristics^14,16,22,27^. We utilised BRET and flow cytometry methods to screen all class B1 GPCRs for interactions with, and effects on cell surface expression of, RAMPs. The BRET screen indicated that almost all class B1 GPCRs could interact, with at least one RAMP – a similar finding to a recent multiplexed suspension bead array (SBA) approach^20^. Our flow cytometry screen correlated well with positive and negative results from previous studies investigating the effects of class B1 GPCRs on RAMP plasma membrane expression^15^, verifying its validity as a method for investigating GPCR: RAMP interactions. As a result of this method, SCTR, PAC1R, GHRHR and GIPR, were identified to promote plasma membrane localisation of RAMP1, RAMP1, RAMP2 and all three RAMPs, respectively. Despite this, there were some differences to published literature. CRFR1 has been reported to promote plasma membrane localisation of RAMP2^21^, whilst VPAC1R has been reported to promote surface expression of all three RAMPs^40^. Whilst significant increases were not detected in this study, surface expression trended towards an increase in each case. Similarly, there was a trend towards an increase in RAMP3 surface expression when coexpressed with SCTR, an interaction previously reported^26^, whilst we also observed a significant elevation in RAMP1 cell surface expression with SCTR. Identification of RAMP interactions have not always been consistent between studies^15^, with discrepancies reported for both VPAC2R^21,40^ and GCGR^22,36^. It is, therefore, clear that newly identified interactions must be treated cautiously, verified further and investigated for functional effects. As such, ELISA and cell surface BRET measurements, along with functional assays, were utilised to confirm interaction of GIPR with all three RAMPs. Nonetheless, the evidence now suggests that a much greater array of GPCRs, than perhaps previously thought, may interact with RAMPs.

Interestingly, only around 50% of the interactions identified in the SBA assay^20^ or our BRET screen appear to translate to effects upon RAMP plasma membrane expression. By measuring BRET between Nluc-GPCR and SNAP-RAMPs only at the cell surface we showed that there was no, or minimal, BRET between Nluc-CRFR2 and SNAP-RAMPs, in stark contrast to the initial BRET screen and SBA study^20^. This suggests that not all GPCR:RAMP complexes traffic to the PM, most likely because the complex is either targeted for degradation or because the receptor resides largely intracellularly. As well as the well-established role of RAMPs to chaperone GPCRs to the cell surface^14,41,42^, our data suggests that RAMPs may also play a role in trapping some GPCRs intracellularly. The latter may be particularly applicable to GPCR:RAMP3 complexes as RAMP3 has been demonstrated, in multiple cell lines, to endogenously traffic to the plasma membrane^23,25^. N-glycosylation of RAMP3 has been attributed to its receptor-independent cell surface localisation^43^, although the endogenous expression of interacting GPCRs, such as CTR and CLR, may also influence this localisation. Therefore, any interaction between RAMP3 and a receptor that resides, at least partially, intracellularly in the basal state may reduce endogenous trafficking of RAMP3 to the PM, thus resulting in a cellular redistribution of RAMP3. Indeed, the atypical chemokine receptor, ACKR3, which resides on the membrane of endocytic vesicles in the resting state^44^, was recently shown to reduce PM expression of RAMP3^23^. Furthermore, intracellular GPCR-RAMP interactions may have interesting implications on GPCRs with high constitutive activity, as well as those that signal intracellularly.

It is important to note that GPCR-RAMP interactions are not limited to class B1 GPCRs as RAMP1 and RAMP3 have been shown to interact with the class C calcium sensing receptor (CaSR)^42^ whilst RAMP3 also interacts with the class A G protein coupled estrogen receptor (GPER)^41^. Recently, a flow cytometry screen of the chemokine family of receptors identified numerous GPCRs that could alter the plasma membrane expression of RAMPs^23^. Furthermore, the strong co-evolution of RAMPs and GPCRs suggests that there are likely to be many more interacting partners^45^. It will be interesting to observe how RAMPs modulate the pharmacology, trafficking and signalling properties of this wide range of GCPRs.

### Signalling pleiotropy at the GIPR

Having identified GIPR to promote RAMP cell surface expression, NanoBiT® assays, mammalian cell signalling assays and chemical inhibitors were used to characterise the signalling profile of the GIPR. Although classically thought of as a G_s_ coupled receptor, GIPR can activate, to varying degrees, G proteins from all 4 subfamilies. Furthermore, despite β-arrestin recruitment to the GIPR being somewhat debated^46–48^, we demonstrated rapid β-arrestin1 and β-arrestin2 recruitment. Unsurprisingly, G_s_ was the most potent effector activated, but our data strongly support promiscuous coupling of the of GIPR to multiple G protein subtypes, albeit with varying potencies and differing components to the response. This G protein promiscuity translated to pleiotropic signalling, with GIPR stimulating [Ca^2+^]_i_ mobilisation and ERK1/2 phosphorylation, together with its well-known role to promote cAMP accumulation. Somewhat surprisingly, given the role of G_i/o_ at other class B1 GPCRs^16,22^, PTX treatment had no effect on cAMP accumulation, but reduced [Ca^2+^]_i_ mobilisation and ERK1/2 phosphorylation. The expression levels of each adenylyl cyclase (AC) isoform are not known in the cell lines used in this study. Therefore, it is possible that G_i_-sensitive AC isoforms are either absent or expressed at very low levels. Alternatively, GIPR may be located in subdomains of the plasma membrane lacking G_i_-sensitive AC. The insulinotropic actions of GIPR are thought to be mediated through activation of the cAMP effectors, PKA and EPAC^49^, to ultimately elevate [Ca^2+^]_i_ levels. The potent (around nM), G_q/11/14_, G_i/o_ and EPAC1/2-dependent, activation of Ca^2+^ release from intracellular stores by GIP, may therefore also contribute to stimulation of insulin release, as is the case for GLP-1^50^. While G_q/11/14_ activation promotes [Ca^2+^]_i_ release via PLCβ activation, it is possible that G_i/o_-dependent release of Gβγ to activate PLCβ, PLCε or PLCη is responsible for the G_i/o_ component^51,52^. The GIPR is known to activate ERK1/2 phosphorylation^13^, and here we provide evidence that this occurs via a range of different intracellular mechanisms. Treatment with pharmacological inhibitors revealed that the first, high potency component is G_q/11/14_ and G_i/o_-dependent, with the smaller low potency component abolished by EPAC1/2 inhibition. However, further experiments will be required to confirm how GIPR-mediates ERK1/2 phosphorylation. The G_q/11/14_ mechanism implied for other receptors involves production of diacylglycerol (DAG) to activate protein kinase C (PKC)^53^, therefore the G_i/o_ mechanism could involve activation of Rap1GAPII or Gβγ-mediated activation of tyrosine kinases, similar to the M2 muscarinic acetylcholine receptor and α_2A_-adrenoreceptor, respectively^53,54^. It is also interesting to note the potent activation of G_12_ and G_13_, especially due to the apparent negative effect of RhoA/ROCK activation on insulin secretion^55,56^.

### RAMPs as allosteric modulators of the GIPR

In HEK-293S cells, RAMP coexpression with GIPR was observed to differentially modulate the initial phase of second messenger signalling pathways: RAMP3 attenuated cAMP signalling, while RAMP1 and RAMP2 abrogated calcium and pERK1/2 signalling. It is not surprising that the effects of the RAMPs on [Ca^2+^]_i_ mobilisation and ERK1/2 phosphorylation were similar as both were predominantly G_q/11/14_ and G_i/o_-dependent with a small contribution from EPAC1/2. The attenuated activation of G_q/11/15_, coupled with the reduction to the G_q/11/14_ and G_i/o_-dependent high potency phase of the ERK1/2 response upon coexpression of RAMP1/2 indicates that the impaired ERK1/2 signalling, and likely [Ca^2+^]_i_ mobilisation, are due to negative regulation of G_q/11/15_. Similarly, the reduced cAMP signalling in the presence of RAMP3 is most likely a result of reduced G_s_ activation. It is also important to note that, despite GIP (Pro3) displaying biased agonism towards pERK1/2 signalling, relative to cAMP accumulation, the effects of RAMP coexpression on intracellular signalling and G protein activation were broadly the same as GIP (1-42) and the unbiased ligand, GIP (D-Ala2).

The multiple domains of RAMPs (extracellular domain, transmembrane domain and C-terminus) allows them to allosterically and directly modulate ligand binding, propagation of receptor activation to the intracellular side of the receptor and, G protein coupling and activation^57,58^. Our G protein dissociation studies suggest that RAMPs appear to differentially modulate G protein activation to alter GIPR signalling. This has also previously been observed for RAMP interactions with CLR, CTR, CRFR1, VPAC2R and GCGR^16,21,22,25,35^. Whilst the effects of RAMPs on second messenger signalling events can be explained by alterations to G protein coupling, it will be important in future studies to establish whether there are also RAMP-induced effects on ligand binding. The mechanism by which the RAMPs shift the rank order of potency of G protein activation may be via allosterically modifying the G protein binding pocket to differentially promote or disrupt receptor induced G protein activation. Alternatively, there may be direct interactions of the RAMP C-terminus with the G protein. Cryo-EM structures of CLR with each RAMP suggest that both mechanisms are plausible^59,60^.

### RAMPs spatially modulate GIPR signalling

To date, most studies involving RAMPs have focused on effects on ligand binding, G protein activation and early phase signalling events. However, it has emerged that RAMPs also influence the cellular fate of interacting receptors, with RAMP3 shown to interact with NSF to promote recycling of CLR and ACKR3^17,23^, or NHERF1 to reduce CLR internalisation^37^. Through flow cytometry and confocal microscopy, we have been able to track GIPR and RAMPs following stimulation with GIP. RAMP coexpression was demonstrated to modulate both GIPR internalisation and receptor signalling following chronic stimulation. RAMP1/2 had no effect on GIP (1-42)-mediated internalisation but progressively reduced cAMP signalling over time, relative to GIPR, and lacked any second phase of ERK1/2 phosphorylation. It is plausible that the altered long-term signalling of GIPR in the presence of RAMP1/2 may be explained by changes to receptor fate. There are contrasting findings regarding the fate of GIPR with separate studies suggesting slow recycling^47^ or lysosomal degradation^61^. It will also be important to establish the impact of RAMPs on the endosomal localisation of GIPR using markers for Rabs^62^. Whilst there appears to be no recycling of GIPR in the absence or presence of RAMP1/2 at saturating agonist concentration. It should, however, be noted that recycling cannot be ruled out for lower concentrations of agonist^63^.

Furthermore, we have demonstrated a dramatic alteration in GIPR localisation in the presence of RAMP3. GIPR: RAMP3 complexes were observed at the plasma membrane after both 1-hour stimulation and a further 4 hours recovery. The prolonged first phase of ERK1/2 phosphorylation, coupled with sustained cAMP production, compared to RAMP1/2, suggest that these effects may be a result of sustained plasma membrane localisation. While the data suggests the GIPR does not internalise when expressed with RAMP3, another possible explanation is that the GIPR internalises to very early endosomes (VEE)^64^, whereby it rapidly recycles to the membrane, thus appearing localised to the membrane in the confocal images and maintaining a high level of plasma membrane localisation in FACS measurements.

GIPR internalisation and recycling has proved a controversial topic, with some studies reporting ligand-dependent GIPR internalisation with little recycling^61^, others suggesting rapid, constitutive internalisation and recycling with no change upon ligand addition^47,65^ or, no internalisation at all^66^. The ability of RAMPs to alter the profile of GIPR internalisation, trafficking and recycling observed in this study may suggest that the conflicting data in the literature, could be associated with potential differences in endogenous RAMP expression levels in different cell lines used in previous studies.

PDZ-domain-containing proteins regulate endosomal sorting of a number of GPCRs, including β2-adrenergic receptor (β2AR) and luteinising hormone receptor (LHR)^67,68^ to promote rapid recycling to the membrane. These effects are dependent upon PDZ recognition sequences at the C-terminus of the receptors. RAMP3, but not RAMP1 or RAMP2, also possesses a C-terminal PDZ-recognition sequence, which is required for promotion of plasma membrane expression of CLR via interaction with NSF or NHERF-1^17,18,37^. In the case of GIPR:RAMP3, the PDZ-recognition sequence is required for maintenance of cell surface expression as deletion resulted in agonist-stimulated internalisation of the GIPR:RAMP3ΔPDZ complex. Removal of the PDZ-recognition motif had no effect on early phase intracellular signalling events, relative to RAMP3, indicating that the ability to activate second messengers was maintained. In contrast, the sustained ERK1/2 phosphorylation observed in the presence of RAMP3 was lost when the PDZ domain was removed, providing evidence that the effect of RAMP3 on ERK1/2 signalling is likely due to maintained plasma membrane localisation. Interestingly, and somewhat surprisingly, the GIPR:RAMP3ΔPDZ complex recycled back to the plasma membrane following 4 hours recovery from agonist stimulation. This recycling indicates that there may be a PDZ-independent mechanism intrinsic to RAMP3 that promotes plasma membrane expression; possibly N-glycosylation or another, as yet unidentified, accessory protein.

The addition of GIPR to the family of GPCRs that are regulated by RAMP3 raises an interesting question regarding the more general role of RAMP3. The effect on the initial phase of signalling for CLR and ACKR3 are not hugely different to CLR:RAMP1/2 or ACKR3 alone, respectively^16,23^. Therefore, the predominant physiological role of RAMP3 may be to regulate the recycling properties and plasma membrane localisation of the interacting receptor.

### The importance of RAMP3 for potentiating β-arrestin recruitment

We provide evidence that β-arrestins are responsible for a second phase of ERK signalling, a feature that is common to many other GPCRs^69,70^. Despite rapid recruitment of β-arrestins, we also demonstrate sustained interaction with the GIPR, and this is further enhanced for β-arrestin2 upon coexpression with RAMP3. The importance of β-arrestin recruitment in GLP-1R-mediated insulin release indicates that they may also play a role in mediating the insulinotropic actions of GIPR. Given the effects of RAMP3 on β-arrestin recruitment and ERK1/2 activation and that ERK1/2 signalling is thought to promote proliferation of β-cells^1,49,71^ it is possible that RAMP3 plays a role in regulating GIP-mediated pancreatic β-cell proliferation.

### The physiological consequence of the RAMP-GIPR interactions

We have demonstrated that RAMPs, particularly RAMP1, play a role in the normal physiological functioning of GIPR in pancreatic islets to regulate blood glucose levels. Through RNAscope we have shown that RAMPs are coexpressed, at the RNA level, with the GIPR in mouse pancreatic α- and β-cells and that RAMP1^-/-^ mice display impaired GIP-mediated regulation of blood glucose levels. Although RAMP3^-/-^ mice did not demonstrate impaired regulation of insulin secretion or blood glucose levels, RAMP3 may still play an important role in pancreatic β-cell proliferation or in GIPR functioning away from the pancreas. It should be noted that it is not known whether GIPR expression levels are altered in the RAMP1^-/-^ or RAMP3^-/-^ mice.

This study has demonstrated the influence of RAMP interactions on GIPR pharmacology and highlights the importance of considering RAMPs when assessing GIPR signaling in recombinant systems. The apparent importance of RAMP1 to GIPR signalling in pancreatic islets raises the possibility of selectively targeting GIPR: RAMP1 complexes through the GIPR: RAMP1 interface as a treatment for type 2 diabetes mellitus. Indeed, exploiting the receptor:RAMP interface for selective drug design has recently been achieved for the anti-migraine drug, erunumab, which selectively targets the CLR: RAMP1 interface^72,73^.

## Materials and Methods

### Peptides

Human GIP (1-42), GIP (D-Ala2), GIP (Pro3), GIP (3-42), CRF, Urocortin, CGRP, AM and AM2 were purchased from Bachem (Bubendorf, Switzerland) and made to 1 mM stocks in water containing 0.1% BSA. Human glucagon, oxyntomodulin, and GLP-1 (7–36)NH_2_ were purchased from Alta Bioscience and prepared as 1 mM stocks in water containing 0.1% BSA.

### Generation of expression plasmids

Glucagon receptor, GLP-1R, FLAG-tagged RAMPs, HA-tagged CLR, CRF1βR and CRF2R were used as previously described^16,21,22,74^. The GIPR, GLP-2R, PTH1R, PTH2R and GHRHR constructs comprised the native signal peptide plus receptor sequence and were provided by Dr. Simon Dowell (GSK, Stevenage, UK). CTR was purchased from cDNA.org.

GIPR possess a putative N-terminal signal peptide that is cleaved during receptor processing and trafficking^75^. Therefore, to label the receptor at its N-terminus, a FLAG-tag was introduced immediately downstream of the predicted signal peptide. This was achieved using a previously described mutated version of pcDNA3.1^48^. Briefly, pcDNA3.1 was modified by the addition of a linker region encoding the influenza hemagglutinin signal peptide (MKTIIALSYIFCLVFAA) between the Kpn-1 and Not-1 sites of the multiple cloning site to produce pcDNA3.1-hgSP. The linker was constructed by annealing two complementary primers containing the hemagglutinin signal peptide sequence and Kpn-1 and Not-1 restriction sites. A FLAG-tag (DYKDDDDK) was introduced immediately downstream of the predicted signal peptide of GIPR by sequential overlapping PCR using primers, which also added a Not-1 and Xba-1 site to the product’s termini. This product was then ligated into pcDNA3.1-hsSP to produce FLAG-GIPR.

SNAP-RAMP constructs were generated via PCR amplification of RAMP1, 2, 3 or 3ΔPDZ DNA, without their native signal sequences, to introduce in-frame 5’ and 3’ restriction sites of EcoRI and EcoRV, respectively. RAMP PCR products were then ligated in frame into pcDNA3.1(+) containing sigSNAP. All SNAP-RAMP constructs were functional (Figure S2), although there was a change in rank potency of CLR agonists for the RAMP2 complex. However, RAMP1 and RAMP3 signalling were identical and SNAP-RAMP constructs were used solely for the purpose of BRET. Nluc-GPCR constructs were PCR amplified, without their native signal sequences, to introduce in-frame 5’ and 3’ BamHI and XbaI restriction sites, respectively. PCR products were then ligated in frame into pcDNA3.1(+) containing sigNLuc. FLAG-RAMP3ΔPDZ, HA-RAMP3ΔPDZ and SNAP-RAMP3ΔPDZ constructs were generated by removing the DTLL PDZ recognition sequence through site-directed mutagenesis (Agilent Technologies, Santa Clara, CA). Mutagenesis and SNAP-RAMP and Nluc-GPCR generation was confirmed by Sanger sequencing (Department of Biochemistry, University of Cambridge).

All GPCR-Rluc expression constructs were generated by ligation of class B GPCR cDNA (purchased from cDNA.org) into a CD33/Myc/RLuc backbone with cloning results confirmed by Sanger sequencing (Eton Biosciences). RAMP-YFP, β-arrestin-1/2-YFP and GRK5, expression plasmids were used as previously described^23^.

### Cell culture and transfection

HEK-293S cells (a gift from AstraZeneca), HEK-293T cells and HEK-293AΔβ-arrestin cells^76^ were cultured in Dulbecco’s Modified Eagle Medium (DMEM)/F12 supplemented with 10 % heat-inactivated fetal bovine serum (FBS, Sigma) and 1% antibiotic antimycotic solution (Sigma). HEK-293A cells were grown in DMEM supplemented with 10 % heat-inactivated FBS. The rank order of potencies was conserved across the three cell types (Figure S15), although absolute potencies were reduced in HEK293S by ∼3 fold (also previously demonstrated to express similar levels of RAMPs^77^), thus ensuring that it was possible to compare data between different HEK-293 cell lines. Suspension cells (HEK-293S) were therefore used for second messenger signalling assays and flow cytometry, HEK-293T cells were used for imaging and BRET assays due to their high transfection efficiency and adherent nature and, HEK-293A cells were used for NanoBiT assays as this technique was previously optimised in these cells^30^. All cell lines used were incubated at 37 °C in humidified 95 % air and 5 % CO_2_. For HEK-293S cells, transient transfections were performed using Fugene HD (Promega) in accordance with the manufacturer’s instructions using a 1:3 (w:v) ratio of DNA:Fugene HD. For HEK-293T and HEK-293AΔβ-arrestin cells, cells were transfected using polyethylenimine (PEI, Polysciences Inc.) and 150 mM NaCl using a 1:6 (w:v) ratio of DNA:PEI. For HEK-293A cells, transient transfections were performed using PEI Max (Polyscience Inc.) and 150 mM NaCl using a 1:6 (w:v) ratio of DNA:PEI Max. pcDNA3.1 was used throughout to maintain a consistent level of total DNA.

### cAMP Accumulation Assay

HEK-293S cells were transfected with GPCR and RAMP/pcDNA3.1 at a 1:1 ratio for 48 hours. Ligand-stimulated cAMP accumulation in the presence or absence of 0.5 mM 3-isobutyl-1-methylxanthine (IBMX) was measured after the indicated times of stimulation using LANCE® cAMP Detection Kit (Perkin Elmer Life Sciences) and a Mithras LB 940 multimode microplate reader, with 100 μM Forskolin (Sigma) used as a positive control as previously described^16,78^.

### Intracellular Calcium Mobilisation Assay

Mobilisation of intracellular calcium was measured in HEK-293S transfected with GIPR and FLAG-RAMP/pcDNA3.1 at a 1:1 ratio as previously decribed^16^. Ligands were robotically added using a BD Pathway 855 high-content bioimager and images were captured every second for 80 s. Fiji (Is Just) Image J was used to create a time series and to determine the intensity of the region of interest for the entire time course. Background fluorescence was corrected for and the maximum intensity used to generate concentration-response curves. 10 μM ionomycin (Cayman Biosciences) was used as a positive control.

### ERK1/2 Phosphorylation Assay

HEK-293S or HEK-293AΔβ-arrestin cells were transfected with GIPR and FLAG-RAMP/pcDNA3.1 at a 1:1 ratio. 48 hours post-transfection, cells were washed and resuspended in Ca^2+^ free HBSS. Cells were seeded at a density of 35000 per well in 384-well white Optiplates. To generate concentration-response curves ligands were added for 5 min, previously determined to be the optimum time for assaying acute ERK1/2 phosphorylation^13^. For time-course experiments, 100 nM GIP was added to the cells for the indicated times. Cells were then lysed using the supplied lysis buffer and assayed for ERK1/2 phosphorylation using the phospho-ERK (Thr202/Tyr204) Cellular Assay Kit (Cisbio). Plates were read using a Mithras LB 940 multimode microplate reader (Berthold Technologies) and 100 μM phorbol 12-myristate 13-acetate (PMA, Sigma) used as a positive control. Normalised dose-response data for GIP (1-42), GIP (D-Ala2) and GIP (Pro3) in cells expressing GIPR and pcDNA3.1/FLAG-RAMP3/ FLAG-RAMP3ΔPDZ were also fitted using the biphasic model in GraphPad Prism 8.4.2 (dashed lines).

### Chemical Inhibitors

Where appropriate, cells were treated with pertussis toxin (PTx, 200 ng/ml), for 16 h prior to assaying, to ADP-ribosylate Gα_i_, thereby uncoupling receptor-mediated Gα_i_-dependent inhibition of cAMP production^16^. To determine the contribution of Gα_q/11/14_ to signalling, cells were pretreated for 30 min, at room temperature, with 100 nM YM-254890 (Alpha Laboratories) to prevent GDP-GTP exchange at Gα_q/11/14_^79^ (100 nM is sufficient to completely block all specific signalling by G_q/11/14_^16^). To determine the contribution of different cAMP driven pathways to intracellular signalling, cells were pre-treated for 15 min, at room temperature, with 100 μM of the non-selective exchange factor directly activated by cAMP (EPAC1/2) inhibitor ESI-09 (Sigma)^80^, or 100 μM of the protein kinase A (PKA) inhibitor Rp-8-Br-cAMPs (Santa Cruz Biotechnology)^81^.

### RAMP-GPCR BRET screen

The BRET screen between GPCR-RLuc and RAMP-YFP was performed as previously described^23^. Briefly, HEK-293T cells transiently transfected with a constant concentration of GPCR-RLuc with increasing amounts of RAMP-YFP were cultured in 96-well, white, clear bottom plates coated with poly-D-lysine for 24 hours. Media was then replaced by 90 μL PBS containing 0.49 mM MgCl_2_.6H_2_O, 0.9 mM CaCl_2_.2H_2_O and the assay initiated by adding 10 μL of coelenterazine-h (Promega) to a final concentration of 5 μM. BRET readings were then measured 10 min after addition of coelenterazine h using a Mithras LB 940 multimode microplate reader. The acceptor/donor ratio (520 nm/460 nm) was calculated, and the data was fitted using either a linear regression or one-site binding (hyperbola) with GraphPad Prism 8.4.2.

### Flow cytometry screen for class B GPCR interactions with RAMPs

HEK-293S or HEK-293AΔβ-arrestin cells were transfected with GPCR and FLAG-RAMP/pcDNA3.1 (for RAMP surface expression) or FLAG-GIPR and HA-RAMP/pcDNA3.1 (for GIPR surface expression) at a 1:1 ratio. After 48 hours, 400000 cells were washed three times in FACS buffer (PBS supplemented with 1% BSA and 0.03% sodium azide) before and after 1 hour incubation at room temperature in 50 μL FACS buffer containing allophycocyanin (APC)-conjugated anti-FLAG monoclonal antibody (BioLegend, diluted 1:100 in FACS buffer). For internalisation and recycling experiments, cells were not treated, treated with 100 nM GIP (1-42) for 1 hour in complete DMEM/F12 at 37 °C, or treated, washed with PBS and incubated for 4 hours in complete DMEM/F12 containing 5 µg/ml cycloheximide (Sigma) to allow receptor recovery and prevent de novo protein synthesis^18^. Internalisation or recovery was stopped by washing with ice cold PBS and assayed as above but kept at 4°C throughout. To account for dead cells 2.5 μL propidium iodide (ThermoFisher Scientific) was added to each sample. Samples were analysed using a BD Accuri C6 flow cytometer, Ex. λ 633 nm and Em. λ 660 nm. GPCR interaction screen data were normalised to cell surface expression for cells co-transfected with HA-CLR and FLAG-RAMP2 as 100% and cells transfected with pcDNA3.1 as 0%. FLAG-GIPR surface expression was normalised to expression in the absence of HA-RAMP as 100% and pcDNA3.1 as 0%. For internalisation and recycling experiments, data were normalised to cell surface expression in the absence of treatment as 100% and vector control as 0%.

### Analysis of cell surface expression by enzyme-linked immunosorbent assay

HEK-293S cells were transfected with GPCR and FLAG-RAMP/pcDNA3.1 at a 1:1 ratio. Cell surface expression of FLAG-RAMPs was determined as previously described^22^. Briefly, 48 hours post-transfection, cells were fixed with 3.7% paraformaldehyde and washed 3 times with PBS before and after incubation with mouse anti-FLAG M2 primary antibody (Sigma, diluted 1:2000 in PBS with 1% BSA) or HRP-conjugated anti-mouse IgG secondary antibody (GE Healthcare, diluted 1:4000 in PBS with 1% BSA). Cells were then incubated in *o*-phenylenediamine (OPD) solution (SigmaFast *o*-phenylenediamine tablets, Sigma, dissolved in 20 ml distilled water) for 3-5 min, before termination of the reaction by addition to 100 μL 1 M sulphuric acid. Plates were read using a Mithras LB 940 multimode microplate reader at 492 nm. Data were normalised to cell surface expression for cells co-transfected with CLR and RAMP2 as 100% and cells transfected with pcDNA3.1 as 0%.

### Cell Surface BRET

HEK-293S cells were transfected with 100 ng Nluc-GPCR and various amounts of each SNAP-RAMP (0, 5, 10, 25, 50, 75, 100, 200 or 500 ng). 24 hours post-transfection, cells were incubated with 200 nM SNAP-Surface® Alexa Fluor® 488 (New England Biolabs, UK, diluted in serum-free DMEM/F12), for 30 min at 37°C with 5% CO_2_. Cells were then washed three times with KREBS buffer (126 mM NaCl, 2.5 mM KCl, 25mM NaHCO_3_, 1.2 mM NaH_2_PO_4_, 1.2 mM MgCl_2_, 2.5 mM CaCl_2_), before being harvested and resuspended in KREBS buffer. Cells were seeded at 20000 cells per well in a white 96-well plate (ThermoFisher Scientific). Nano-Glo® Live Cell Substrate (Promega) was added to each well and BRET readings (460 nm and 515 nm) were measured after 10 min. The acceptor/donor ratio is shown relative to the transfected DNA ratios and data fitted using either a linear regression or one-site (hyperbola) with GraphPad Prism 8.4.2.

### Cell Surface BRET Imaging

120000 HEK-293S cells were seeded onto poly-D-lysine-coated 35mm 4-chamber MatTek dishes (Ashland) prior to transfection. After 24 hours, cells were transfected with Nluc-CLR or Nluc-GIPR and each SNAP-RAMP/pcDNA3.1. 24 hours after transfection, cells were labelled by replacing complete growth medium with 500 μl serum free DMEM/F12 containing 200 nM SNAP Surface Alexa Fluor-488 (New England Biolabs) and were incubated for 30 min at 37°C with 5% CO_2_. Before imaging, cells were washed and incubated with 500 μl HBSS supplemented with 1.8 g/L glucose. RAMP expression and localisation was visualised by imaging fluorescence through a fluorescein isothiocyanate (FITC) channel (1 second exposure, 488/10 nm excitation, 583/22 nm emission). Cells were then incubated with Nano-Glo® Substrate (Promega) to a final concentration of 10 μM, for 15 min, and bioluminescence and BRET images were subsequently captured using an open channel (2 second exposure) and FITC channel (10 second exposure, 509/22 nm emission), respectively. Bioluminescence imaging was performed using an Olympus LV200 microscope, equipped with a 60x oil immersion objective lens. BRET ratio measurements, using membrane-localised fluorescence and bioluminescence signals, were performed using Fiji (Is Just) Image J.

### NanoBiT G protein activation assay

HEK-293A cells were transiently transfected with GIPR, appropriate Gα-LgBiT and Gβ subunits^30^, Gγ_2_-SmBiT and FLAG-RAMP/pcDNA3.1 at a 2:1:3:3:2. For G_q_, G_11_, G_14_ and G_15_, cells were also transfected with RIC8A, a chaperone protein required for Gq family signalling^82^, at a 1:1 ratio to GIPR. The optimal Gβ subunit used for each Gαγ_2_ are as follows; Gα_s_, Gβ_1_; Gα_i2_, Gβ_1_; Gα_i3_, Gβ_1_; Gα_z_, Gβ_1_; Gα_q_, Gβ_1_; Gα_11_, Gβ_3_; Gα_14_, Gβ_2_; Gα_15_, Gβ_2_; Gα_12_, Gβ_1_; Gα_13_, Gβ_3_. 24 hours after transfection, cells were harvested and seeded at 60000 cells per well into poly-D-lysine-coated clear-bottomed 96 well plates (Corning) and cultured for a further 24 hours. Media was then removed, and cells were washed with HBSS plus 10 mM HEPES before addition of 80 μl HBSS containing 10 mM HEPES and 0.1 % BSA. 10 μl of coelenterazine-h (diluted in HBSS containing 10 mM HEPES and 0.1 % BSA) was then added to each well to a final concentration of 5 μM, and the plate incubated for 1 hour in the dark. After incubation, a baseline was established for 2 min before ligands were robotically added using a Hamamatsu Functional Drug Screening System (FDSS) and luminescence measured every 10 seconds for 10 min. GIP (1-42) and GIP (D-Ala2) were added in a log dilution series between 1 μM and 0.1 pM, whilst GIP (Pro3) was added in a 0.5 log dilution series between 1 μM and 1 nM. Ligand-induced change in luminescent units were corrected to baseline and vehicle, and the area under the curve (AUC), for the entire timecourse, used to generate concentration-response curves. Data were normalised to the maximal response, determined by fitting to the three-parameter logistic model, for all data where E_max_ could be determined robustly. Where this was not possible the response to 1 μM ligand was used. For G_14_, raw AUC data was collated due to the variability of the data. Where appropriate, normalised dose-response data were also fitted using the biphasic model in GraphPad Prism 8.4.2 (dashed lines).

### Internalisation and resensitisation imaging

The cellular localisation of GIPR and RAMPs was determined as described^23^. Briefly, HEK-293T cells transiently transfected with the indicated combination of GIPR ± RAMP were treated with or without 100 nM GIP (1-42) for 1 hour in complete DMEM/F12 at 37 °C. For receptor resensitisation after agonist stimulation, cells were washed with PBS and incubated for 4 hours in complete DMEM/F12 containing 5 μg/ml cycloheximide. Cells were then fixed in 4% paraformaldehyde, blocked in PBS + 4% BSA, incubated with appropriate primary and secondary antibodies and images visualised and processed as previously described^23^. Pearson’s correlation coefficients (r) were determined using the colocalization threshold plugin for ImageJ as described in Weston et al., 2014^83^. Four separate Regions of Interest (ROI) were selected and mean ± SD was determined.

### β-Arrestin recruitment

HEK-293T cells were transfected with myc-GIPR-Rluc, β-arrestin-1/2-YFP, GRK5 and FLAG-RAMP/pcDNA3.1 at a 1:5:4:1 ratio and grown overnight. 150000 cells were seeded into poly-L-lysine coated 96-well plates (Perkin Elmer) in reduced serum media (MEM + 2% FBS + 1% antibiotic antimycotic solution). The following day, β-arrestin-1/2 recruitment was measured after 6 min (determined from initial 60 min timecourse experiments as the time point at which β-arrestin-1/2 recruitment was maximal) or 60 min (to assess long term β-arrestin-1/2 recruitment) using a Berthold Mithras LB 940 multimode microplate reader as previously described^84^. The GIP (1-42)-induced change in 530 nm/485 nm) ratio was corrected to vehicle treated cells. Data were converted to milliBRET (mBRET) units by multiplying by 1000.

### In situ hybridisation (RNAscope) and immunohistochemistry

Pancreatic tissue was collected from six wildtype 129/S6-SvEv-TC1 mice and fixed in 4% PFA at 4°C overnight. The tissue was removed from PFA and washed in PBS before dehydrating and embedding in paraffin. Paraffin sections were cut to 5 μm thickness. RNAscope *in situ* hybridisation to detect RNA transcripts was performed according to the manufacturer protocol (Advanced Cell Diagnostics) using probes for RAMP1 (#532681) and RAMP3 (#497131). Antibody staining was performed following *in situ* hybridisation with guinea pig anti-insulin antibody (Invitrogen, diluted 1:1000) and rabbit anti-glucagon antibody (ZYMED, diluted 1:300) in 3% BSA in PBS with 0.1% Triton X overnight at 4°C. This was followed by secondary antibody staining with goat anti-guinea pig IgG (Jackson ImmunoResearch, diluted 1:400) and donkey anti-rabbit IgG (Jackson ImmunoResearch, diluted 1:200), respectively, for 1 hour at room temperature.

### Blood glucose and insulin measurements

All mice used in this study were between 8 to 20 weeks of age and are of the 129/S6-SvEv-TC1 background. The generation of *Ramp1* and *Ramp3* knockout mice were previously described^38,39^. All animal experiments were approved by the Institutional Animal Care and Use Committee of the University of North Carolina at Chapel Hill. This study was powered to attain statistical significance of p < 0.05 with a 90% probability between RAMP1^+/+^ and RAMP1^−/−^ and RAMP3^+/+^ and RAMP3^−/−^ mice. Male and female mice were fasted for 4 hours (9am to 1pm) with free access to water prior to baseline blood collection and treatment. Mice we treated by intraperitoneal injection with glucose (1g/kg, Sigma #G5767) containing GIP (D-Ala2) (50 nmol/kg, Tocris #6699) in sterile saline. Fasting blood was collected by submandibular bleed immediately before treatment and 20 min after treatment. Blood glucose was measured with a glucometer immediately upon collection and serum was isolated and stored at −80°C until insulin was measured by mouse insulin ELISA (Alpco) according to the manufacturer protocol.

### Data Analysis

Data analyses for all assays were performed in GraphPad Prism 8.4.2 (San Diego, CA, USA). For cAMP accumulation, [Ca^2+^]_i_ mobilisation and ERK1/2 phosphorylation assays, data were fitted to obtain concentration–response curves using a three-parameter logistic equation, to obtain values for pEC_50_ (-logEC_50_) and E_max_ (as a percentage of forskolin, ionomycin and PMA stimulation, respectively). Where data are expressed relative to the maximal response, data were normalised to the E_max_ determined from the three-parameter logistic fit. Where dose-response curves clearly displayed both a high potency and low potency phase, data were also fitted to the biphasic model and are displayed as dashed lines. Values obtained from biphasic fits are quoted in the text and were used to guide interpretation of results, whilst tables were populated with and data analysed using the values obtained from the three-parameter fit. To assess whether there was any ligand bias for GIP (1-42), GIP (D-Ala2) or GIP (Pro3) for each intracellular signalling pathway and activation of each subclass of G protein, Log intrinsic relative activity (LogRA_i_)^85,86^ were calculated relative to GIP (1-42), or GIPR expressed alone, respectively using the following equation;

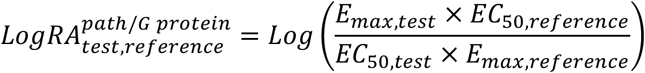

Statistical differences were analysed using one-way ANOVA and Dunnett’s post-hoc test, Student’s t-test or a Kruskal-Wallis test, as appropriate. A probability of p<0.05 was considered significant, values are stated as mean ± standard error of the mean (SEM).

## Author Contributions

GL, PMS, DRP, DW, PZ, KMC conceived and designed the research; MH performed second messenger signalling, β-arrestin recruitment and ligand binding assays; DIM performed BRET screen and receptor trafficking imaging, MTH, DS and MH performed FACS analysis and analysed data, SR performed preliminary experiments, MH, HYY, TT, PZ, performed the mammalian signalling assays, G protein activation; SAS, BAZ, AI and SC generated constructs, SJH, SJB, MS advised on BRET imaging, SC and MS performed cell surface BRET assays and imaging, JBP performed the mouse experiments, MH and GL wrote the manuscript, SJH, SJB, PMS, DW, PZ, KMC and DRP revised and edited the manuscript.

## Acknowledgements

Authors acknowledge the support of BBSRC Doctoral Training Partnership BB/JO1454/1 (MH) and BBSRC (grant number BB/M000176/2) awarded to GL and DRP, Rosetrees foundation (to HYY and GL), Endowment Fund for education from Ministry of Finance Republic of Indonesia (DS), the National Health and Medical Research Council of Australia (NHMRC) project grants, 1120919 (PMS) and 1126857 (DW), the US National Institutes of Health grants, NHLBI HL129086, NIDDK DK119145 (to KMC), NHLBI HL134279 (to DIM) and NICHD HD095585 (to JBP) and the PRIME 19gm5910013, the LEAP JP19gm0010004 and the BINDS JP19am0101095 from the Japan Agency for Medical Research and Development (AMED) and the Uehara Memorial Foundation (to AI). HYY is also supported by an international scholarship from the Cambridge Trust. PMS is an NHMRC Senior Principal Research Fellow and DW is an NHMRC Senior Research Fellow.

