## Supplementary material for "RAMPs regulate signalling bias and internalisation of the GIPR": supp info

**Address for correspondence:**

Supplementary Figures S1-13

Supplementary Tables S1-S8

**Supplementary Figure 1**

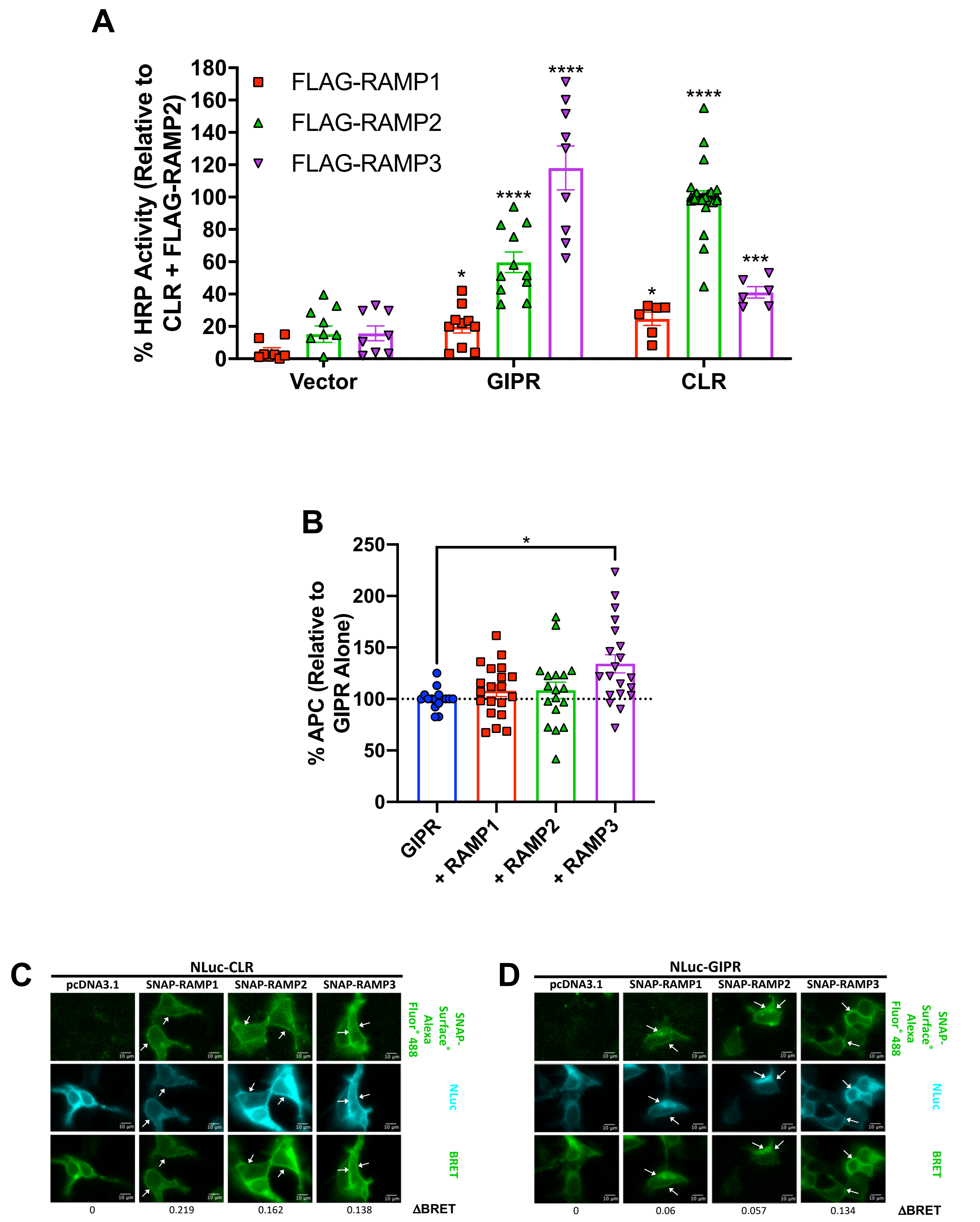


**Figure S1. Cell surface ELISA and plasma membrane expression of FLAG-GIPR.**
*A*. HEK-293S cells were cotransfected with each FLAG-RAMP and one of pcDNA3.1, GIPR or HA-CLR, at a 1:1 ratio. Plasma membrane (PM) expression of FLAG-RAMPs was determined by enzyme-linked immunosorbent assay using an HRP-tagged anti-FLAG monoclonal antibody. An increase in FLAG-RAMP PM expression was indicated by an increase in signal at 492 nm. Surface expression was normalised to FLAG-RAMP2 when cotransfected with HA-CLR as 100 % and pcDNA3.1 (as 0%). Endogenous surface expression of FLAG-RAMPs was determined by

cotransfection with pcDNA3.1. All values are the mean ± S.E.M. Data were assessed for statistical differences, at $p<0.05$, in cell surface FLAG-RAMP expression compared to expression in the absence of receptor using a one-way ANOVA with Dunnett's post-hoc test (*, $p<0.05$; ***, $p<0.001$; ****, $p<0.0001$). *B*. HEK-293S cells were cotransfected with FLAG-GIPR and one of pcDNA3.1, HA-RAMP1, HA-RAMP2 or HA-RAMP3 at a 1:1 ratio. PM expression of FLAG-GIPR was determined by flow cytometry using an APC-conjugated anti-FLAG monoclonal antibody. Surface expression was normalised to that of FLAG-GIPR alone as 100 % and pcDNA3.1-zeo as 0%. All values are the mean ± S.E.M of at least 11 individual data sets. Data were assessed for statistical differences, at $p<0.05$, in cell surface FLAG-GIPR expression compared to expression in the absence of RAMP using a one-way ANOVA with Dunnett's post-hoc test (*, $p<0.05$). *C-D*. Representative live cell imaging of BRET interactions between SNAP-RAMPs coexpressed with either Nluc-CLR (*C*) or Nluc-GIPR (*D*) in HEK-293S cells. Arrows indicate PM localisation. BRET values were obtained by dividing the signal obtained for SNAP Surface® Alexa Fluor® 488 by the Nluc signal with ΔBRET values for the images displayed.

### Supplementary Figure 2

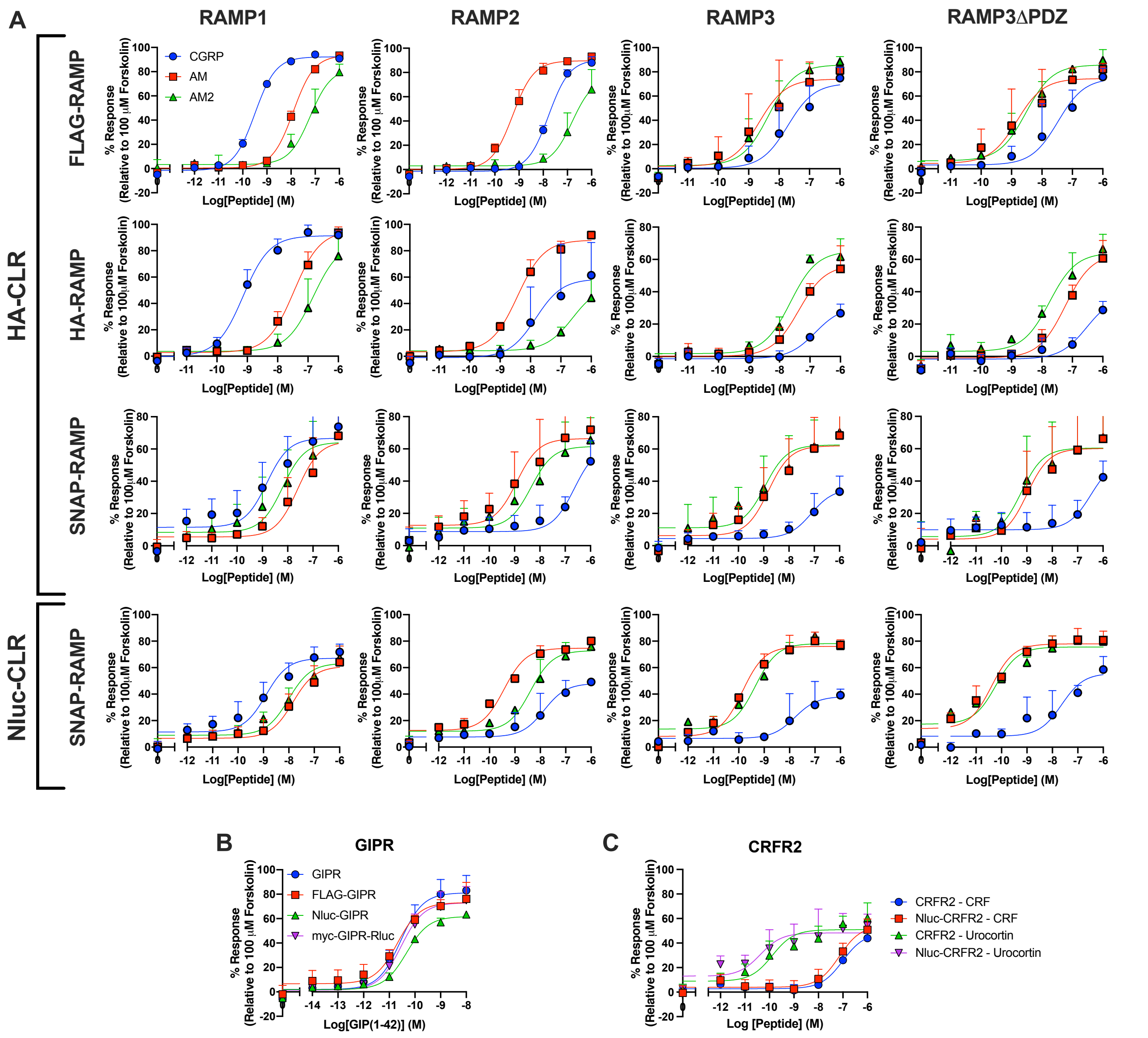


**Figure S2. Verification that all newly generated expression constructs were functional.** *A*. cAMP accumulation was determined in HEK-293S cells transfected with either HA-CLR or Nluc-CLR and either FLAG, HA or SNAP-tagged RAMPs (including RAMP3ΔPDZ) following 30 min stimulation with calcitonin gene-related peptide (CGRP; blue circles), adrenomedullin (AM; red squares) or adrenomedullin 2 (AM2; green triangles). *B*. cAMP accumulation was determined in cells transfected with one of untagged GIPR, FLAG-GIPR, Nluc-GIPR or myc-GIPR-Rluc following 8 min stimulation with GIP (1-42). *C*. cAMP accumulation was determined in HEK-293S cells transfected with either untagged CRFR2 or Nluc-CRFR2 following 8 min stimulation with CRF or urocortin. Data are expressed relative to 100 μM forskolin and are the mean + SD of 2-5 individual data sets.

**Supplementary Figure 3**

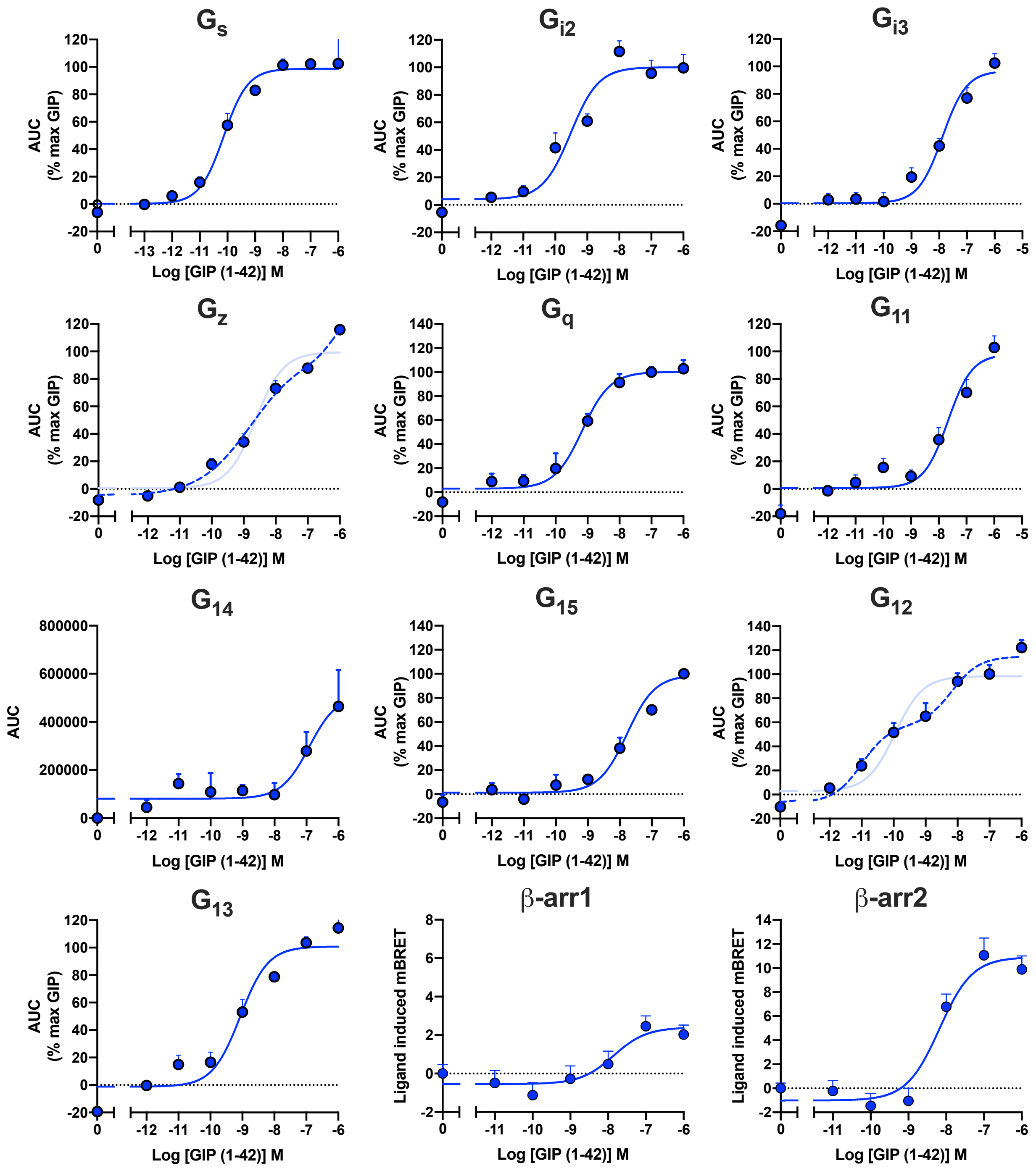


**Figure S3. Concentration-response curves for activation of each G protein in response to GIP (1-42).**

HEK-293A cells were cotransfected with GIPR, appropriate Gα-LgBiT and Gβ subunits, $G\gamma_2$-SmBiT and pcDNA3.1 at a 2:1:3:3:2 ratio. RIC8A was also included for $G_q$, $G_{11}$, $G_{14}$ and $G_{15}$. G protein activation was measured by the GIP (1-42)-induced change in RLU. Data points were corrected to baseline and vehicle and the AUC used to produce the concentration-response curves shown for each G protein. Data were normalised to the maximum response to GIP (1-42), determined by three-parameter logistic fits, for each condition and are expressed as mean + SEM of 3-7 individual experiments. Biphasic fits for $G_z$ and $G_{12}$ are displayed as dashed lines, with the equivalent three parameter fits faded. Peak β-arrestin-1/2 recruitment was measured in HEK-293T cells transiently expressing GIPR-Rluc, GRK5, each FLAG-RAMP/pcDNA3.1 and β-arrestin-1/2-YFP after 6 min stimulation with GIP (1-42). Data are expressed as ligand-induced delta milli BRET (mBRET) and are the mean + SEM of 3-8 individual experiments.

### Supplementary Figure 4

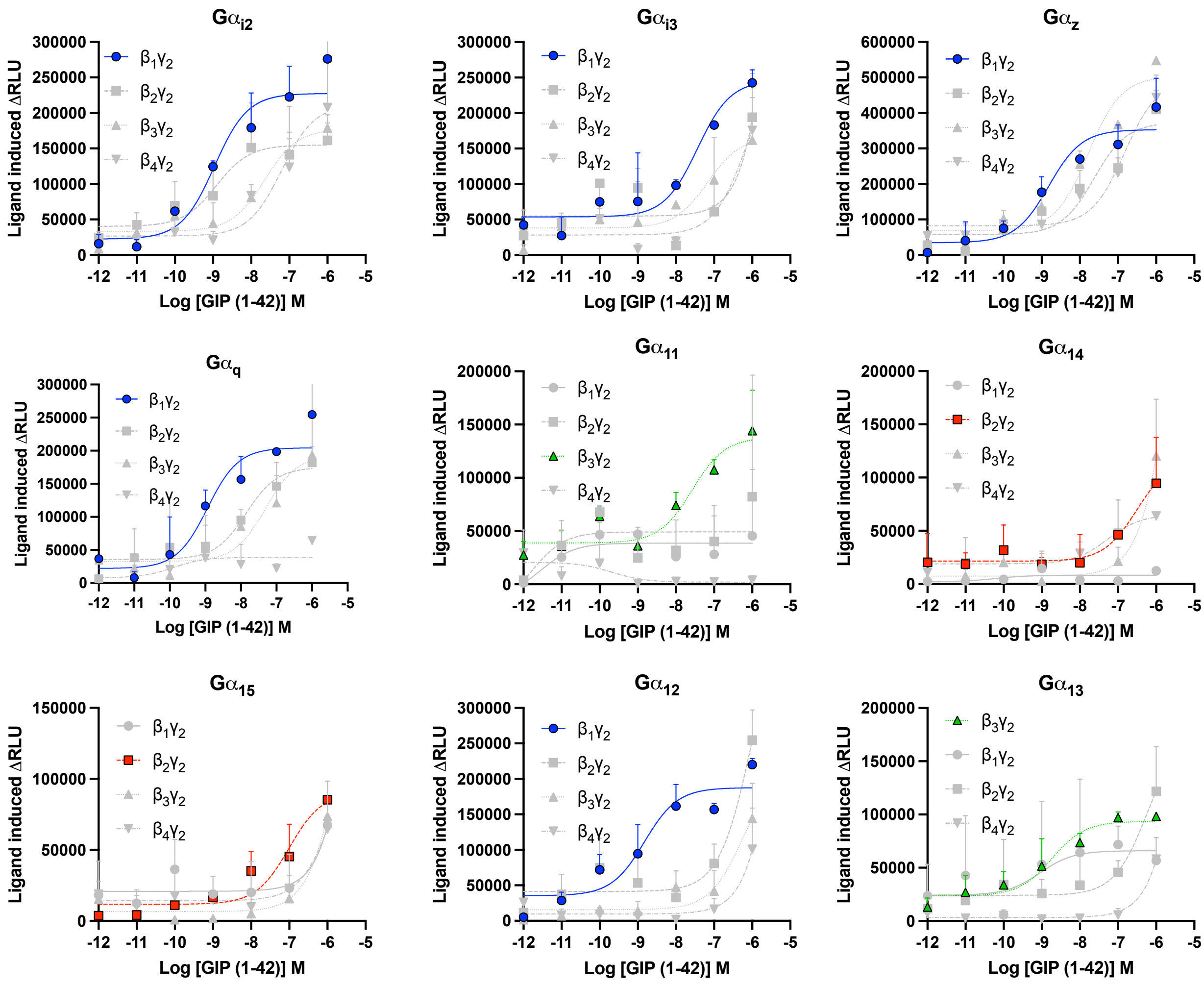


**Figure S4. Optimization of Gβγ combination**

HEK-293A cells were cotransfected with GIPR, appropriate Gα-LgBiT, Gγ2-SmBiT, pcDNA3.1 and each Gβ subunit, at a 2:1:3:2:3 ratio. RIC8A was also included for $G_q$, $G_{11}$, $G_{14}$ and $G_{15}$. G protein activation was measured by the GIP (1-42)-induced change in RLU. Data points were corrected to baseline and vehicle and the AUC used to produce the concentration-response curves shown for each G protein. The optimal Gβγ combination for each Gα-LgBiT is displayed in colour. Data are the mean + SD of 2-3 individual experiments performed in triplicate.

**Supplementary Figure 5**

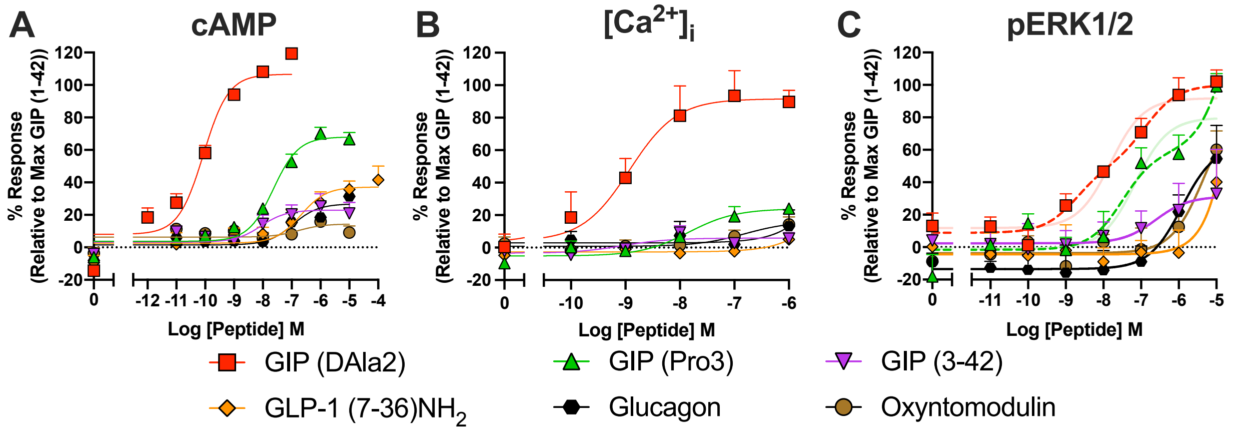


**Figure S5. Signalling profile of GIP-based and glucagon family ligands**

*A-C.* Cells were transiently transfected with GIPR with cAMP accumulation (*A*), $[Ca^{2+}]_i$ mobilisation (*B*) and ERK1/2 phosphorylation (*C*) measured as above in response to stimulation with other GIP-based (GIP (D-Ala2), GIP (Pro3) and GIP (3-42)) and glucagon family ligands (glucagon, GLP-1 (7-36)$NH_2$, oxyntomodulin and liraglutide). Data are expressed relative to the maximal GIP (1-42) response, determined by three-parameter logistic fits and are mean + SEM of 3-10 individual experiments with quantitative data displayed in (Table 1). The dashed line represents the biphasic fits for GIP (D-Ala2) and GIP (Pro3) pERK1/2 responses, with the equivalent three parameter fits faded.

### Supplementary Figure 6

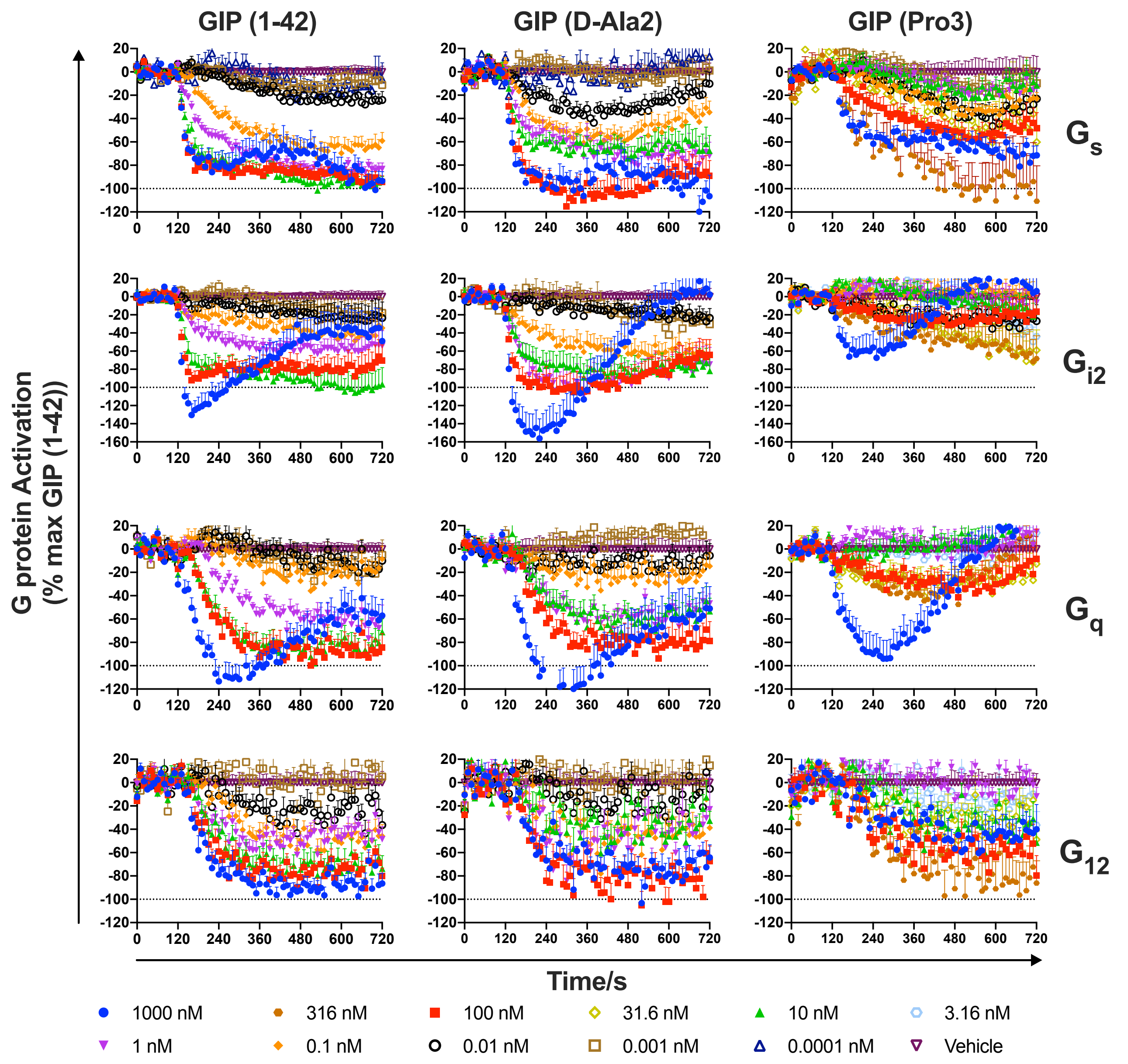


**Figure S6. Activation of G$\alpha_s$, G$\alpha_{i2}$, G$\alpha_q$ G$\alpha_{12}$ in response to GIP (1-42), GIP (D-Ala2) or GIP (Pro3).**

HEK-293A cells were cotransfected with GIPR, one of G$\alpha_s$-LgBiT, G$\alpha_{i2}$-LgBiT, G$\alpha_q$-LgBiT or G$\alpha_{12}$-LgBiT, G$\beta_1$, G$\gamma_2$-SmBiT, and pcDNA3.1 at a 2:1:3:3:2 ratio. RIC8A was also included for $G_q$. G protein activation is expressed as percentage agonist-induced G protein dissociation. G protein activation was measured by the agonist-induced change in RLU and data are expressed as a percentage of G protein activation induced by GIP (1-42) for each G protein. Data are expressed as mean + SEM of 4-7 individual experiments with quantitative data displayed in (Table S2).

### Supplementary Figure 7

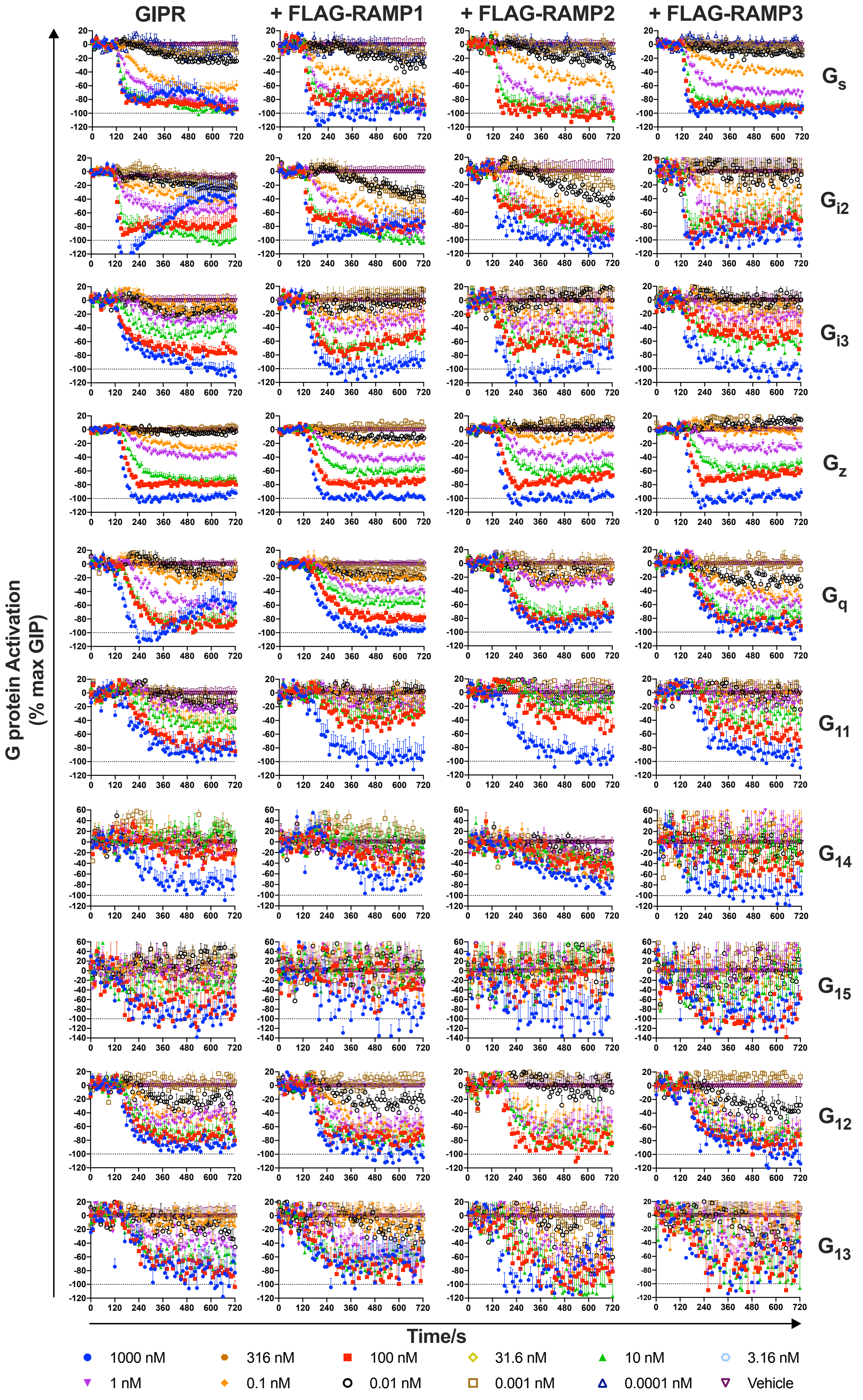

**Figure S7. Traces for G protein activation for each G protein with each RAMP for GIP (1-42).** HEK-293A cells were cotransfected with GIPR, appropriate $G\alpha$-LgBiT and $G\beta$ subunits, $G\gamma_2$-SmBiT and FLAG-RAMP/pcDNA3.1 at a 2:1:3:3:2 ratio. RIC8A was also included for $G_q$, $G_{11}$, $G_{14}$ and $G_{15}$. G protein activation was measured by the agonist-induced change in RLU and data were normalised to the maximum response to GIP (1-42) for each condition. Data are the mean + SEM of 3-7 individual experiments.

**Supplementary Figure 8**

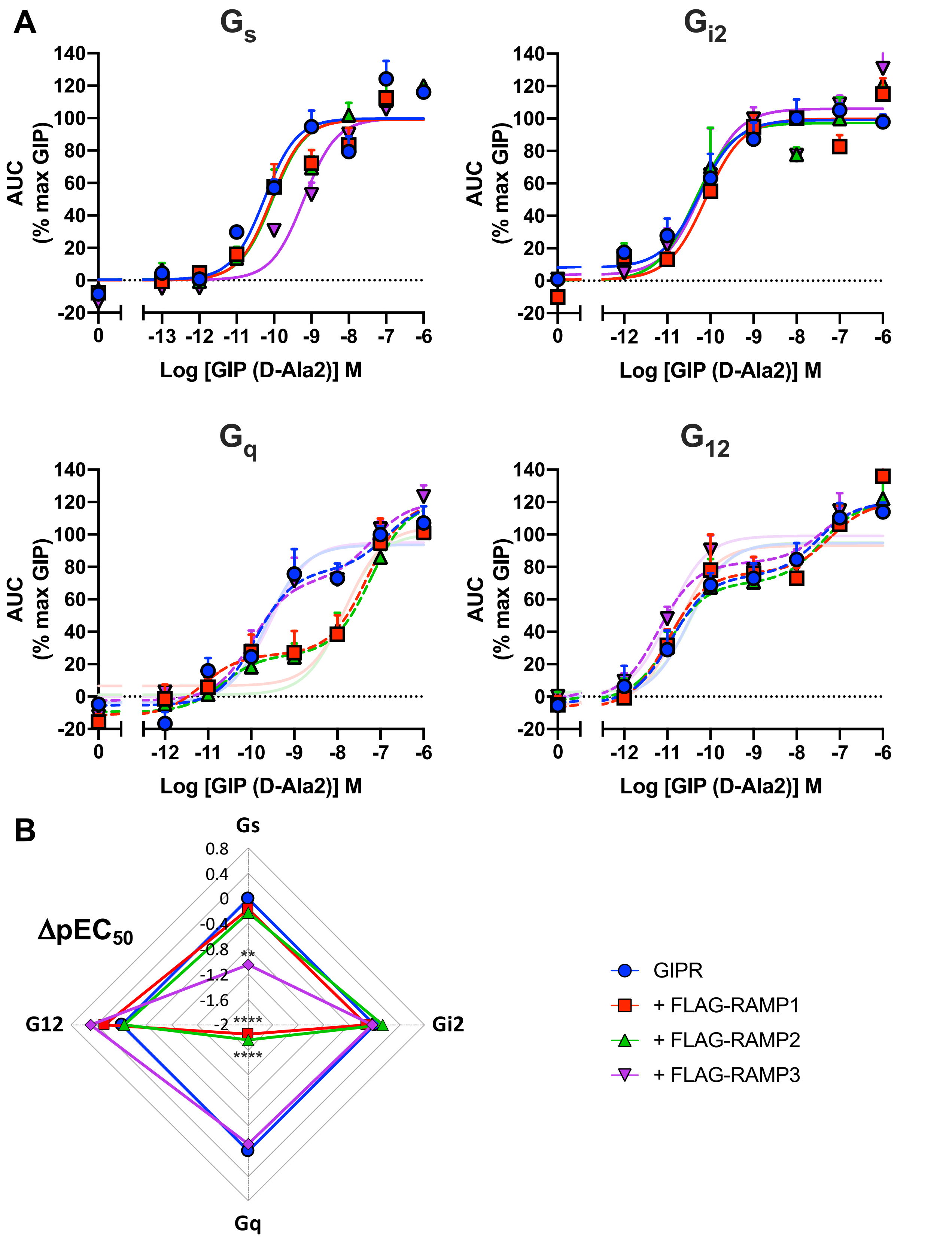


**Figure S8. Activation of Gα$_s$, Gα$_{i2}$, Gα$_q$ Gα$_{12}$ in response to GIP (D-Ala2).**

*A*. HEK-293A cells were cotransfected with GIPR, one of Gα$_s$-LgBiT, Gα$_{i2}$-LgBiT, Gα$_q$-LgBiT or Gα$_{12}$-LgBiT, Gβ$_1$, Gγ$_2$-SmBiT and FLAG-RAMP/pcDNA3.1 at a 2:1:3:3:2 ratio. RIC8A was also included for G$_q$. G protein activation was measured by the GIP (1-42)-induced change in RLU. Data points were corrected to baseline and vehicle and the AUC used to produce the concentration-response curves shown for each G protein. Data were normalised to the maximum response to GIP (D-Ala2) for each condition and are the mean + SEM of 3-6 individual experiments with quantitative data displayed in (Table S4). *B*. Radial plot showing the change in log potency of G protein activation induced by coexpression with each FLAG-RAMP relative to GIPR + pcDNA3.1. Data were assessed for statistical differences, at $p<0.05$, in the change in $pEC_{50}$ compared to GIPR expressed with pcDNA3.1 using a one-way ANOVA with Dunnett's post-hoc test (**, $p<0.01$; ****, $p<0.0001$). Biphasic fits for $G_q$ and $G_{12}$ are displayed as dashed lines, with equivalent three-parameter fits faded.

### Supplementary Figure 9

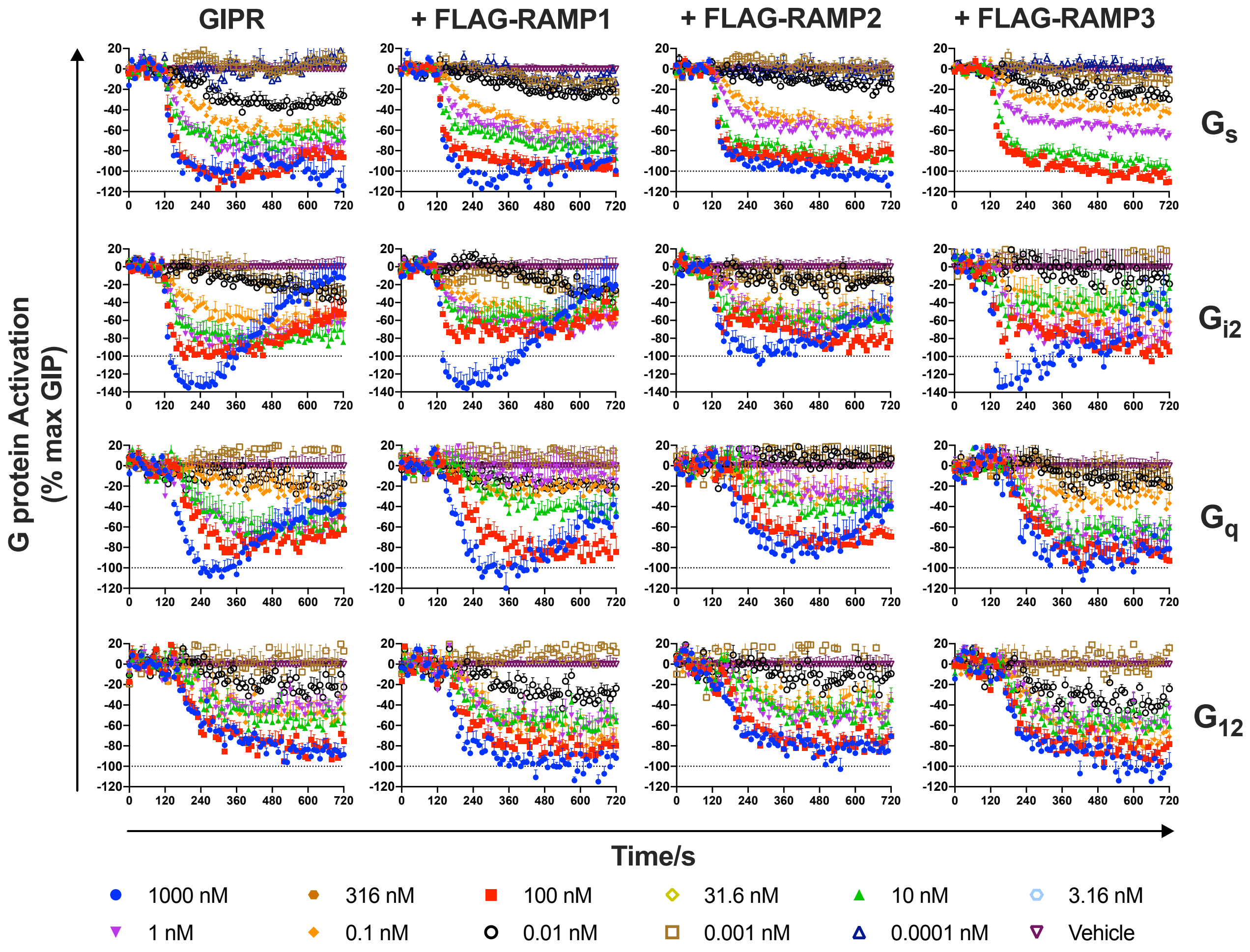


**Figure S9. Traces for G protein activation for each G protein with each RAMP for GIP (D-Ala2).** HEK-293A cells were cotransfected with GIPR, one of G$\alpha_s$-LgBiT, G$\alpha_{i2}$-LgBiT, G$\alpha_q$-LgBiT or G$\alpha_{12}$-LgBiT, G$\beta_1$, G$\gamma_2$-SmBiT and FLAG-RAMP/pcDNA3.1 at a 2:1:3:3:2 ratio. RIC8A was also included for G$_q$. G protein activation was measured by the agonist-induced change in RLU and data were normalised to the maximum response to GIP (D-Ala2) for each condition. Data are expressed as mean + SEM of 3-6 individual experiments with quantitative data displayed in (Table S4).

**Supplementary Figure 10**

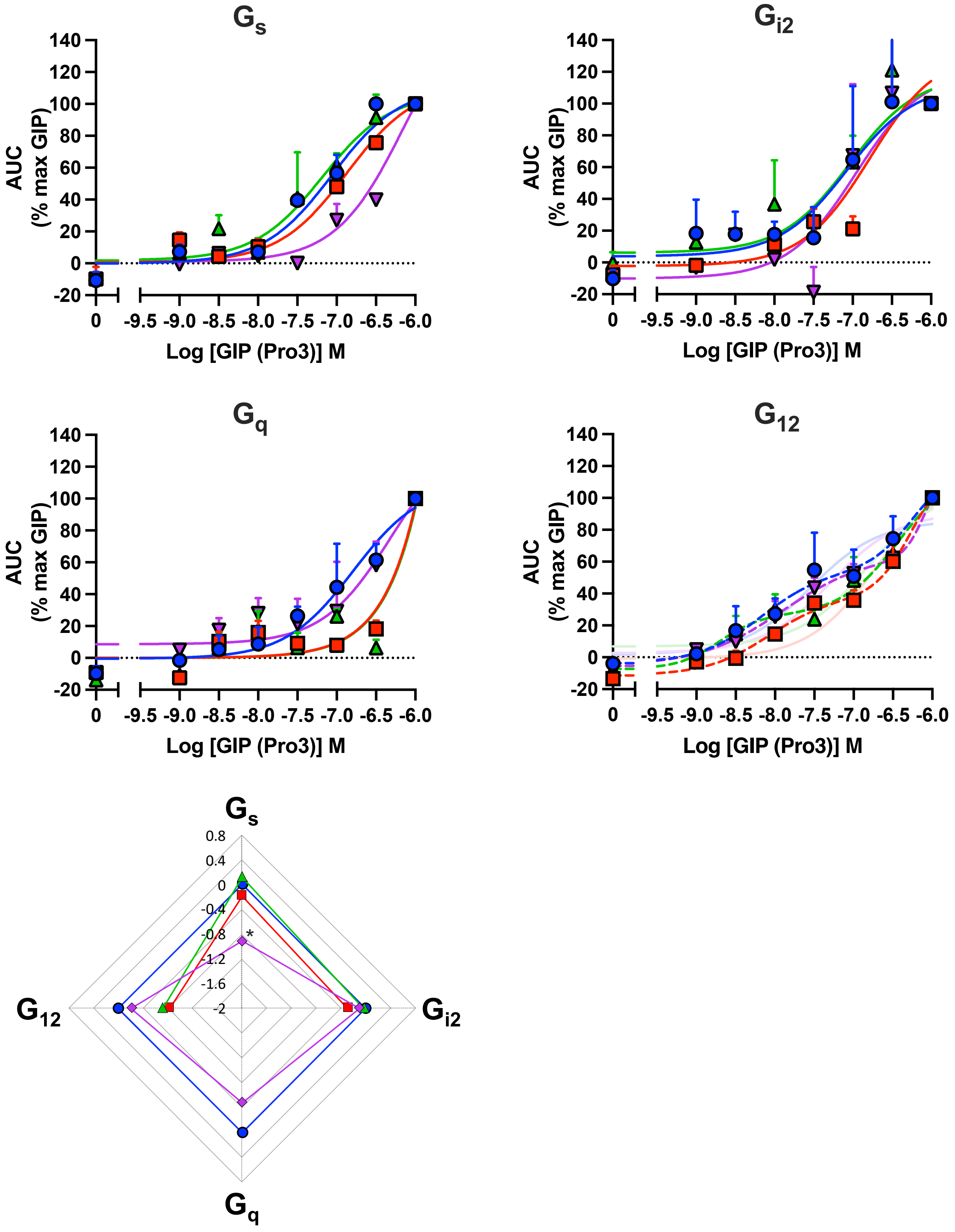


**Figure S10. Activation of Gα$_s$, Gα$_{i2}$, Gα$_q$ Gα$_{12}$ in response to GIP (Pro3).**

*A*. HEK-293A cells were cotransfected with GIPR, one of Gα$_s$-LgBiT, Gα$_{i2}$-LgBiT, Gα$_q$-LgBiT or Gα$_{12}$-LgBiT, Gβ$_1$, Gγ$_2$-SmBiT and FLAG-RAMP/pcDNA3.1 at a 2:1:3:3:2 ratio. RIC8A was also included for G$_q$. G protein activation was measured by the GIP (1-42)-induced change in RLU. Data points were corrected to baseline and vehicle and the AUC used to produce the concentration-response curves shown for each G protein. Data were normalised to the maximum response to GIP (Pro3) for each condition and are the mean + SEM of 3-6 individual experiments with quantitative data displayed in (Table S5). *B*. Radial plot showing the change in log potency of G protein activation induced by coexpression with each FLAG-RAMP relative to GIPR + pcDNA3.1. Data were assessed for statistical differences, at $p<0.05$, in the change in $pEC_{50}$ compared to GIPR expressed with pcDNA3.1 using a one-way ANOVA with Dunnett's post-hoc test (*, $p<0.05$). Biphasic fits for $G_{12}$ are displayed as dashed lines, with equivalent three-parameter fits faded.

**Supplementary Figure 11**

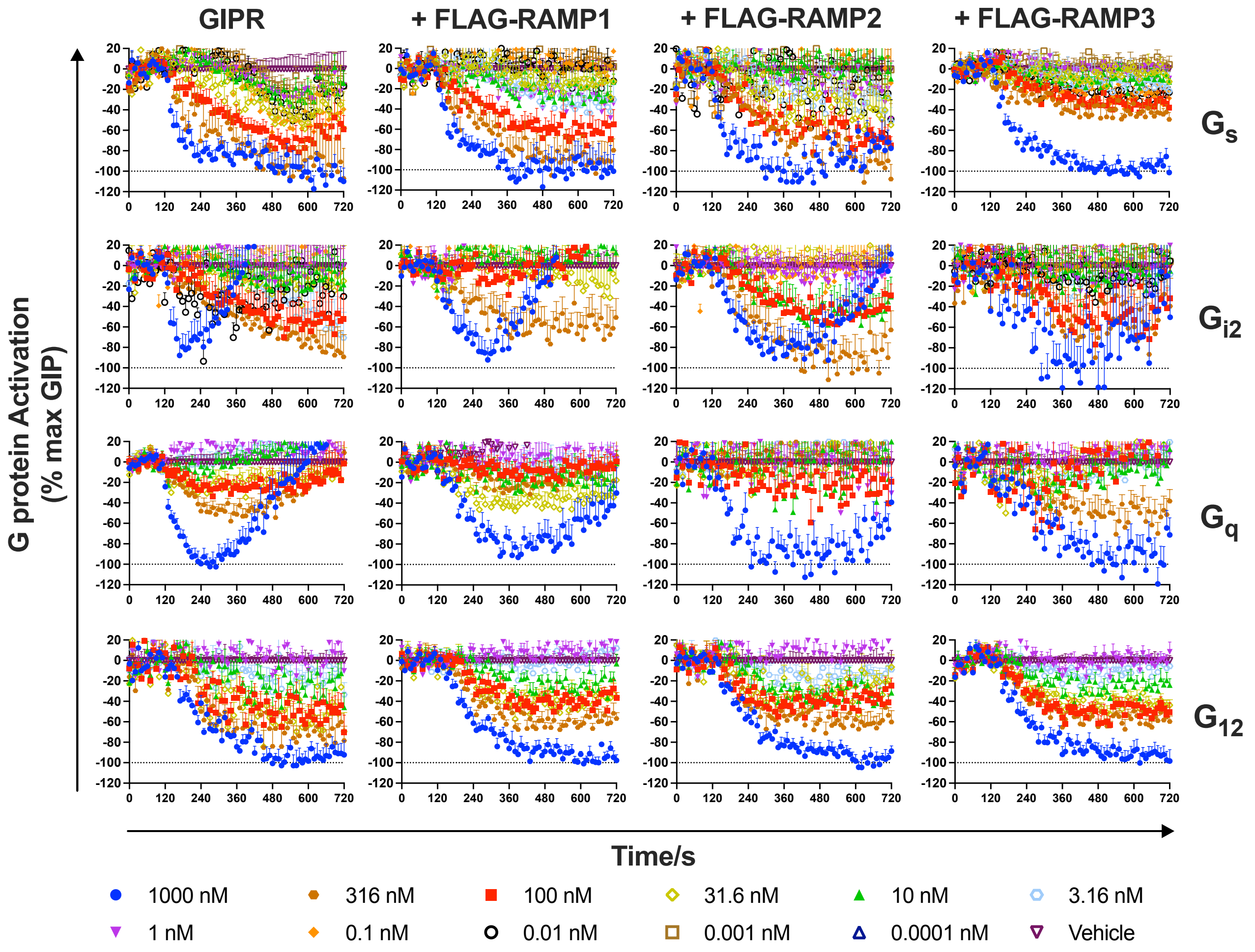


**Figure S11. Traces for G protein activation for each G protein with each RAMP for GIP (D-Pro3).**

HEK-293A cells were cotransfected with GIPR, one of $G\alpha_s$-LgBiT, $G\alpha_{i2}$-LgBiT, $G\alpha_q$-LgBiT or $G\alpha_{12}$-LgBiT, $G\beta_1$, $G\gamma_2$-SmBiT and FLAG-RAMP/pcDNA3.1 at a 2:1:3:3:2 ratio. RIC8A was also included for $G_q$. G protein activation was measured by the agonist-induced change in RLU and data were normalised to the maximum response to GIP (Pro3) for each condition. Data are expressed as mean + SEM of 3-6 individual experiments with quantitative data displayed in (Table S5).

**Supplementary Figure 12**

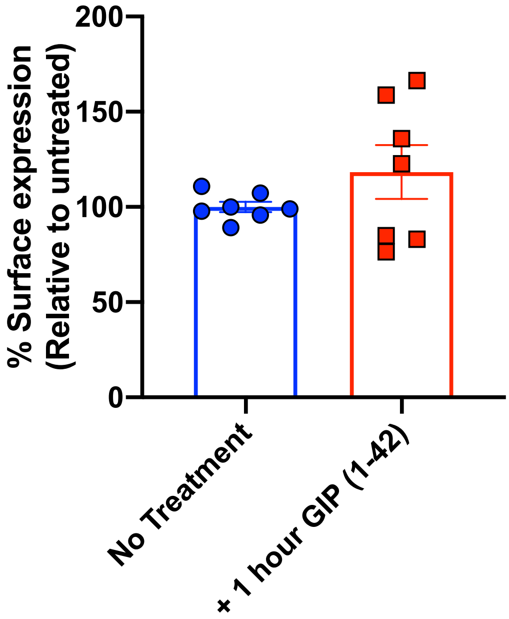


**Figure S12. Treatment with GIP (1-42) reduces PM expression of FLAG-GIPR in a β-arrestin-dependent manner.**

HEK-293AΔβ-arrestin cells were cotransfected with FLAG-GIPR and pcDNA3.1 at a 1:1 ratio. Cells were either treated with 100 nM GIP (1-42) for 1 hour, or treated, washed and allowed to recover for 4 hours, in the presence of cycloheximide. PM expression of FLAG-GIPR was determined by flow cytometry using an APC-conjugated anti-FLAG monoclonal antibody. Surface expression was normalised to the level observed in the absence of treatment (as 100%) and pcDNA3.1 (as 0%). All values are the mean ± S.E.M and were assessed for statistical differences, at $p<0.05$, in the change in FLAG-GIPR surface expression compared to expression in the absence of treatment using a Kruskal-Wallis test (**, $p<0.01$; ***, $p<0.001$).

**Supplementary Figure 13**

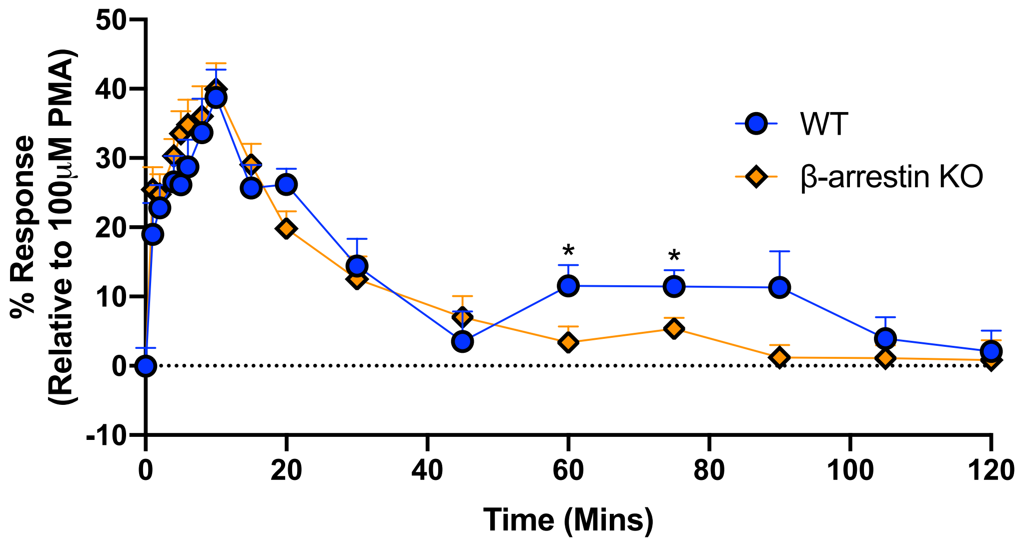


**Figure S13. The second phase of ERK1/2 signalling is β-arrestin-dependent**
Temporal ERK1/2 phosphorylation following stimulation with 100 nM GIP (1-42) was determined in HEK-293S cells or HEK-293AΔβ-arrestin cells transfected with GIPR. Data are expressed relative to 100 μM PMA and are the mean ± SEM of at least 8 individual data sets with quantitative data displayed in (Table S6). Data were assessed for statistical differences, at *p*<0.05, using Student's t-test (*, *p* < 0.05) using Student's t-test.

**Supplementary Figure 14**

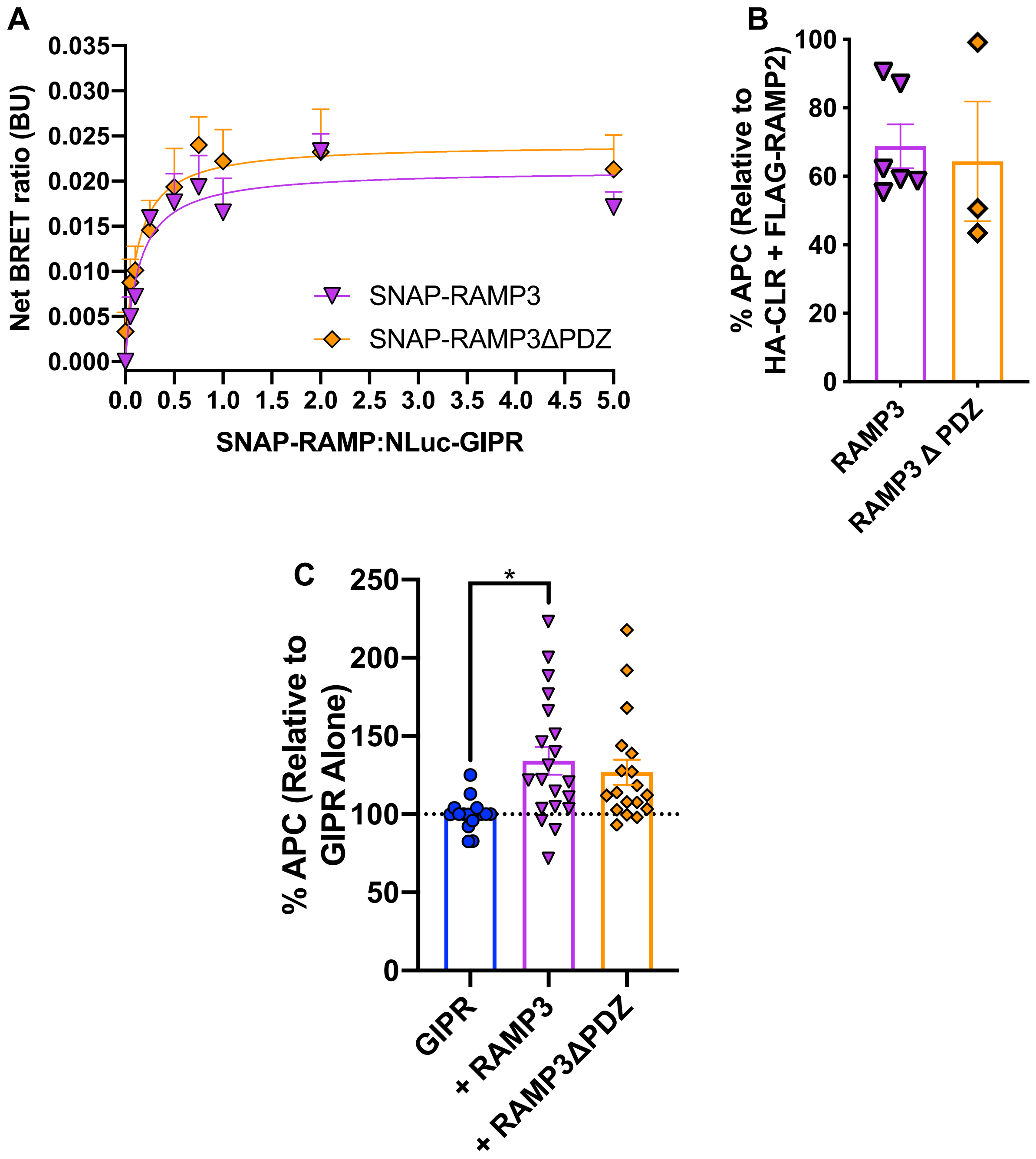


**Figure S14. GIPR promotes RAMP3ΔPDZ PM expression but RAMP3ΔPDZ has no significant effect on GIPR cell surface expression.**

*A*. HEK-293S cells were cotransfected with a constant concentration of Nluc-GIPR and increasing amounts of either SNAP-RAMP3 or SNAP-RAMP3ΔPDZ. ΔBRET was fitted to one site binding saturation curves using GraphPad Prism 8.4.2. *B-C*. HEK-293S cells were cotransfected with GIPR and either FLAG-RAMP3 or FLAG-RAMP3ΔPDZ (for RAMP PM expression, *B*) or FLAG-GIPR and either HA-RAMP3 or HA-RAMP3ΔPDZ (for GIPR PM expression, *C*). PM expression of FLAG-RAMPs or FLAG-GIPR was determined by flow cytometry. PM expression was normalised to that of FLAG-RAMP2 when cotransfected with HA-CLR as 100 % (for RAMP PM expression) or FLAG-GIPR alone (for GIPR PM expression) and pcDNA3.1-zeo as 0%. All values are the mean ± S.E.M of at least 3-11 individual data sets. Data were assessed for statistical differences, at $p<0.05$, in cell surface FLAG-RAMP expression compared to expression in the absence of receptor using a one-way ANOVA with Dunnett's post-hoc test (*, $p < 0.05$).

### Supplementary Figure 15

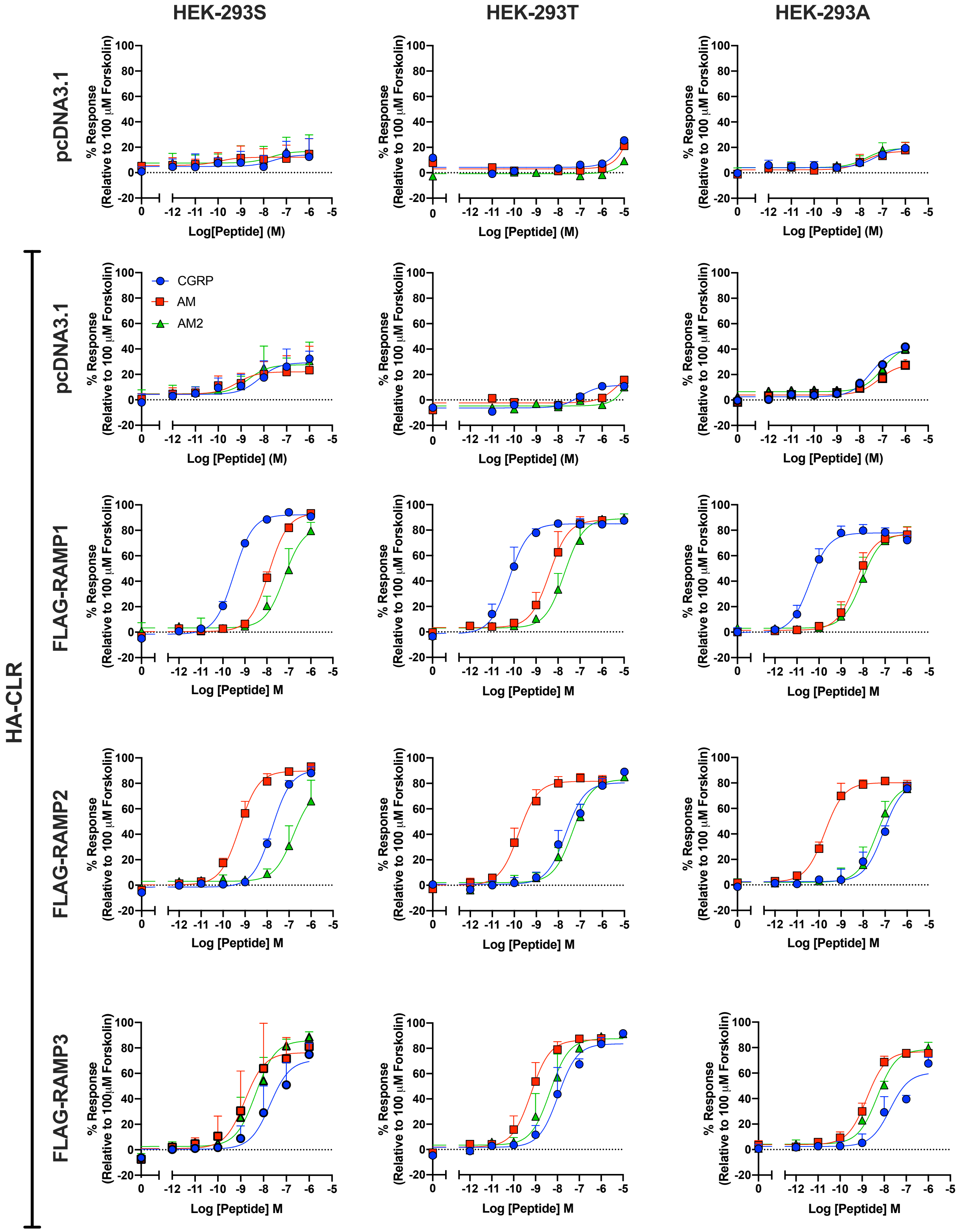


**Figure S15. Comparison of cAMP signalling between the three parental HEK-293 cell lines used in this study.**
cAMP accumulation was determined in HEK-293S, HEK-293T or HEK-293A cells transfected with HA-CLR and FLAG-RAMPs following 30 min stimulation with CGRP, AM or AM2. Data are expressed relative to 100 μM forskolin and are the mean ± SD of 2-4 individual data sets.

Table S1

| IUPHAR receptor name | N | Receptor Expression (RLU) +/- SEM | BRET50 (RAMP1) | Bmax (RAMP1) | BRET50 (RAMP2) | Bmax (RAMP2) | BRET50 (RAMP3) | Bmax (RAMP3) |
|---|---|---|---|---|---|---|---|---|
| CLR | 3 | 401536 +/- 58253 | 1.547 +/- 0.932 | 0.449 +/- 0.030 | 1.803 +/- 0.313 | 0.872 +/- 0.163 | 1.875 +/- 1.335 | 1.618 +/- 0.533 |
| $\beta_2$ADR | 3 | 359986 +/- 34576 | - | <0.1* | - | <0.1* | - | <0.1* |
| CTR | 3 | 557831 +/- 34371 | 1.019 +/- 0.100 | 0.439 +/- 0.132 | 2.121 +/- 0.353 | 0.772 +/- 0.039 | 1.187 +/- 0.160 | 0.883 +/- 0.040 |
| PTH1R | 3 | 272805 +/- 57337 | - | <0.1* | 0.903 +/- 0.122 | 0.174 +/- 0.046 | 1.597 +/- 0.163 | 0.300 +/- 0.060 |
| PTH2R | 3 | 178518 +/- 45350 | 4.598 +/- 3.056 | 0.498 +/- 0.066 | 8.904 +/- 1.030 | 0.509 +/- 0.010 | 7.109 +/- 0.932 | 0.947 +/- 0.244 |
| GCGR | 3 | 277561 +/- 63162 | 0.898 +/- 0.142 | 0.935 +/- 0.008 | 0.996 +/- 0.097 | 0.570 +/- 0.020 | 16.02 +/- 5.915 | 2.218 +/- 0.624 |
| GIPR | 3 | 164648 +/- 24338 | 1.583 +/- 0.636 | 0.801 +/- 0.079 | 0.963 +/- 0.122 | 0.671 +/- 0.068 | 6.617 +/- 3.216 | 1.681 +/- 0.065 |
| GLP1R | 3 | 247464 +/- 69032 | 0.576 +/- 0.420 | 0.430 +/- 0.056 | 1.322 +/- 0.285 | 1.025 +/- 0.067 | 4.042 +/- 1.661 | 1.076 +/- 0.078 |
| GLP2R | 3 | 328165 +/- 30034 | 2.823 +/- 0.616 | 1.099 +/- 0.158 | 2.229 +/- 0.315 | 0.796 +/- 0.071 | 6.328 +/- 0.018 | 1.343 +/- 0.012 |
| PAC1R | 3 | 87878 +/- 7802 | - | <0.1* | 2.980 +/- 1.491 | 0.185 +/- 0.061 | 6.172 +/- 5.218 | 0.273 +/- 0.068 |
| GHRHR | 3 | 42536 +/- 6683 | 13.88 +/- 3.193 | 0.908 +/- 0.151 | 24.05 +/- 3.840 | 1.308 +/- 0.371 | 26.08 +/- 2.855 | 1.631 +/- 0.242 |
| SCTR | 3 | 352128 +/- 46891 | 0.026 +/- 0.004 | 0.271 +/- 0.015 | 3.485 +/- 1.220 | 0.638 +/- 0.163 | 2.745 +/- 0.650 | 0.976 +/- 0.141 |
| CRF1R | 3 | 107131 +/- 9656 | - | <0.1* | 2.066 +/- 0.798 | 0.362 +/- 0.047 | 3.722 +/- 0.331 | 0.533 +/- 0.008 |
| CRF2R | 3 | 204995 +/- 15113 | LF | LF | 1.635 +/- 0.105 | 0.113 +/- 0.050 | 3.123 +/- 0.897 | 0.347 +/- 0.049 |
| VPAC1R | 3 | 63817 +/- 6438 | 19.43 +/- 13.99 | 0.664 +/- 0.130 | 24.27 +/- 14.45 | 0.455 +/- 0.071 | 35.83 +/- 18.04 | 1.250 +/- 0.193 |
| VPAC2R | 3 | 502196 +/- 21720 | 0.558 +/- 0.087 | 0.437 +/- 0.007 | 0.966 +/- 0.194 | 0.574 +/- 0.068 | 1.666 +/- 0.199 | 0.546 +/- 0.030 |

RLU: Relative Fluorescence Units, LF: Linear Fit, SEM: Standard Error of the Mean
* Denotes that the maximum measured BRET value was <0.1; "-" indicates no detectable BRET50

**Table S2**: Potency ($pEC_{50}$) values for G protein activation, following stimulation with GIP (1-42), GIP (D-Ala2) or GIP (Pro3) in HEK 293A cells expressing GIPR.

| | | $G_s$ | $G_{i2}$ | $G_q$ | $G_{12}$ |
|---|---|---|---|---|---|
| **GIP (1-42)** | **$pEC_{50}$[a]** | 10.14±0.09 | 9.52±0.17 | 9.16±0.13 | 9.96±0.16 |
| | **$E_{max}$[b]** | 98.7±2.8 | 100.1±4.9 | 100.1±4.2 | 98.4±4.6 |
| | **$pEC_{50\_1}$[c]** | ND | ND | ND | 11.00±0.22 |
| | **$pEC_{50\_2}$[d]** | ND | ND | ND | 8.18±0.24 |
| | **Frac[e]** | ND | ND | ND | 52.7±6 |
| | **n** | 7 | 7 | 6 | 7 |
| **GIP (D-Ala2)** | **$pEC_{50}$[a]** | 9.92±0.24 | 10.19±0.33 | 9.26±0.35 | 10.28±0.26 |
| | **$E_{max}$[b]** | 102.6±8.5 | 100.5±8.5 | 99.7±11.0 | 96.4±7.1 |
| | **$pEC_{50\_1}$[c]** | ND | ND | 9.77±0.43 | 10.97±0.30 |
| | **$pEC_{50\_2}$[d]** | ND | ND | 6.45±0.40 | 7.42±0.30 |
| | **Frac[e]** | ND | ND | 45.3±9 | 52.0±6 |
| | **n** | 5 | 5 | 5 | 4 |
| **GIP (Pro3)** | **$pEC_{50}$[a]** | 7.02±0.22 | 7.71±0.64 | 7.56±0.38 | 8.09±0.27 |
| | **$E_{max}$[b]** | 86.5±11.0 | 16.8±5.3 | 54.5±9.8 | 69.5±9.2 |
| | **n** | 5 | 4 | 5 | 4 |

Data are the mean ± SEM of (*n*) individual data sets. Data points were corrected to baseline and vehicle and the AUC after agonist addition used to produce concentration-response curves for each G protein. Values were obtained by fitting to the three-parameter logistic model.

[a] The negative logarithm of the agonist concentration required to produce a half-maximal response.

[b] The maximal response to the ligand expressed as a percentage of the response observed for GIP (1-42), determined by fitting to the three-parameter Log [Agonist] vs response model for each individual experiment.

[c] The negative logarithm of the agonist concentration required to produce a 50% of the first component of the response, determined by fitting to the biphasic Log [Agonist] vs response model

[d] The negative logarithm of the agonist concentration required to produce a 50% of the second component of the response, determined by fitting to the biphasic Log [Agonist] vs response model

[e] The fraction of the concentration-response curve derived from the first, high potency component

**Table S3**: Potency ($pEC_{50}$) values for G protein activation, following stimulation with GIP (1-42), in HEK 293A cells expressing GIPR and either pcDNA3.1, FLAG-RAMP1, FLAG-RAMP2 or FLAG-RAMP3.

| | | $G_s$ | $G_{i2}$ | $G_{i3}$ | $G_z$ | $G_q$ | $G_{11}$ | $G_{14}$ | $G_{15}$ | $G_{12}$ | $G_{13}$ |
|---|---|---|---|---|---|---|---|---|---|---|---|
| **GIPR** | $pEC_{50}$[a] | 10.14±0.09 | 9.52±0.17 | 7.88±0.15 | 8.63±0.11 | 9.16±0.13 | 7.67±0.17 | 6.91±0.42 | 7.78±0.14 | 9.96±0.16 | 9.07±0.15 |
| | $pEC_{50\_1}$[b] | ND | ND | ND | 8.77±0.55 | ND | ND | ND | ND | 11.00±0.22 | ND |
| | $pEC_{50\_2}$[c] | ND | ND | ND | 6.18±0.61 | ND | ND | ND | ND | 8.18±0.24 | ND |
| | Frac[d] | ND | ND | ND | 76.1±22 | ND | ND | ND | ND | 52.7±6 | ND |
| | n | 7 | 7 | 5 | 6 | 6 | 5 | 6 | 4 | 7 | 6 |
| **+ RAMP1** | $pEC_{50}$[a] | 10.15±0.16 | 9.94±0.13 | 8.65±0.22 | 8.44±0.16 | 8.08±0.18*** | 6.70±0.23* | 6.55±0.79 | 6.27±0.46* | 10.36±0.17 | 9.41±0.24 |
| | $pEC_{50\_1}$[b] | ND | ND | ND | 9.35±0.46 | ND | ND | ND | ND | 10.88±0.26 | ND |
| | $pEC_{50\_2}$[c] | ND | ND | ND | 6.76±0.32 | ND | ND | ND | ND | 8.30±0.42 | ND |
| | Frac[d] | ND | ND | ND | 49.4±16 | ND | ND | ND | ND | 62.9±8 | ND |
| | n | 3 | 5 | 4 | 3 | 5 | 3 | 4 | 3 | 4 | 4 |
| **+ RAMP2** | $pEC_{50}$[a] | 9.81±0.09 | 10.15±0.23 | 8.36±0.29 | 8.48±0.17 | 8.54±0.15* | 6.52±0.22** | 6.43±1.37 | 6.37±0.30* | 10.20±0.19 | 9.52±0.39 |
| | $pEC_{50\_1}$[b] | ND | ND | ND | 9.27±0.35 | ND | ND | ND | ND | 10.48±0.26 | ND |
| | $pEC_{50\_2}$[c] | ND | ND | ND | 6.63±0.29 | ND | ND | ND | ND | 7.81±0.51 | ND |
| | Frac[d] | ND | ND | ND | 50.6±13 | ND | ND | ND | ND | 69.3±9 | ND |
| | n | 5 | 4 | 3 | 3 | 5 | 3 | 4 | 3 | 4 | 3 |
| **+ RAMP3** | $pEC_{50}$[a] | 9.46±0.09*** | 9.76±0.21 | 8.18±0.18 | 8.16±0.14 | 9.12±0.18 | 7.55±0.21 | 6.96±0.62 | 8.25±0.26 | 10.76±0.17 | 9.55±0.23 |
| | $pEC_{50\_1}$[b] | ND | ND | ND | 8.69±0.49 | ND | ND | ND | ND | 10.98±0.23 | ND |
| | $pEC_{50\_2}$[c] | ND | ND | ND | 6.29±0.45 | ND | ND | ND | ND | 7.59±0.66 | ND |
| | Frac[d] | ND | ND | ND | 61.1±20 | ND | ND | ND | ND | 75.0±7 | ND |
| | n | 5 | 5 | 4 | 3 | 5 | 3 | 4 | 3 | 3 | 4 |

Data are the mean ± SEM of *n* individual data sets. Data points were corrected to baseline and vehicle and the AUC after agonist addition used to produce concentration-response curves for each G protein. Data were normalised to the maximum response to GIP (1-42) for each condition.

[a] The negative logarithm of the agonist concentration required to produce a half-maximal response, determined by fitting to the three-parameter Log [Agonist] vs response.

[b] The negative logarithm of the agonist concentration required to produce a 50% of the first component of the response, determined by fitting to the biphasic Log [Agonist] vs response model

[c] The negative logarithm of the agonist concentration required to produce a 50% of the second component of the response, determined by fitting to the biphasic Log [Agonist] vs response model

[d] The fraction of the concentration-response curve derived from the first, high potency component

Data were assessed for statistical differences, at $p<0.05$, in the change in $pEC_{50}$ compared to GIPR expressed with pcDNA3.1 using a one-way ANOVA with Dunnett's post-hoc test (*, $p < 0.05$; **, $p < 0.01$; ***, $p < 0.001$).

**Table S4**: Potency ($pEC_{50}$) values for G protein activation, following stimulation with GIP (D-Ala2), in HEK 293A cells expressing GIPR and either pcDNA3.1, FLAG-RAMP1, FLAG-RAMP2 or FLAG-RAMP3.

| | | $G_s$ | $G_{i2}$ | $G_q$ | $G_{12}$ |
|---|---|---|---|---|---|
| **GIPR** | **$pEC_{50}$[a]** | 10.26±0.18 | 10.24±0.20 | 9.63±0.21 | 10.49±0.21 |
| | **$pEC_{50\_1}$[b]** | ND | ND | 9.89±0.25 | 10.91±0.24 |
| | **$pEC_{50\_2}$[c]** | ND | ND | 6.87±0.45 | 7.50±0.36 |
| | **Frac[d]** | ND | ND | 66.3±8 | 63.3±6 |
| | **n** | 4 | 5 | 5 | 6 |
| **+ RAMP1** | **$pEC_{50}$[a]** | 10.08±0.21 | 10.11±0.15 | 7.79±0.24**** | 10.75±0.27 |
| | **$pEC_{50\_1}$[b]** | ND | ND | 11.14±0.64 | 11.00±0.28 |
| | **$pEC_{50\_2}$[c]** | ND | ND | 7.32±0.21 | 7.22±0.48 |
| | **Frac[d]** | ND | ND | 28.8±7** | 65.8±7 |
| | **n** | 3 | 5 | 5 | 4 |
| **+ RAMP2** | **$pEC_{50}$[a]** | 10.04±0.15 | 10.37±0.26 | 7.87±0.20**** | 10.45±0.26 |
| | **$pEC_{50\_1}$[b]** | ND | ND | 10.67±0.63 | 10.96±0.29 |
| | **$pEC_{50\_2}$[c]** | ND | ND | 7.23±0.19 | 7.42±0.39 |
| | **Frac[d]** | ND | ND | 27.0±8** | 59.5±7 |
| | **n** | 5 | 3 | 5 | 6 |
| **+ RAMP3** | **$pEC_{50}$[a]** | 9.20±0.15** | 10.21±0.26 | 9.53±0.20 | 10.97±0.22 |
| | **$pEC_{50\_1}$[b]** | ND | ND | 9.97±0.25 | 11.21±0.23 |
| | **$pEC_{50\_2}$[c]** | ND | ND | 7.21±0.34 | 7.41±0.46 |
| | **Frac[d]** | ND | ND | 60.7±7 | 69.5±6 |
| | **n** | 3 | 3 | 5 | 4 |

Data are the mean ± SEM of *n* individual data sets. Data points were corrected to baseline and vehicle and the AUC after agonist addition used to produce concentration-response curves for each G protein. Data were normalised to the maximum response to GIP (D-Ala2) for each condition.

[a] The negative logarithm of the agonist concentration required to produce a half-maximal response, determined by fitting to the three-parameter Log [Agonist] vs response.

[b] The negative logarithm of the agonist concentration required to produce a 50% of the first component of the response, determined by fitting to the biphasic Log [Agonist] vs response model

[c] The negative logarithm of the agonist concentration required to produce a 50% of the second component of the response, determined by fitting to the biphasic Log [Agonist] vs response model

[d] The fraction of the concentration-response curve derived from the first, high potency component

Data were assessed for statistical differences, at $p<0.05$, in the change in $pEC_{50}$ compared to GIPR expressed with pcDNA3.1 using a one-way ANOVA with Dunnett's post-hoc test (**, $p < 0.01$; ****, $p < 0.0001$).

**Table S5**: Potency ($pEC_{50}$) values for G protein activation, following stimulation with GIP (Pro3), in HEK 293A cells expressing GIPR and either pcDNA3.1, FLAG-RAMP1, FLAG-RAMP2 or FLAG-RAMP3.

| | | $G_s$ | $G_{i2}$ | $G_q$ | $G_{12}$ |
|---|---|---|---|---|---|
| **GIPR** | **$pEC_{50}$[a]** | 7.06±0.13 | 7.07±0.42 | 6.79±0.26 | 7.55±0.29 |
| | **$pEC_{50\_1}$[b]** | ND | ND | ND | 8.18±0.61 |
| | **$pEC_{50\_2}$[c]** | ND | ND | ND | 6.25±0.35 |
| | **Frac[d]** | ND | ND | ND | 47.1±29 |
| | **n** | 4 | 4 | 4 | 4 |
| **+ RAMP1** | **$pEC_{50}$[a]** | 6.87±0.13 | 6.78±0.43 | ND | 6.72±0.15 |
| | **$pEC_{50\_1}$[b]** | ND | ND | ND | 8.08±0.31 |
| | **$pEC_{50\_2}$[c]** | ND | ND | ND | 6.27±0.10 |
| | **Frac[d]** | ND | ND | ND | 37.9±12 |
| | **n** | 3 | 4 | 5 | 4 |
| **+ RAMP2** | **$pEC_{50}$[a]** | 7.18±0.20 | 7.04±0.24 | ND | 6.83±0.25 |
| | **$pEC_{50\_1}$[b]** | ND | ND | ND | 8.71±0.42 |
| | **$pEC_{50\_2}$[c]** | ND | ND | ND | 6.42±0.19 |
| | **Frac[d]** | ND | ND | ND | 26.5±13 |
| | **n** | 4 | 4 | 4 | 4 |
| **+ RAMP3** | **$pEC_{50}$[a]** | 6.14±0.23* | 6.96±0.36 | 6.31±0.36 | 7.29±0.15 |
| | **$pEC_{50\_1}$[b]** | ND | ND | ND | 7.90±0.39 |
| | **$pEC_{50\_2}$[c]** | ND | ND | ND | 6.10±0.16 |
| | **Frac[d]** | ND | ND | ND | 55.5±17 |
| | **n** | 4 | 4 | 5 | 4 |

Data are the mean ± SEM of *n* individual data sets. Data points were corrected to baseline and vehicle and the AUC after agonist addition used to produce concentration-response curves for each G protein. Data were normalised to the maximum response to GIP (Pro3) for each condition.

[a] The negative logarithm of the agonist concentration required to produce a half-maximal response, determined by fitting to the three-parameter Log [Agonist] vs response.

[b] The negative logarithm of the agonist concentration required to produce a 50% of the first component of the response, determined by fitting to the biphasic Log [Agonist] vs response model

[c] The negative logarithm of the agonist concentration required to produce a 50% of the second component of the response, determined by fitting to the biphasic Log [Agonist] vs response model

[d] The fraction of the concentration-response curve derived from the first, high potency component

Data were assessed for statistical differences, at $p<0.05$, in the change in $pEC_{50}$ compared to GIPR expressed with pcDNA3.1 using a one-way ANOVA with Dunnett's post-hoc test (*, $p < 0.05$).

**Table S6**: ERK1/2 phosphorylation, following 0, 1, 2, 4, 5, 6, 8, 10, 15, 20, 30, 45, 60, 75, 90, 105 and 120 minutes stimulation with GIP (1-42), in either HEK-293S cells or HEK-293AΔβ-arrestin cells expressing GIPR and pcDNA3.1.

| | HEK 293S | | HEK 293AΔβ-arrestin | |
|---|---|---|---|---|
| **Time** | **Response** | **n** | **Response** | **n** |
| **0** | -0.1±2.6 | 8 | -0.0±1.0 | 10 |
| **1** | 19.0±4.5 | 9 | 25.4±3.3 | 10 |
| **2** | 22.8±3.3 | 10 | 24.8±2.9 | 9 |
| **4** | 26.6±3.7 | 10 | 30.3±2.5 | 10 |
| **5** | 26.1±2.8 | 10 | 33.5±3.2 | 10 |
| **6** | 28.7±3.9 | 10 | 34.8±3.6 | 9 |
| **8** | 33.7±4.9 | 9 | 36.0±4.3 | 9 |
| **10** | 38.8±4.0 | 9 | 40.0±3.8 | 9 |
| **15** | 25.7±3.3 | 9 | 29.1±3.0 | 10 |
| **20** | 26.1±2.3 | 8 | 19.8±2.5 | 10 |
| **30** | 14.4±3.9 | 9 | 12.5±3.3 | 10 |
| **45** | 3.5±4.4 | 10 | 7.0±3.0 | 9 |
| **60** | 11.6±3.0 | 8 | 3.4±2.3* | 10 |
| **75** | 11.5±2.3 | 8 | 5.4±1.6* | 8 |
| **90** | 11.3±5.2 | 9 | 1.2±1.8 | 10 |
| **105** | 4.0±3.1 | 10 | 1.1±1.8 | 10 |
| **120** | 2.1±3.0 | 10 | 0.8±2.9 | 8 |

Data are the mean ± SEM of n individual data sets.

Data were normalised to 100 μM PMA.

Data were assessed for statistical differences, at *p*<0.05, in the change in $pEC_{50}$ compared to GIPR expressed with pcDNA3.1 using Student's t-test (*, *p*<0.05).

**Table S7**: ERK1/2 phosphorylation, following 0, 1, 2, 4, 5, 6, 8, 10, 15, 20, 30, 45, 60, 75, 90, 105 and 120 minutes stimulation with GIP (1-42), in HEK 293S cells expressing GIPR and either pcDNA3.1, FLAG-RAMP1, FLAG-RAMP2 or FLAG-RAMP3.

| Time | + pcDNA3.1-zeo | + FLAG-RAMP1 | + FLAG-RAMP2 | +FLAG-RAMP3 |
|---|---|---|---|---|
| **0** | -0.1±2.6 (*8*) | 0.8±2.0 (*8*) | 0.5±2.2 (*6*) | 3.9±4.8 (*7*) |
| **1** | 19.0±4.5 (*9*) | 3.6±1.6$^{*}$ (*11*) | 4.5±2.0 (*6*) | 29.7±5.0 (*12*) |
| **2** | 22.8±3.3 (*10*) | 7.5±2.3$^{*}$ (*11*) | 2.7±1.6$^{**}$ (*6*) | 27.1±4.7 (*12*) |
| **4** | 26.6±3.7 (*10*) | 11.3±1.7$^{*}$ (*11*) | 6.1±3.4$^{*}$ (*5*) | 33.6±5.4 (*12*) |
| **5** | 26.1±2.8 (*10*) | 16.0±3.1 (*11*) | 2.9±1.6$^{**}$ (*6*) | 34.1±5.5 (*11*) |
| **6** | 28.7±3.9 (*10*) | 16.8±3.0 (*11*) | 11.3±4.0$^{*}$ (*6*) | 35.2±5.6 (*12*) |
| **8** | 33.7±4.9 (*9*) | 22.6±3.4 (*11*) | 10.6±2.6$^{**}$ (*6*) | 39.0±4.6 (*12*) |
| **10** | 38.8±4.0 (*9*) | 19.4±3.6$^{**}$ (*11*) | 14.7±3.1$^{**}$ (*5*) | 41.3±3.9 (*12*) |
| **15** | 25.7±3.3 (*9*) | 20.3±2.4 (*10*) | 22.4±2.6 (*4*) | 42.7±6.1$^{*}$ (*12*) |
| **20** | 26.1±2.3 (*8*) | 13.8±3.1$^{*}$ (*11*) | 22.7±2.6 (*6*) | 39.0±2.9$^{**}$ (*12*) |
| **30** | 14.4±3.9 (*9*) | 6.7±2.3 (*11*) | 16.2±3.7 (*5*) | 34.8±3.2$^{***}$ (*10*) |
| **45** | 3.5±4.4 (*10*) | 3.7±2.2 (*10*) | 4.1±5.1 (*5*) | 23.5±2.3$^{***}$ (*10*) |
| **60** | 11.6±3.0 (*8*) | 4.1±1.9 (*10*) | -2.1±3.7$^{*}$ (*6*) | 24.2±5.5$^{*}$ (*7*) |
| **75** | 11.5±2.3 (*8*) | 6.4±2.6 (*9*) | -1.9±4.9 (*6*) | 24.2±5.5$^{*}$ (*7*) |
| **90** | 11.3±5.2 (*9*) | 5.6±2.7 (*11*) | -3.1±3.9 (*6*) | 14.7±4.9 (*11*) |
| **105** | 4.0±3.1 (*10*) | 3.3±2.3 (*11*) | -2.8±2.1 (*6*) | 12.2±4.8 (*12*) |
| **120** | 2.1±3.0 (*10*) | -3.5±3.7 (*11*) | -2.6±2.2 (*6*) | 12.7±4.6 (*12*) |

Data are the mean ± SEM of (*n*) individual data sets.
Data were normalised to 100 μM PMA.
Data were assessed for statistical differences, at $p<0.05$, in the change in $pEC_{50}$ compared to GIPR expressed with pcDNA3.1 using Student's t-test (*, $p < 0.05$; **, $p < 0.01$; ***, $p < 0.001$).

**Table S8**: ERK1/2 phosphorylation, following 0, 1, 2, 4, 5, 6, 8, 10, 15, 20, 30, 45, 60, 75, 90, 105 and 120 minutes stimulation with GIP (1-42), in HEK 293S cells expressing GIPR and either pcDNA3.1, or FLAG-RAMP3ΔPDZ.

| | GIPR | | + FLAG-RAMP3ΔPDZ | |
|---|---|---|---|---|
| **Time** | **Response** | **n** | **Response** | **n** |
| **0** | -0.1±2.6 | 8 | 4.5±4.8 | 5 |
| **1** | 19.0±4.5 | 9 | 8.7±2.5 | 8 |
| **2** | 22.8±3.3 | 10 | 15.9±2.0 | 8 |
| **4** | 26.6±3.7 | 10 | 17.0±1.9 | 7 |
| **5** | 26.1±2.8 | 10 | 20.5±3.9 | 8 |
| **6** | 28.7±3.9 | 10 | 19.9±3.2 | 8 |
| **8** | 33.7±4.9 | 9 | 25.5±2.7 | 8 |
| **10** | 38.8±4.0 | 9 | 30.9±2.7 | 8 |
| **15** | 25.7±3.3 | 9 | 25.3±3.6 | 8 |
| **20** | 26.1±2.3 | 8 | 14.5±4.3* | 8 |
| **30** | 14.4±3.9 | 9 | 7.8±4.1 | 8 |
| **45** | 3.5±4.4 | 10 | 8.1±4.5 | 7 |
| **60** | 11.6±3.0 | 8 | 16.6±7.6 | 7 |
| **75** | 11.5±2.3 | 8 | 20.4±4.4 | 7 |
| **90** | 11.3±5.2 | 9 | 23.3±5.2 | 7 |
| **105** | 4.0±3.1 | 10 | 15.6±4.8* | 7 |
| **120** | 2.1±3.0 | 10 | 11.7±2.8* | 8 |

Data are the mean ± SEM of n individual data sets.
Data were normalised to 100 μM PMA.
Data were assessed for statistical differences, at $p<0.05$, in the change in $pEC_{50}$ compared to GIPR expressed with pcDNA3.1 using Student's t-test (*, $p<0.05$).

**Table S9**: Potency ($pEC_{50}$) and $E_{max}$ values for β-arrestin-1 and β-arrestin-2 recruitment, following stimulation with GIP (1-42), in HEK 293T cells expressing GIPR and either pcDNA3.1, FLAG-RAMP3 or FLAG-RAMP3ΔPDZ.

| | | 6 Min | | 60 Min | |
|---|---|---|---|---|---|
| | | **GIPR** | **+ RAMP3ΔPDZ** | **GIPR** | **+ RAMP3 ΔPDZ** |
| **β-arrestin-1** | **$pEC_{50}$[a]** | 8.17±0.38 | 8.45±0.48 | 7.99±0.40 | 7.77±0.16 |
| | **$E_{max}$[b]** | 2.53±0.33 | 2.98±1.43 | 3.72±1.15 | 2.80±0.86 |
| | **n** | 4 | 4 | 4 | 4 |
| **β-arrestin-2** | **$pEC_{50}$[a]** | 8.37±0.36 | 8.34±0.23 | 9.05±0.24 | 8.64±0.46 |
| | **$E_{max}$[b]** | 8.3±0.96 | 8.53±1.25 | 6.42±1.12 | 7.99±1.41 |
| | **n** | 4 | 4 | 4 | 4 |

Data are the mean ± SEM of *n* individual data sets.

[a] The negative logarithm of the agonist concentration required to produce a half-maximal response.

[b] The maximal response to GIP (1-42) minus the baseline in mBRET units.
